# Benchmarking large-scale single-cell RNA-seq analysis

**DOI:** 10.1101/2025.10.28.681564

**Authors:** Ilaria Billato, Herve Pages, Vince Carey, Levi Waldron, Gabriele Sales, Chiara Romualdi, Davide Risso

## Abstract

The increasing size of single-cell RNA sequencing (scRNA-seq) datasets poses major computational challenges. This work benchmarks the scalability, efficiency, and accuracy of five widely used analysis frameworks (Seurat, OSCA, scrapper, Scanpy, and rapids singlecell), focusing on the impact of algorithmic and infrastructural choices on performance. We performed a systematic comparison of these workflows using representative datasets, including a 1.3 million and a 4.2 million cell dataset for scalability and four smaller datasets (BE1, sc_mixology, cord blood CITE-seq and Hao) with ground truth labels to assess clustering accuracy. Principal Component Analysis (PCA) was used as a paradigmatic step to evaluate the computational performance of six SVD algorithms (exact, ARPACK, IRLBA, randomized, Jacobi, and incremental PCA) across multiple data representations (dense, sparse, HDF5) and hardware configurations (CPU vs GPU). All methods showed high concordance in PCA results, with negligible loss of accuracy in truncated approaches. GPU-based computation using rapids singlecell provided a **15*×*** speed-up over the best CPU methods, with moderate memory usage. On CPU, ARPACK and IRLBA were the most efficient for sparse matrices, while randomized SVD performed best for HDF5-backed data. Among full pipelines, rapids singlecell was the fastest, whereas OSCA and scrapper achieved the highest clustering accuracy (ARI up to 0.97) in datasets with known cell identities. Performance differences were largely driven by the choice of highly variable genes (HVGs) and PCA implementation. The study highlights that scalability in scRNA-seq analysis depends critically on both algorithmic and infrastructural factors. GPU acceleration and optimized BLAS/LAPACK configurations markedly enhance performance, while Bioconductor-based pipelines remain robust in accuracy. The provided benchmarks offer practical guidelines for efficient and reliable analysis of large-scale single-cell datasets.

## 1 Introduction

As single-cell RNA sequencing (scRNA-seq) reaches its teenage years [1], we are witnessing a rapid increase in the size and complexity of experiments and datasets [2]. In fact, whereas early scRNA-seq comprised hundreds to few thousands cells, typically in a single condition and often without biological replicates [3], contemporary experiments include cells from multiple individuals measured across conditions (e.g., treatments, genotypes, health states). Often, researchers target 5 − 10,000 cells per replicate and the final dataset easily comprises hundreds of thousands of cells (e.g., [4, 5]). Furthermore, single-cell atlases have become mature and thanks to programmatic access, it is now straightforward to download and locally analyze datasets made of millions of cells from multiple organs across different labs [6, 7].

This increasing complexity poses significant computational challenges throughout the analysis pipeline, from data preprocessing to downstream interpretation. These challenges are exacerbated by the exploratory nature of many scRNA-seq analyses, which need to process large datasets in an interactive way, e.g., trying different analysis paths and exploring the downstream results, often using desktops or laptops rather than high-performance computing (HPC), requiring frugal and efficient workflows.

A typical scRNA-seq analysis workflow starts with gene expression quantitation, a process that includes assigning reads to barcodes, aligning reads to the appropriate genome or transcriptome, and quantifying gene expression by counting Unique Molecular Identifiers (UMIs) assigned to each gene. This process is typically performed in HPC clusters and is usually carried-out with stand-alone software or standardized pipelines. This step is typically not part of the interactive process described above; hence for the sake of this article, we will consider the output of this step as the starting point of the analysis. Interested readers can refer to the literature for a benchmark of the different preprocessing pipelines (e.g., [8]).

After preprocessing, the analysis workflow comprises the following steps (Fig. 1a): (i) quality control; (ii) gene and cell filtering; (iii) normalization; (iv) identification of highly variable genes; (v) dimensionality reduction, generally employed using principal component analysis (PCA) or similar methods; (vi) data visualization, using methods such as t-SNE and UMAP, which are commonly applied to data reduced through PCA; (vii) clustering for the identification of groups of cells with similar transcriptional profiles; and (viii) cell type annotation, the process by which cells or clusters are labeled using either external reference datasets or the expression of known marker genes [9].

**Fig. 1.**
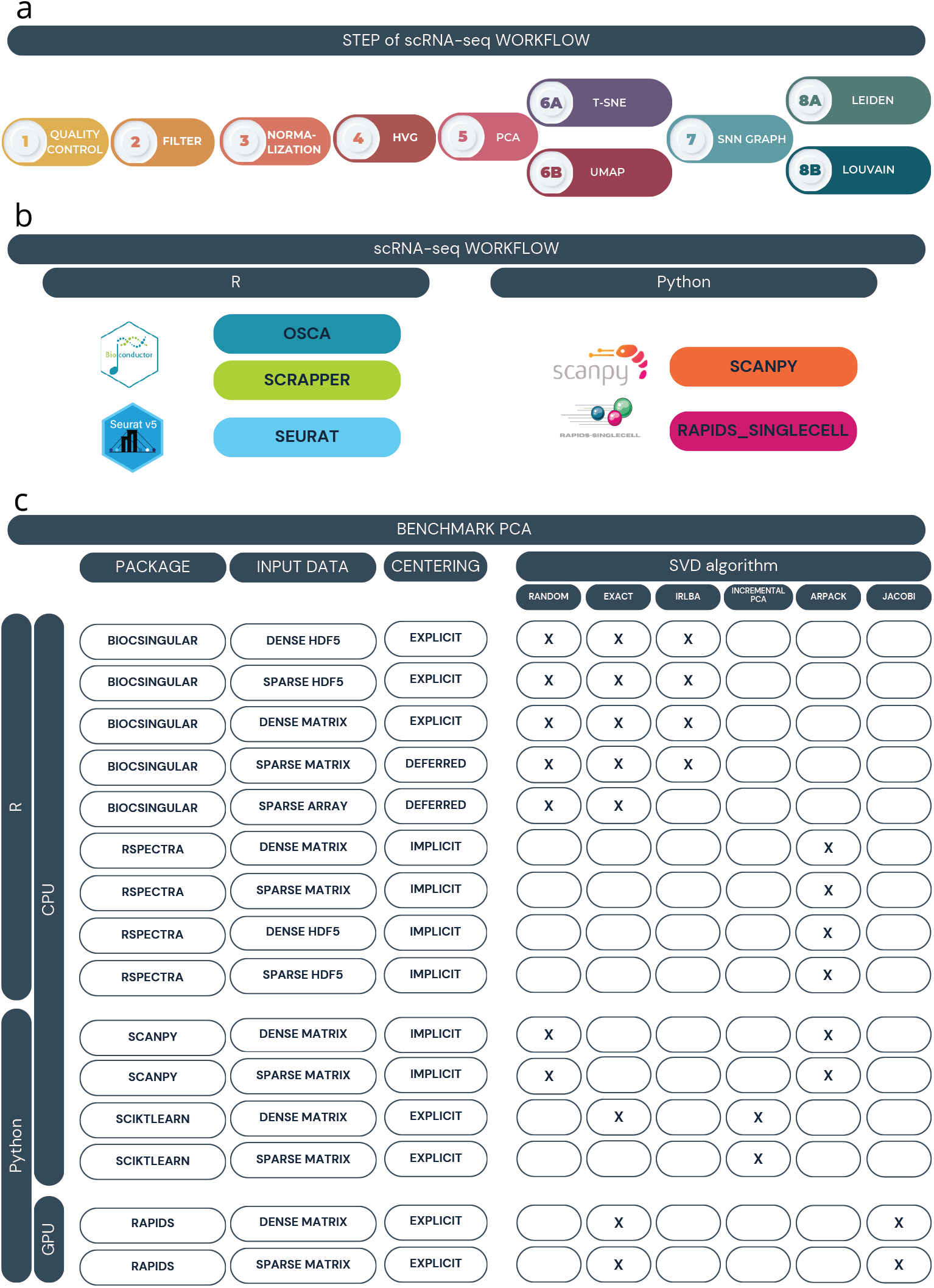
Overview of PCA benchmarking and single-cell workflow comparison. (a) Schematic of the typical steps in single-cell data processing workflows, including quality control, filtering, normalization, highly variable genes selection, dimensionality reduction (PCA, UMAP, t-SNE), SNN Graph and clustering (Louvain, Leiden). (b) List of R and Python-based single-cell RNA-seq analysis frameworks used for comparing full workflows: OSCA, Scrapper, and Seurat (R); Scanpy and rapids singlecell (Python).(c) Summary of PCA implementations evaluated in the benchmark, categorized by programming language (R or Python), computation type (CPU or GPU), input data format (dense matrix, sparse matrix, or HDF5), library name, and centering computation. The table also indicates the supported SVD algorithms for each method (random, exact, IRLBA, Incremental PCA, ARPACK, Jacobi).

Each of these steps requires its own processing time and memory usage and, in the last few years, several approaches have been proposed in different programming languages (mostly R and Python).

While there have been attempts in the literature to benchmark individual analysis steps, such as normalization, dimensionality reduction, clustering, trajectory inference, cell-type annotation, data integration [10–15], the comparison of entire analysis workflows has received less attention. Rich et al. [16] explores the impact of parameter choice and their default values in Seurat and Scanpy, but does not thoroughly examine their scalability. On the other hand, Tian et al. [17], while benchmarking all the different steps of a typical analysis, does not consider the workflow typically recommended by the most popular frameworks. Moreover, the dataset used for benchmarking is relatively small and insufficient to tackle scalability issues.

Here, we focus on the three most popular analysis frameworks, namely, Bioconductor [9], scverse/Scanpy [18], and Seurat [19] (Fig. 1b). For Bioconductor, we consider both the standard Orchestrating Single-Cell Analysis (OSCA) workflow and scrapper, a more recent efficient implementation that offloads most computations to C++ (see Methods). To compare these workflows, we rely on their respective documentation, purposely choosing the “quick start” or “basic” tutorials, which comprise the steps that are always present in one’s analysis; more advanced methods, such as multi-sample integration, batch-effect reduction, reference-based annotations, are not included in our benchmark.

While traditional approaches rely on CPU architectures, although often allowing for multithreading and parallel computing, recent efforts have explored the use of GPUs to enhance speed and scalability [20]. Hence, we include in our benchmark the rapids singlecell framework [21], an open-source library that implements a standard single-cell analysis workflow leveraging NVIDIA GPUs (Fig. 1b). In this context, benchmarking tools and workflows is essential for evaluating the performance of different methods in different experimental settings (such as the number of cells, sequencing depth, or biological complexity) not just in terms of efficiency, but to ensure that scalability does not come at the price of reduced accuracy.

The goal of this article is three-fold: first, we aim to benchmark the five abovementioned scRNA-seq workflows, namely Seurat, OSCA, scrapper, Scanpy, and rapids singlecell; second, by focusing on PCA as an exemplar analysis step, we explore in greater details how different algorithms and input choices can affect the efficiency and scalability of methods; last, we explore efficient workflows, in CPU and GPU, establishing guidelines and recommendations for the user in need of efficient analysis of large data.

The two main components of our benchmark are closely linked in practice, though, for clarity, we present them separately: Section 2.1 focuses on the choice of the SVD implementation as an example of the algorithmic decisions each workflow make internally. This PCA benchmark is designed to isolate and explain one specific source of the runtime and memory differences observed between workflows, which are compared in full in Section 2.2. There, we show that other important factors, such as the choice of highly variable genes (HVGs), differ between workflows and affect their accuracy, complementing the efficiency-focused PCA benchmark with an accuracy-focused comparison of complete analysis strategies.

## 2 Results

### 2.1 Benchmarking PCA

While quality control, normalization, and clustering are all essential steps in single-cell data analysis, we chose to focus on PCA as an exemplar step to thoroughly benchmark. In fact, PCA represents a key focal point of the entire pipeline as it is a non-trivial, often computationally heavy step, necessary for the subsequent stages of the analysis. Moreover, several algorithms exist to compute PCA and the choice of the algorithm can have profound impact on its computational load [15].

First, we give a brief introduction to PCA, reviewing standard methods for computing the principal components of a data matrix, i.e., the Singular Value Decomposition (SVD), as well as some of the most popular approximations, based on truncated SVD. Briefly, PCA is a multivariate technique that consists of defining a new coordinate system defined by a set of orthogonal vectors, which correspond to the direction of maximal variability in the data. The principal components (PCs) are hence a set of linear combination of the original variables (in our case the genes) that explains the most variation in the data.

PCA is typically used as a dimensionality reduction technique, in which the first few components are retained and the data are projected onto their subspace. PCA ensures that this is the “best” linear subspace, in the sense that it preserves the most variation in the data.

In mathematical terms, the principal components are the eigenvectors of the covariance matrix and can be computed by its eigendecomposition. Alternatively, the PCs can be computed directly by the SVD of the original expression matrix. We refer the reader to [22] and [23] for a more thorough introduction and the mathematical details. In practice, PCA implementations almost universally rely on SVD rather than on direct eigendecomposition of the covariance matrix. Forming the covariance (or Gram) matrix explicitly requires computing an *m* × *m* (or *n* × *n*) product, which squares the condition number of the original data matrix and can amplify rounding errors, particularly for the smaller singular values/PCs. SVD-based algorithms avoid this explicit product, operating instead directly on *X* – often through matrix-vector products alone – which is both more numerically stable and more compatible with the sparse and out-of-memory matrix representations considered throughout this benchmark [24].

SVD is a standard matrix factorization technique, and many algorithms exist for its computation (see e.g., [24]). However, when only the first few PCs are needed, approximated methods, collectively known as truncated or partial SVD, are computationally advantageous (see Methods for details).

When choosing the appropriate SVD algorithm, the input format may play an important role. In fact, it is likely that an algorithm that very efficiently computes the first *k* principal components when the data are stored in RAM memory, becomes highly inefficient when the data are stored out of RAM memory, as in the case of delayed operations on HDF5-backed datasets [25–27]. Similar considerations may be made for dense versus sparse matrix representations.

In our benchmark, we compared the computational time and RAM memory usage of six SVD algorithms, implemented in R and Python, applied to three different input formats for a total of 32 combinations (Fig. 1c and Supplementary Table S2). The process was applied to four subsets of the 1.3M cell dataset of increasing size and repeated ten times for both time and memory assessments, for a total of 2,240 SVD calculations.

In particular, the SVD algorithms considered include ARPACK [28], random [29], exact [30], IRLBA [31], Jacobi [32], and incrementalPCA [33], with technical details provided in the Methods section. In R, we used the BiocSingular [34] and RSpectra [35] packages, while in Python, we relied on Scanpy [18], scikit-learn [36], and RAPIDS [20] (Fig. 1c).

Several input data formats were considered: the data were stored either in an HDF5 [25] file or in RAM memory, and in each case they were stored as dense or sparse matrices. We compared these input formats with the *SparseArray* representation [37] (Fig. 1c). For HDF5-backed data, in addition to the dense and sparse (TENxMatrix) on-disk formats, we considered a third representation (`hdf5_sd`), in which the dense HDF5 dataset is read with HDF5Array(`…, as.sparse = TRUE`). In this case the data remain stored on disk in dense format, but each block is loaded into memory as a sparse *SparseArray* object, so that block-processed operations can exploit sparsity without requiring a sparse on-disk layout [38]. This exploration allowed us to discern how the choice of input data influenced the results. For sparse matrices, whenever possible, we employed implicit or deferred centering and scaling to leverage the sparsity of the matrix as efficiently as possible.

To evaluate the scalability, we considered three datasets (random subsamples of the 1.3M cell dataset [39]) with the same genes but with an increasing number of cells: 100k, 500k, 1M (see Methods for details). To ensure the robustness of the results, each method was executed 10 times on the same machine, using a single core, mitigating potential biases arising from parallel computation.

Indeed, certain algorithms are optimized for parallel processing, while others are not. In some cases, increasing the number of cores does not enhance computational performance (Supplementary Fig. S1b). For example, with the IRLBA algorithm, parallel processing results in a substantial increase in computation time, rising from 0.39 minutes with a single core to 22.56 minutes with two cores (Supplementary Fig. S1a). This is due to the software implementation, which switches from C to pure R code when the user requests more than one core.

Hereafter, the term “method”, which indicates the combination of algorithm and input type, is used as the comparison unit. The names of the methods follow this structure: package name + input data type + deferred (optional) + SVD algorithm.

#### 2.1.1 Accuracy of truncated SVD

Before analyzing the performance in terms of computational time and memory usage, we evaluate the degree of agreement among the different methods to ensure that the faster algorithms return a good approximation of the true principal components. We measured the consistency across methods by computing the correlation between principal components (PCs). Because PCA is rotation-invariant and the singular vectors are defined only up to sign changes, we report the absolute value of the correlation when comparing components, using the exact SVD algorithm as the ground truth.

In single-cell analysis, the number of selected PCs can vary substantially depending on the dataset and analytical requirements. However, a maximum of 50 PCs are often assumed to capture sufficient variability. Hence, we consider here only the first 50 PCs. Figure 2 shows the correlation between exact SVD (bioc dense exact) and all the other methods in the 1.3M cell dataset. Strikingly, the correlation between exact SVD and most other methods is exactly equal to 1 for all considered PCs. The only exception is random SVD, which, especially in its R implementation, shows a significant decline in correlation values (up to values close to 0) starting from the 35th PC. A notable difference in the implementations of random SVD is the number of default power iterations, which is higher in Python than in R, leading to a more accurate approximation. However, it is important to note that typically the variance explained by the 35th and later PCs is very low. For instance, in the 1.3M cell dataset, the PCs from 35 to 50 in bioc_dense_random explain only 0.2% of the total variability (Supplementary Tables S11–S15). Hence, this result is unlikely to have a large practical importance for real data analysis. Supplementary Figure S2 shows the same analysis on random subsets of 100k, 500k, 1M cells.

**Fig. 2.**
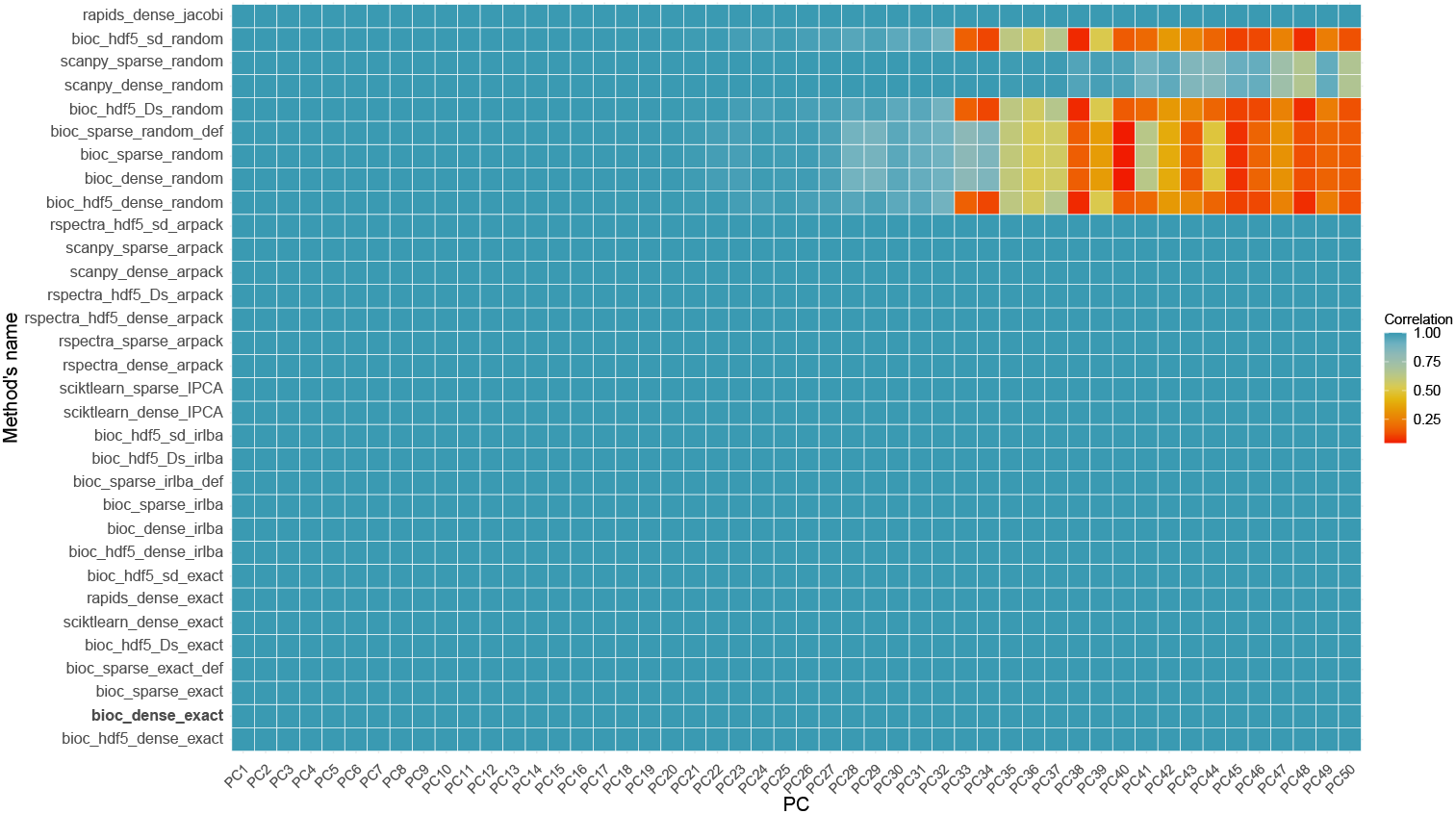
Correlation heatmap between principal components (PC) Correlation heatmap between principal components (PCs) obtained using different PCA algorithms and the Exact Algorithm as reference on 1.3M cell dataset. Each row represents a different combination of PCA algorithm and input matrix, and each column a principal component (PC1–PC50). The color indicates the absolute Pearson correlation between the PC obtained from each method and the corresponding PC from the exact reference.

#### 2.1.2 Computational time and memory usage

Figure 3 reports the time (right side) and memory consumption (left side) for each combination of SVD algorithm implementation, computational platform, and input type for the 1.3M cell dataset (see Supplementary Tables S10 and S6 for the exact numerical results).

**Fig. 3.**
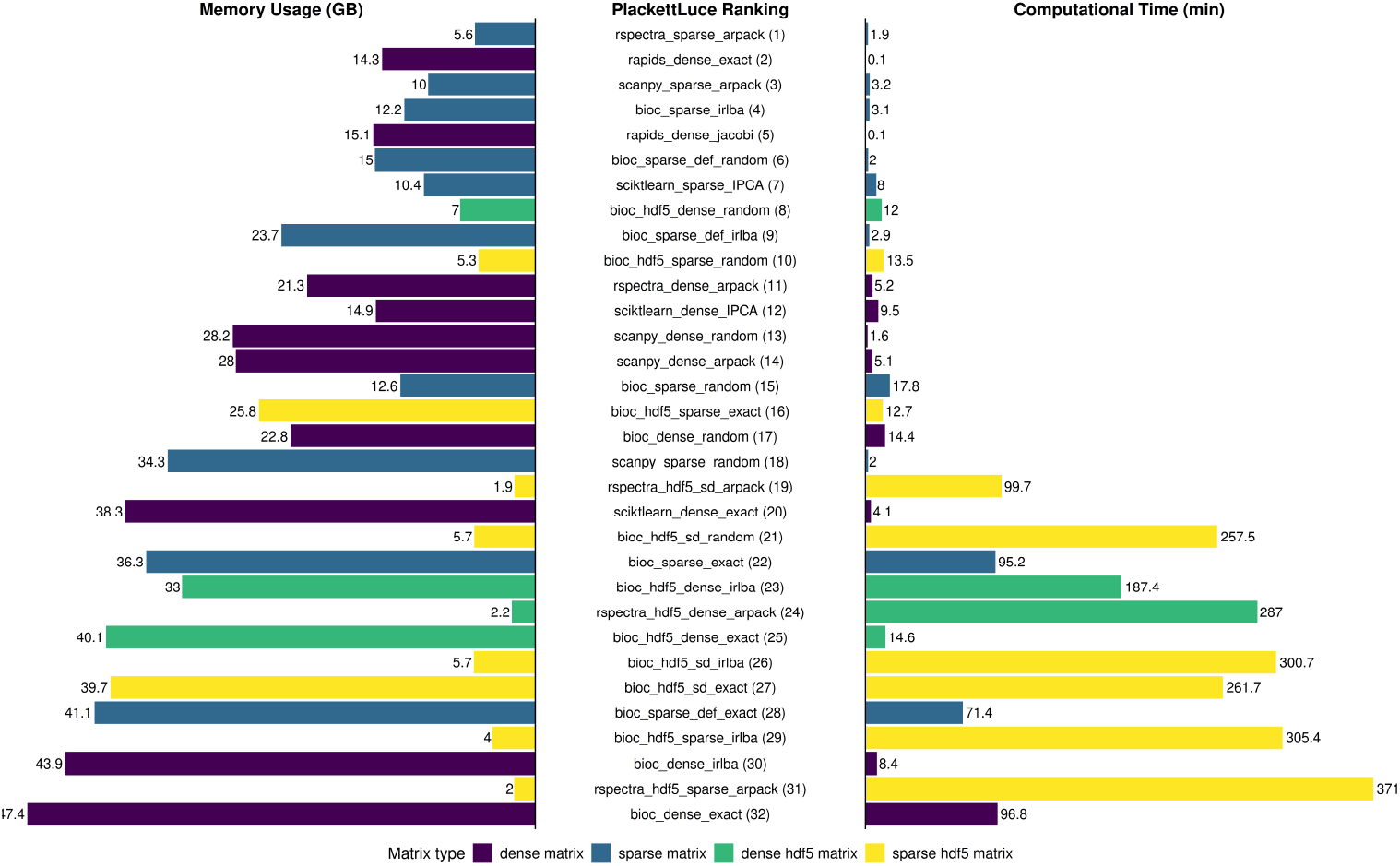
Comparison of PCA method performance across input matrix types for 1.3M cell dataset. Barplots show (left) peak memory usage (GB), (center) Plackett–Luce ranking (lower is better), and (right) computational time (minutes) for the 1.3M cell dataset. Each bar represents a specific combination of SVD algorithm and matrix format. Colors indicate the type of input matrix: dense matrix (purple), sparse matrix (blue), dense HDF5 matrix (green), and sparse HDF5 matrix (yellow), the latter including both the sparse on-disk (TENxMatrix) format and the dense HDF5 dataset read as sparse (`hdf5_sd`).

The GPU-aware RAPIDS library is the fastest approach, independent of the SVD algorithm, being at least one order of magnitude faster than CPU methods. Its memory footprint is moderate, even though it works only on dense matrices, providing an excellent choice for those that have a GPU available. In fact, it only takes 7.5 seconds to compute the exact SVD of a matrix with 1.3 million cells (Fig. 3, Supplementary Table S6).

The performance of exact SVD using CPU is surprisingly different in stock installations of Python and R. Indeed, Python’s scikit-learn implementation of exact SVD is 20 times faster than R’s base *svd* function (Fig. 3, Supplementary Table S6), despite both functions calling the same underlying LAPACK routine in FORTRAN (*gesdd*). This surprising result can be explained by the different LAPACK configuration between the two systems: R’s default BLAS/LAPACK on many systems is a reference implementation, which is not optimized for performance. Python, especially in environments like Conda [40], usually links to highly optimized libraries like OpenBLAS [41] or MKL (Intel Math Kernel Library). These optimized libraries use hardware-specific vectorization and threading to speed up linear algebra. In fact, running R’s exact SVD with optimized OpenBlas results in a speed up of up to 30×, leading to computing times similar to those of Python (Supplementary Table S16).

When the input matrix is large and one does not need the full singular values/vectors, truncated SVD algorithms are advantageous compared to computing the exact SVD, both in terms of speed and memory. The fastest approaches are random SVD as implemented in Scanpy, both applied to dense and sparse matrices (1.6 and 2 minutes, respectively, Fig. 3, Supplementary Table S6) and the ARPACK algorithm applied to sparse matrices, especially as implemented in the RSpectra package (1.9 minutes, Fig. 3, Supplementary Table S6). Notably, using a sparse matrix representation as input saves RAM memory, using e.g., 5.6GB for RSpectra sparse, compared to 21.3GB for RSpectra dense. Similar considerations can be made for ARPACK in Scanpy and random SVD in BiocSingular, but surprisingly random SVD in Scanpy uses more RAM memory for sparse matrices than for dense matrices: this is due to Scanpy densifying the matrix prior to running random SVD.

IRLBA is a fast alternative for truncated SVD for in-memory matrices, especially when represented in a sparse format (3.1 minutes, Fig. 3, Supplementary Table S6). However, IRLBA and ARPACK are both very slow when applied to out-of-memory data, exceeding 1.5 hours when applied to HDF5 matrices. This is likely due to the multiple passes on the data needed by these algorithms, which make them suffer from the I/O bottleneck. On the other hand, exact and random SVD perform better for HDF5 than for in-memory inputs, making it the best choice for out-of-memory data: e.g., random SVD applied to dense HDF5 data takes 12 minutes.

We next used the Plackett-Luce model [42, 43] to establish a single, final method ranking, encompassing computational time, memory usage, and the average correlation among the top 50 principal components (Fig. 3). Plackett-Luce is a statistical model for ranked data that estimates a relative “worth” score for each item from its rank across multiple criteria, combining several performance measures into a single overall ranking while accounting for ties. This final ranking clearly highlights four methods that outperform all the other: ARPACK sparse (both implemented in RSpectra and Scanpy), IRLBA sparse, and Rapids dense (both exact and Jacobi).

Finally, we assessed the scalability of each algorithm by taking random subsets of the 1.3M cell dataset and evaluate the time and memory consumption of each method across dataset sizes (Fig. 4). In terms of time, all methods scale linearly with the number of cells. Strikingly, with dense input matrices Python methods are more scalable than R methods. This is largely due to the availability of optimized BLAS libraries, which are not always installed by default across different operating systems. Linking R to an optimized BLAS/LAPACK almost entirely accounts for this difference (see Supplementary Table S16). On the other hand, R is very efficient at handling sparse matrices, thanks in part to the BiocSingular ability to defer centering and scaling (Fig. 4a, Supplementary Tables S3–S5). The IRLBA and ARPACK algorithms are the slowest when the input is out of memory, due to the multiple read accesses (Fig. 4a; see Methods).

**Fig. 4.**
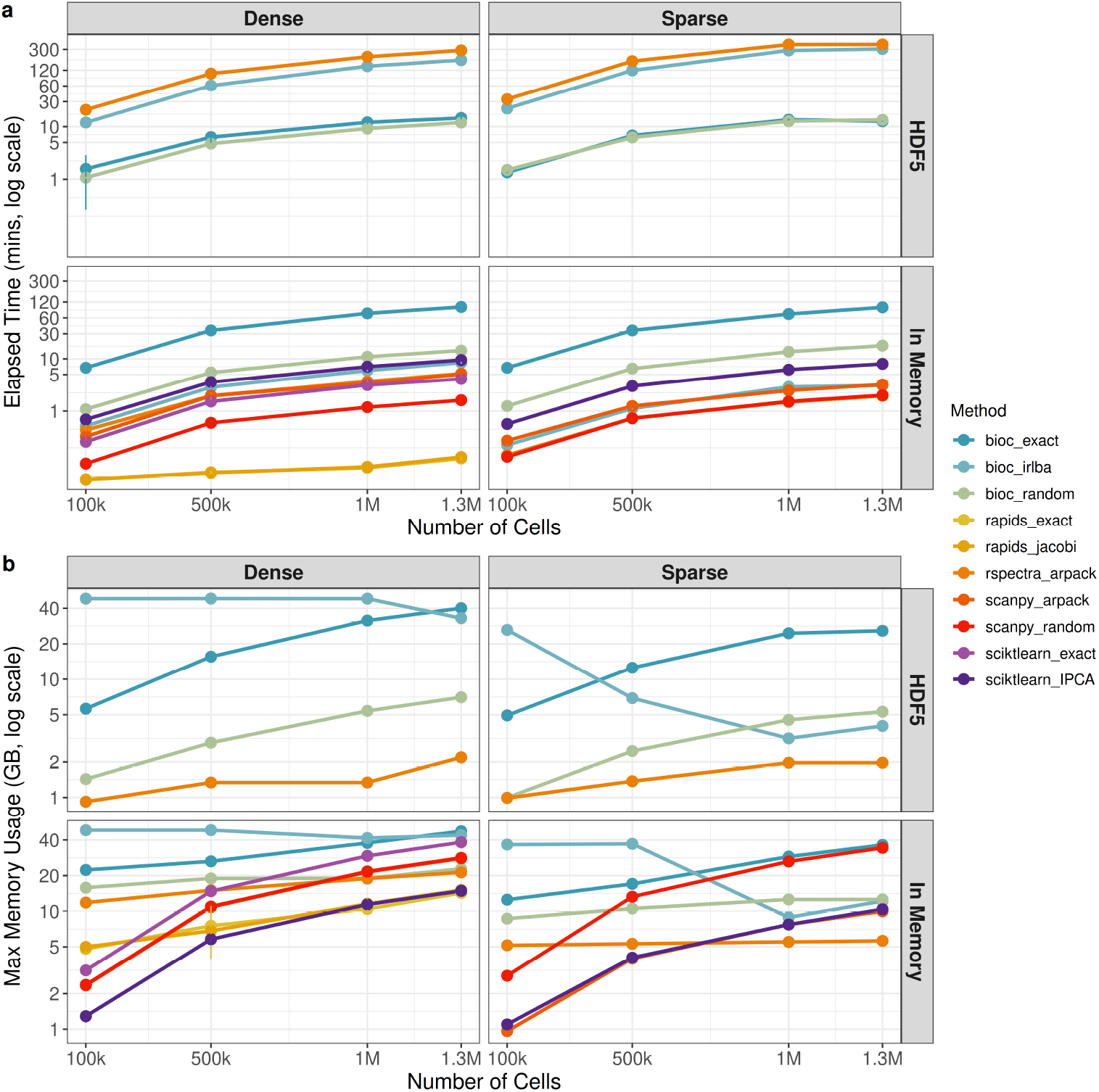
Scalability Assessment of PCA Methods by Input Dimensions, Runtime, and Memory Consumption. (a) Elapsed time (in minutes, log_10_ scale) required to perform principal component analysis (PCA) across a range of dataset sizes (100k, 500k, 1M, 1.3M cells), using different combinations of methods, matrix formats (dense or sparse), and storage types (in-memory or HDF5). (b) Maximum memory usage (in GB, log_10_ scale) required to perform principal component analysis (PCA) across a range of dataset sizes (100k, 500k, 1M, 1.3M cells), using different combinations of methods, matrix formats (dense or sparse), and storage types (in-memory or HDF5).

In terms of RAM memory, as expected, sparse matrices lead on average to lower usage than dense matrices (Fig. 4b, Supplementary Tables S7–S10). Interestingly, random SVD and ARPACK show better RAM scalability than other methods, as evident from the lower slope of the lines (Fig. 4b); this confirms that for larger data sets ARPACK is a good choice. A surprising pattern is observed for IRLBA on sparse matrices (both in and out of memory): RAM usage is much higher for low-dimensional matrices than for much higher dimensions. The IRLBA algorithm involves multiple read access to the input matrix (two per iteration; see Methods) and caching in memory may be employed by the algorithm for smaller matrices to improve computational time.

Taken together, these results show that the choice of the algorithm is critically dependent on the input data and the computing infrastructure. If the data are small enough to fit in the GPU’s VRAM, this is by far the most computationally efficient method; if one cannot use a GPU, but the data fit in the CPU’s RAM, ARPACK, random SVD, and IRLBA are all good choices. If the data are sparse, RSpectra is the fastest and most memory-efficient implementation, while if the data are dense, Scanpy’s random SVD method is the fastest, while incremental PCA is the most memory-efficient. Finally, when the data are out-of-memory, the best approach is random SVD as implemented in BiocSingular, which results in a reasonable time and a low memory footprint.

#### 2.1.3 Application to spatial transcriptomics

Even large scRNA-seq multi-sample studies tipically comprise up to 100k–1M cells, consistent with the scalability study presented in Figure 4. Conversely, imaging-based spatial transcriptomics, which measure gene expression at cell-level resolution on a tissue slide, yields datasets that measure a limited number of genes in up to hundreds of cells per sample, potentially leading to multi-million cell datasets.

To benchmark PCA for such datasets, we applied all the methods to the 4.2M-cell Zhuang-ABCA-1 MERFISH dataset (1,122 genes; see Methods). This allowed us to assess how the same method/input combinations behave when cell number increases by more than 3 times relative to the 1.3M dataset (Supplementary Tables S17–S18 and Supplementary Fig. S3).

Several method/backend combinations that completed comfortably on 1.3M cells, namely rapids singlecell (exact and Jacobi), RSpectra applied to all three HDF5-backed representations, and BiocSingular IRLBA on HDF5-backed sparse matrices, failed to finish within the 48-hour wall-clock limit on the 4.2M-cell matrix and were terminated. In addition, scikit-learn’s exact/full SVD failed outright on the dense representation due to a LAPACK 32-bit indexing overflow, revealing a hard implementation-level ceiling independent of available memory.

Among the 21 method-input combinations that did complete, we computed a combined Plackett-Luce ranking using computational time, memory usage, and average correlation with the exact-SVD reference across PC1–PC50 as judges, following the same approach used for the 1.3M-cell benchmark (Supplementary Fig. S3). Sparse, in-memory CPU methods dominate the top of the ranking: RSpectra sparse ARPACK ranked first overall (6.1 minutes, 7.2 GB), closely followed by Scanpy sparse ARPACK and BiocSingular sparse IRLBA (both under 12 GB and 6-7 minutes), consistent with the 1.3M-cell results.

HDF5-backed dense IRLBA, already slow at 1.3M cells, confirmed its lack of scalability, completing in approximately 34 hours and ranking second-to-last, consistent with the I/O-bound behavior of multi-pass algorithms on out-of-memory data described above. Finally, exact SVD on a dense in-memory matrix ranked last overall despite a moderate runtime (59.7 minutes), owing to its comparatively large memory footprint (165.5 GB), by far the highest of any completed method. As in the 1.3M-cell dataset, PC-level correlation with the exact-SVD reference remained high across implementations, confirming that runtime and memory, rather than approximation quality, are the limiting factors at this scale (Supplementary Fig. S4).

### 2.2 Single-cell RNA-seq Workflow

Having characterized the computational cost of PCA in isolation (Section 2.1), we now turn to how these same algorithmic choices play out within complete, real-world analysis pipelines, alongside a second axis of comparison, i.e. clustering accuracy against ground-truth cell-type labels. We assess each pipeline using its standard analysis steps as recommended by the developers.

In addition to computational time and memory usage, we aim at comparing each workflow’s accuracy. To do so, we selected five datasets that contain different degrees of ground truth, which allows us to evaluate the ability of the workflows to capture real biological characteristics of the analyzed data (Supplementary Table S1).

In particular, we tested the pipelines using the BE1 [44] and sc_mixology [17] datasets, composed of mixtures of cell lines: the cell line of origin of each cell is recorded, making these data useful to evaluate the ability of the workflows to recover known cell clusters. In addition, we used two CITEseq datasets, from Cord Blood [45] and Peripheral Blood [46], which, in addition to scRNA-seq, include cell surface markers, allowing the manual gating of cells for cell typing independent of scRNA-seq; treating these cell types as ground truth, we evaluate the ability of the workflows to recover them using only the scRNA-seq profiles. Specifically, we evaluate the concordance of clustering results, measured by the Adjusted Rand Index (ARI) and the cell-type separation in PCA space (see Methods for details).

To study the computational efficiency and scalability of the methods, we additionally employ the 1.3M cell dataset from 10x Genomics [39]. Taken together, these datasets allow us to test the workflows on different organisms, data throughput, and sample types (Supplementary Table S1).

The time in seconds for each step of the analysis is reported in Supplementary Table S19 and in Figure 5 for each dataset. As expected the computational time is proportional to the dimension of the dataset ranging from a few seconds for sc_mixology to more than 2 hours for the 1.3M cell dataset.

**Fig. 5.**
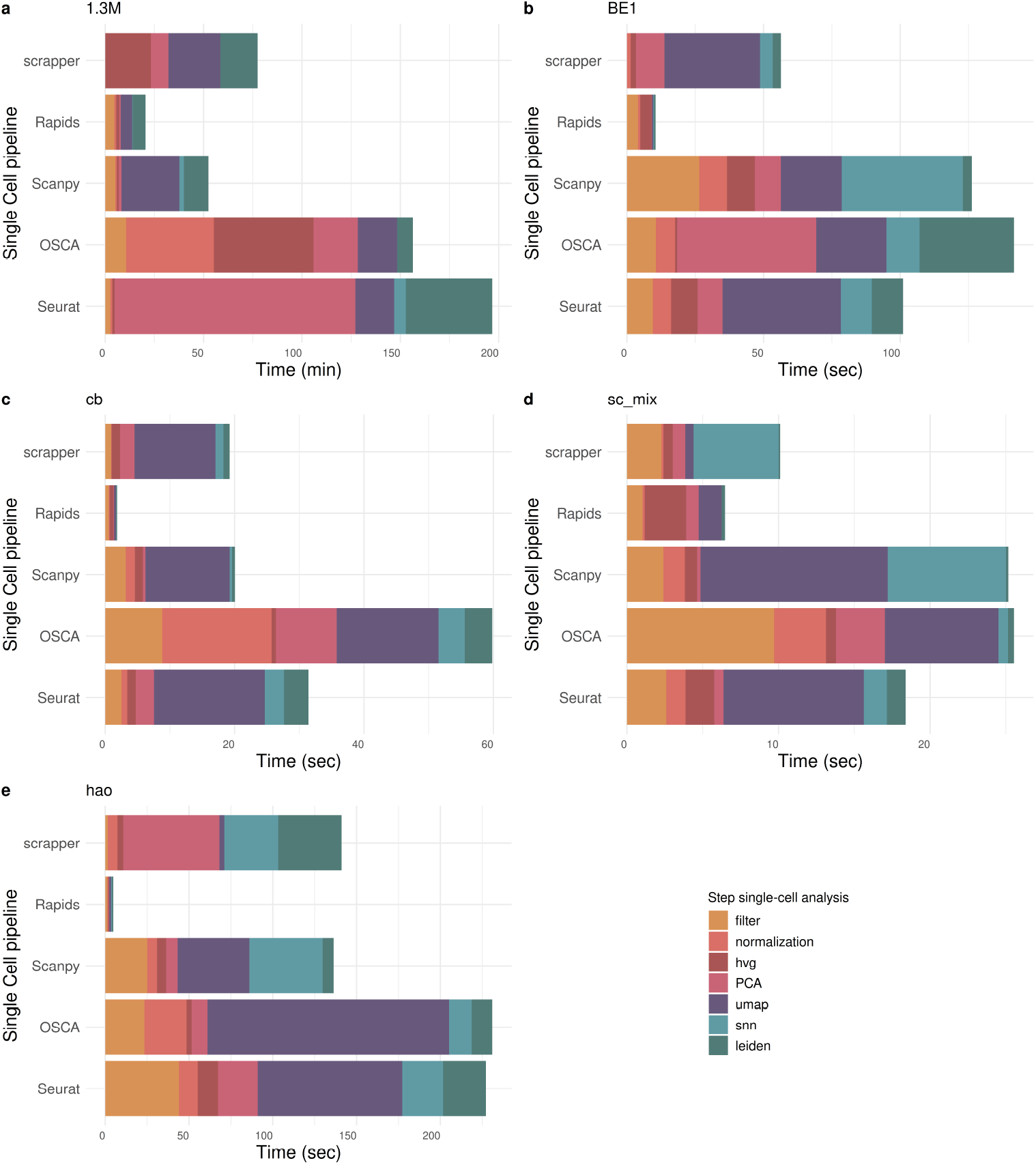
Barplot of elapsed time for each dataset used in the workflow benchmark. (a) Each panel (a–e) displays the computational time required by different single-cell analysis pipelines across various datasets: (a) 1.3M cells, (b) BE1, (c) cb, (d) sc_mixology, and (e) Hao. Pipelines include Seurat, OSCA, Scanpy, Rapids, and Scrapper. Bars are colored by processing step: filtering, normalization, selection of highly variable genes (HVG), PCA, UMAP, snn, and Leiden clustering. Time is expressed in minutes for the 1.3M cell dataset and in seconds for the others.

As expected, the rapids single-cell pipeline is the fastest across all datasets, highlighting the potential of leveraging GPU acceleration for single-cell analysis, as already reported [20]. On the other hand, Seurat and OSCA exhibit the largest computational times. This is particularly evident in the 1.3M cell dataset, with RAPIDS taking about 20 minutes and OSCA and Seurat taking about two and a half and three hours, respectively (Fig 5a). In this dataset, Seurat’s computational performance is hindered by PCA and Leiden clustering, while OSCA’s computational bottleneck is upstream, normalization and highly variable gene (HVG) selection being the most computationally heavy steps (Fig. 5a). The default SVD algorithm used by each workflow, together with its underlying implementation, is summarized in Table S24, and is discussed further below in relation to the performance differences observed here.

For smaller datasets, the gaps are less important, given that all workflows are able to finish the analysis in minutes, if not seconds. Nonetheless, the general trend is confirmed (Fig. 5b-e).

For clarity, Figure 5 reports computational time for UMAP and Leiden clustering only among the processing steps shown; the corresponding timings for t-SNE and Louvain clustering are reported in Supplementary Table S19 and show consistent trends across workflows and datasets. For example, in the BE1 dataset, RAPIDS is the fastest method for both Louvain (1.01s) and Leiden (0.70s) clustering, while OSCA is the slowest for both (103.96s and 29.73s, respectively), mirroring the relative ranking observed for UMAP and Leiden in Figure 5.

Peak memory usage varies substantially across workflows and datasets, and does not always correlate with dataset size (Supplementary Table S20). RAPIDS consistently shows the lowest memory footprint across all four datasets, while OSCA and Scanpy require disproportionately high memory on the BE1 dataset compared to the other workflows.

We next turned to evaluating the accuracy of the clustering derived from each pipeline: Table 1 shows the ARI between the clustering derived from each workflow and the ground truth cell labels, for both the Louvain and Leiden community detection algorithms. Notably, all workflows perform similarly in the sc_mixology (Supplementary Fig. S6), cord blood datasets (Supplementary Fig. S7) and Hao dataset (Supplementary Fig. S8), leading to an average ARI of 0.98, 0.78, 0.60, respectively, reflecting the different complexity of the three datasets. We did not observe any major differences between Leiden and Louvain clustering (Table 1).

**Table 1.** Adjusted Rand Index (ARI) between the workflow and between single-cell RNA-seq dataset. Hao v2: CD4 T, CD8 T and other T cells merged into a single T-cell class.

| ARI | Algorithm | Seurat | OSCA | Scanpy | Rapids | scrapper |
| --- | --- | --- | --- | --- | --- | --- |
| cb | louvain | 0.88 | 0.81 | 0.75 | 0.78 | 0.80 |
|  | leiden | 0.7 | 0.81 | 0.75 | 0.77 | 0.81 |
| sc_mix | louvain | 0.97 | 0.96 | 0.99 | 0.99 | 0.96 |
|  | leiden | 0.98 | 0.96 | 0.99 | 0.99 | 0.96 |
| BE1 | louvain | 0.67 | 0.97 | 0.68 | 0.68 | 0.97 |
|  | leiden | 0.68 | 0.95 | 0.68 | 0.68 | 0.97 |
| Hao | louvain | 0.59 | 0.59 | 0.55 | 0.62 | 0.56 |
|  | leiden | 0.59 | 0.58 | 0.55 | 0.62 | 0.56 |
| Hao v2 | louvain | 0.72 | 0.72 | 0.69 | 0.70 | 0.72 |
|  | leiden | 0.72 | 0.72 | 0.69 | 0.70 | 0.72 |

We next asked whether the five workflows retained the same cells after quality control (Supplementary Fig. S5). The large majority of cells passed filtering in all workflows (92.9% in sc_mixology, 90.1% in BE1, 86.0% in cb, and 91.5% in Hao). The main discrepancy was a set of cells retained only by OSCA and scrapper (5.2%, 8.8% and 7.3% in sc_mixology, cb and Hao, respectively), whereas in BE1 about 9% of cells were retained by Scanpy and Rapids (and in part Seurat) but removed by OSCA and scrapper. Differences in ARI across workflows are therefore unlikely to be driven primarily by differences in the retained cells.

On the Hao dataset, ARI values were lower and more homogeneous across workflows than on the other datasets with ground truth (0.55–0.62; Table 1). CD4^+^ and CD8^+^ T cells had the lowest neighborhood purity among all cell types for every workflow (0.73–0.87), and other T cells were also below the values observed for B cells, dendritic cells and monocytes (Supplementary Table S23 and Supplementary Figure S9d). This shows that T-cell subsets are extensively intermixed in the RNA-based PCA space. Since an RNA-only analysis cannot fully separate them, this label resolution penalizes all workflows. To quantify this effect, we recomputed the ARI after merging CD4^+^, CD8^+^ and other T cells into a single T-cell class (Hao v2 in Table 1). With this lineage-level annotation, ARI increased consistently for all workflows (0.69–0.72), and Seurat, OSCA and scrapper reached the same value (0.72). These results indicate that most of the disagreement with the reference on Hao stems from the resolution of T-cell subsets rather than from differences between workflows.

Of interest is the evaluation of the pipelines on the BE1 dataset, in which OSCA and scrapper achieve an almost perfect concordance with the ground truth, with an ARI of 0.95-0.97, while the other methods stop shy of 0.7 (Table 1). The main reason for this difference is the failure to separate two closely related cell lines, A549 and CCL-185-IG (shown in orange and blue, respectively, in Fig. 6a-e), which is derived from A549 cells [44]. From the t-SNE representations it is evident that OSCA and scrapper separate the two cell lines (orange and blue points) into two distinct clusters, while Seurat, Scanpy and RAPIDS achieve only partial separation between these two populations (Fig. 6a-e).

**Fig. 6.**
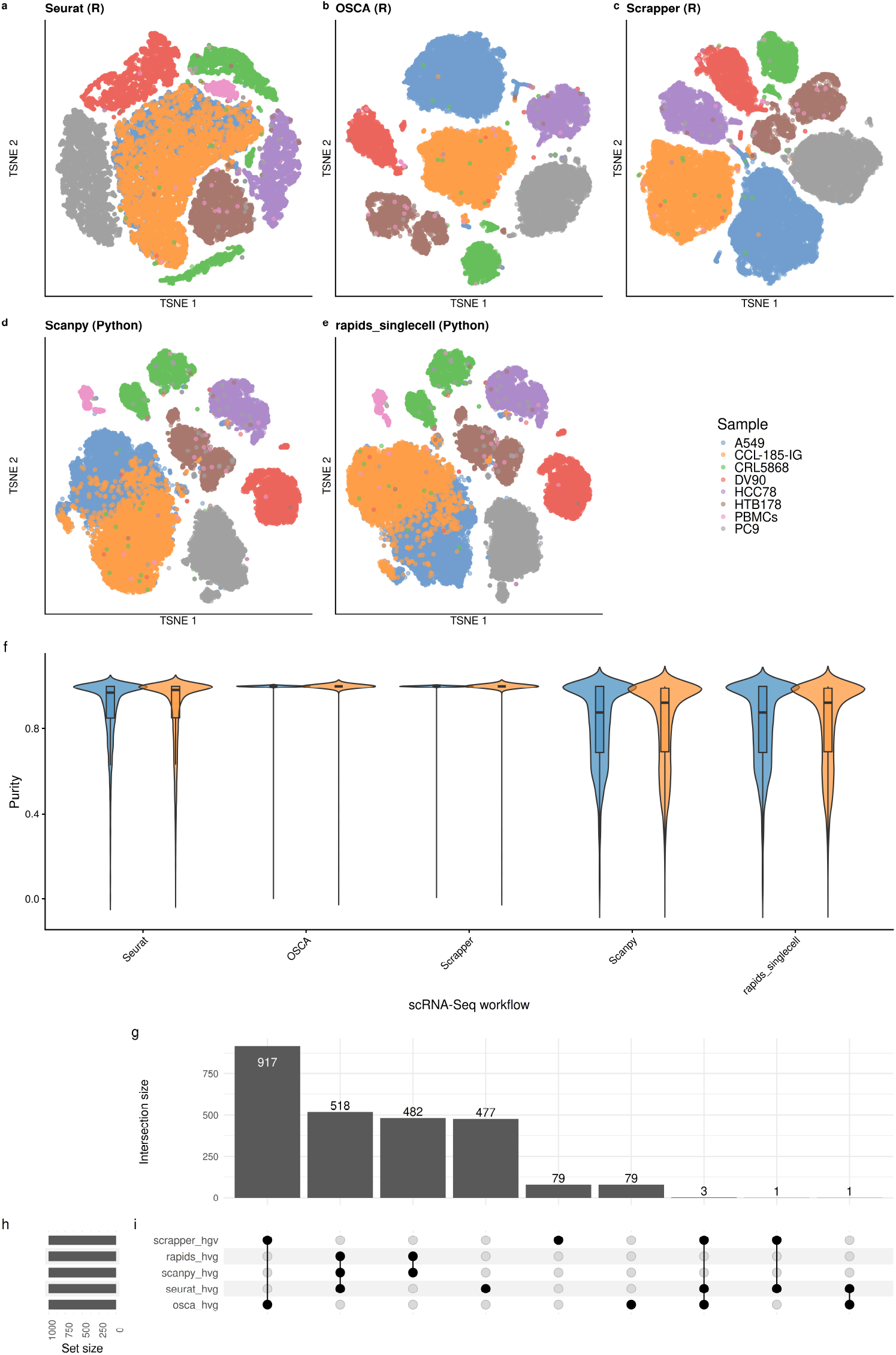
T-SNE plot and HVGs in the BE1 dataset. (a–e) t-SNE embeddings of the BE1 dataset colored by sample identity, generated using five different single-cell workflows: Seurat (a), OSCA (b), Scrapper (c), Scanpy (d), and rapids singlecell (e). Each workflow applies its own normalization and highly variable gene (HVG) selection procedure prior to dimensionality reduction. (f–h) UpSet plot showing the intersection of HVG sets selected by each workflow. Panel (f) indicates the size of each intersection set, (g) shows the number of genes selected per method (set size), and (h) depicts the overlap structure across methods.

To quantify the degree of separation between the two cell types and to ensure that this is not an artifact of the t-SNE representation, we computed the cell type purity in the space of the 50 PCs for each method (Fig. 6f and Supplementary Fig. S9) and the cumulative percentage of variance explained by the cell lines for each PC for each method (Supplementary Fig. S10; see Methods). These analyses confirmed that OSCA and scrapper are better able to separate these two closely-related cell lines.

We next explored the reason why this happens, by exploring the individual steps of the workflows, including QC, normalization, and clustering. There are two major implementation choices that differ between the OSCA-scrapper group and the Scanpy-RAPIDS-Seurat group: first, each workflow uses different ways to account for the mean-variance relation in selecting HVGs (see Methods), leading to a unique set of genes for each method (Fig. 6g); furthermore, Seurat, Scanpy, and rapids singlecell default to scale the data prior to computing the PCA, while OSCA and scrapper only center the data, avoiding scaling them.

The difference in HVG selection reflects distinct strategies for modeling the mean-variance relationship of gene expression. OSCA and scrapper fit a trend to the mean-variance relationship across genes (via `modelGeneVar`/`modelGeneVariances`) and select genes whose variance most exceeds this trend, isolating biological variability from technical noise. In contrast, Seurat’s `vst` method standardizes gene expression after estimating a local mean-variance relationship, while Scanpy and rapids singlecell rely on dispersion-based binning of genes with fixed mean and dispersion thresholds (see Methods).

To directly test whether the observed results depend on these two algorithmic choices, we repeated the Seurat and Scanpy workflows, with two modifications that we tested separately and jointly: (i) using the HVGs selected by OSCA in place of each workflow’s own default HVG selection method; (ii) avoiding scaling the data prior to PCA. Strikingly, each modification by itself did not improve the ARI substantially (Supplementary Table S21) and only jointly modifying the HVG selection and scaling method resulted in an improved accuracy, much closer to that of OSCA and scrapper (ARI of 0.96; Supplementary Table S21). Cell line purity in PCA space showed a consistent but more graded pattern (Supplementary Table S22). Each modification alone partially increased the purity of A549 and CCL-185-IG cells (e.g., from 0.83 to 0.94 for CCL-185-IG in Scanpy when using the OSCA HVGs). This suggests that the separation between the two cell lines improves gradually, but is recovered by graph-based clustering only when it is nearly complete. Jointly applying both modifications raised the purity of both cell lines to ≥ 0.998 in Seurat and Scanpy, matching OSCA. Conversely, introducing scaling in the OSCA workflow reduced the purity of A549 and CCL-185-IG to 0.95 and 0.93, respectively, and the ARI to 0.91. These values are comparable to those of Seurat and Scanpy with OSCA HVGs and scaling. These results indicate that HVG selection, in combination with data scaling before PCA, drives the accuracy gap observed on BE1.

Overall, the OSCA and scrapper approach to model the gene-variance trend, combined with no scaling prior to computing the PCA, seem to achieve greater performance than the alternative methods (see also Supplementary Figure S9 and Supplementary Table S23).

Analogous t-SNE embeddings, colored by ground-truth cell type, for the sc_mixology, cord blood, Hao datasets are shown in Supplementary Figures S6, S7 and S8, respectively, and show similar trends to those observed for BE1.

Taken together, these results show how the algorithmic choices of the analysis workflows can impact the downstream results and how the selection of HVGs and scaling are important steps in the pipeline.

Figure 7 summarizes the trade-offs between computational time, memory usage, and clustering accuracy across the four datasets with ground-truth cell-type annotations. This plot confirms that no single workflow dominates across all criteria: rapids singlecell consistently ranks first or second on both memory usage and computational time, while OSCA or scrapper typically rank first for ARI with ground-truth cell-type annotations (Table 1); Seurat and Scanpy generally occupy intermediate positions. Importantly, this figure highlights that the decision on the most appropriate method depends on the complexity of the clustering task: all workflows work well on the simple task of distinguishing well-separated cell lines (e.g., sc_mixology), while when in need to separate closely-related cell types (e.g., cord blood or BE1), a slower but more accurate method may be needed.

**Fig. 7.**
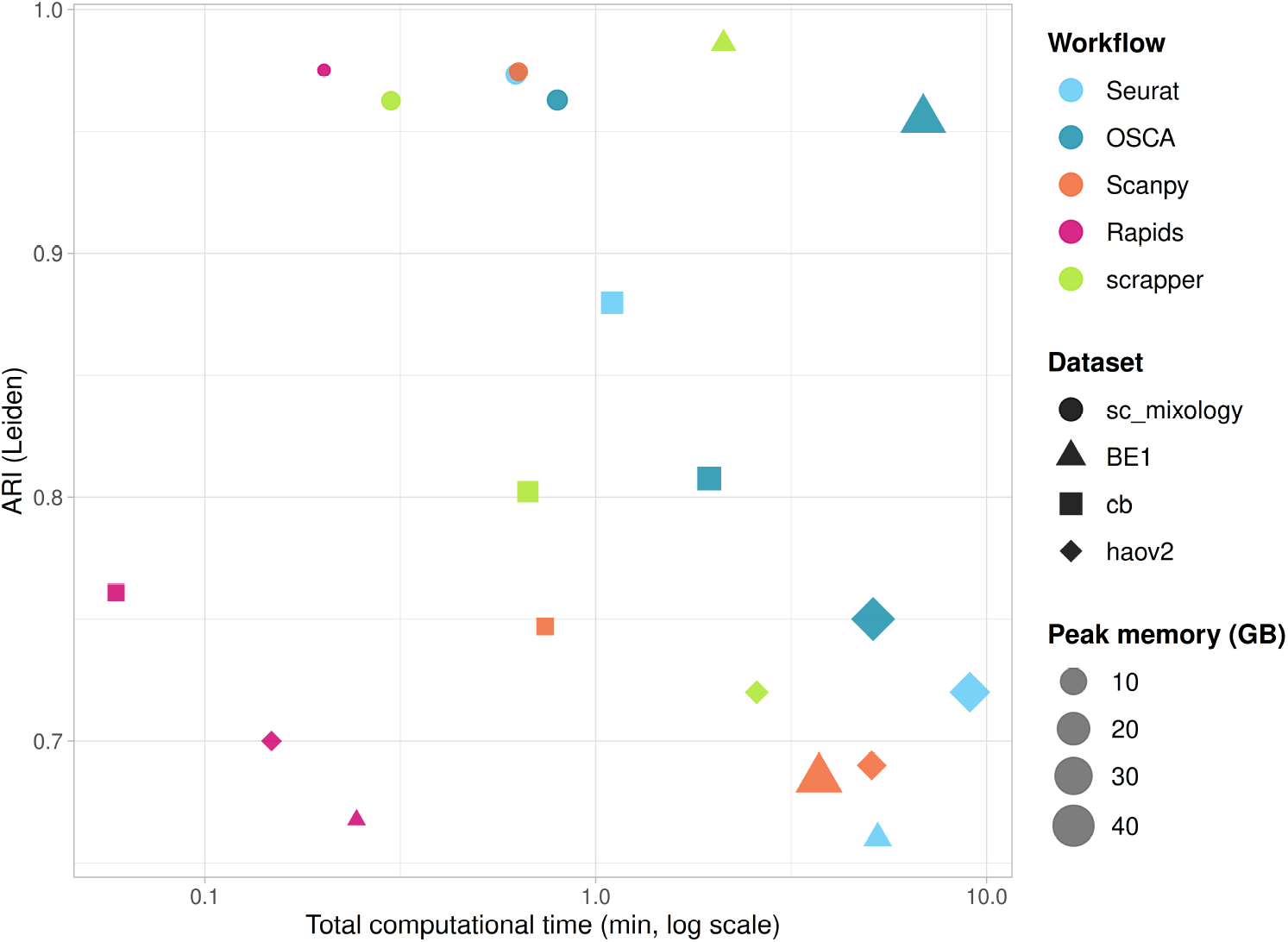
Trade-off between computational time, memory usage, and clustering accuracy across single-cell workflows. Each point represents one workflow (color: Seurat, OSCA, Scanpy, Rapids, scrapper) applied to one of the four datasets with ground-truth cell-type annotations (shape: sc_mixology, BE1, cb, Hao v2). The x-axis reports the total computational time (min, log_10_ scale), the y-axis the Adjusted Rand Index (ARI) between Leiden clustering and the ground-truth labels, and point size is proportional to peak memory usage (GB). Points closer to the top-left corner, and smaller in size, indicate a better balance between efficiency and accuracy. For the Hao dataset, the lineage-level annotation (Hao v2), in which CD4^+^, CD8^+^ and other T cells are merged into a single T-cell class, is shown (see Table 1).

## 3 Discussion

The ever-increasing size of single-cell datasets and the increasing availability of singlecell atlases has rendered the scalability of workflows, both in terms of speed and memory usage, crucial. Here we compared five single-cell workflows, including the popular Seurat and Scanpy packages, as well as more scalable alternatives, in terms of accuracy and scalability. Focusing on PCA as an exemplar step, we demonstrated that the type of input, the choice of algorithm, and the software configuration all critically influence the scalability of the methods.

The differences in PCA performance characterized in Section 2.1 (Fig. 3, Table S24) directly help explain part of the performance gap observed between full workflows (Fig. 5), as both analyses are performed on a comparable number of highly variable genes (1,000). For example, for the 1.3M cell dataset, PCA is the single most time-consuming step for Seurat (Fig. 5a), consistent with its reliance on ARPACK (through the BPCells package [47]), which is among the slowest methods for out-of-memory data in our PCA benchmark (Fig. 3). Conversely, OSCA uses randomized SVD on data stored on disk in HDF5 format, which is much faster (Fig. 3 and Fig. 5).

Out-of-memory approaches may be useful when dealing with very large data. However, HDF5-backed methods’ performance is affected by the alignment between on-disk chunk boundaries and the blocks of data actually read. HDF5 files store array data in fixed-size, typically rectangular chunks, and any operation touching even a single element of a chunk requires that entire chunk to be read (and, if modified, decompressed and rewritten) [38]. When a subset of cells is extracted from a larger HDF5-backed dataset and written to a new HDF5 file, as in the case of our 100k/500k/1M subsamples of the 1.3M-cell dataset, the resulting chunk boundaries may no longer align with the structure of the original file. Consequently, a given operation may need to read more chunks – and hence more data – than would be strictly necessary for an optimally chunked dataset of the same size.

This chunk-misalignment effect compounds the inherent I/O overhead of multi-pass, out-of-memory algorithms such as IRLBA and ARPACK discussed above, and may contribute to some of the non-monotonic timing patterns we observe across dataset sizes for HDF5-backed methods (Fig. 4). We did not re-optimize chunk dimensions for each subsampled dataset, as doing so would depart from the experience of a typical user working with default parameters rather than a specifically hand-tuned analysis; users working with performance-critical, repeatedly-subsetted HDF5-backed data may want to consider re-chunking their files to match their access pattern.

Accelerating single-cell analyses with GPUs is a promising avenue for the near future: while GPUs are routinely used in deep learning, also in the context of singlecell analyses [48], their use to accelerate matrix multiplications and other common steps in the analysis workflows is still not widespread. One notable exception is the RAPIDS single-cell library, that we proved accurate and scalable in terms of both time and memory usage. In fact, their use in PCA calculations results in a speed up of approximately 15× compared to the fastest CPU alternative. More work is needed to make it easy for R/Bioconductor developers to leverage GPU computations in their packages: the GPUMatrix [49] and anvl [50] R packages are promising starting points to achieve this goal.

One limitation of GPU is the amount of VRAM available: while the datasets used in this benchmark were small enough to fit in VRAM, for larger datasets the amount of memory needed is a critical factor for the use of GPU, as larger-than-VRAM datasets may lead to out-of-memory errors or result in slower performance due to data transfer between RAM and VRAM.

Applying our PCA benchmark to the 4.2M-cell Zhuang-ABCA-1 MERFISH dataset, we found that even a moderate 3× increase in the number of cells had a dramatic effect: several GPU- and HDF5-backed methods that performed competitively on 1.3M cells exceeded a 48-hour wall-clock limit or failed outright (see Section 2.1), including a hard LAPACK 32-bit indexing ceiling for scikit-learn’s exact SVD that is unrelated to available memory. This suggests that, as single-cell and spatial datasets continue to grow, implementation-level limits will need explicit consideration alongside algorithmic computational complexity.

Looking ahead to datasets an order of magnitude larger than those considered here, we see three main bottlenecks. First, a RAM-versus-time trade-off becomes unavoidable once data exceed GPU VRAM: rapids_singlecell/rmm supports managed memory, which oversubscribes VRAM by spilling to host RAM, but this comes at a performance penalty that grows with the amount of spilled data, and out-of-core execution via Dask trades further speed for the ability to process datasets that do not fit in memory at all [51].

Second, sparse matrix representations remain the most memory-efficient choice for droplet-based scRNA-seq, but they are not a universal solution at extreme scale: on GPU, standard sparse formats use 32-bit indices and are therefore limited to at most 2^31^ − 1 explicitly stored non-zero values per matrix (or per Dask block), a hard ceiling that is independent of available VRAM or RAM and that datasets approaching hundreds of millions of cells may begin to approach [51]. Row-chunked, Dask-based execution can sidestep this per-block limit, at the cost of the added complexity and overhead of distributed, out-of-core computation.

Third, our results on both CPU and GPU point to data transfer and I/O, rather than the arithmetic of the SVD itself, as an increasingly dominant cost at scale: HDF5-backed CPU methods that require multiple passes over out-of-memory data (e.g., IRLBA, ARPACK) suffered the largest slowdowns in our benchmark, and the same consideration are likely to apply to GPU pipelines, where repeated CPU–GPU data transfer, rather than GPU compute throughput, is often the limiting factor. Taken together, we believe that future benchmarking and methods-development efforts should explicitly target these three axes: memory-tiering strategies (managed memory, out-of-core/Dask), sparse-format scalability limits, and I/O-aware algorithm design.

BLAS/LAPACK optimization has, perhaps not surprisingly, a profound impact on the computational performance of the methods. Indeed the same exact R code can be sped up by up to 30× by simply linking to an optimized BLAS/LAPACK instead of using the default reference implementation. In this benchmark we have decided to keep the default BLAS/LAPACK version in R and Python to mimic the experience of the typical user, who will likely keep the default R installation, perhaps not even realizing that such an important performance gain can be achieved with a different BLAS/LAPACK implementation. Nonetheless, our recommendation is to run R with an optimized BLAS/LAPACK and we provide instructions to do so in Ubuntu with a simple *apt* call (see Methods).

The BE1 case study (Section 2.2) illustrates that HVG selection can be a major driver of accuracy differences between workflows. We caution, however, against generalizing this result as a universal ranking of HVG selection methods: on the other three datasets with ground-truth annotations (cb, sc_mixology, Hao), workflows achieve comparatively similar accuracy despite using the same different HVG selection strategies (Table 1). This indicates that the relative importance of HVG selection, cell/gene filtering, and other pipeline choices (SNN graph construction, choice of clustering algorithm, clustering resolution) is dataset-dependent rather than fixed, and that no single step can be assumed a priori to dominate accuracy differences across all single-cell datasets. This shows that benchmarking a full workflow, rather than any single step in isolation, is necessary to identify which choice matters for a given dataset.

While, like most benchmarks in this space, our comparison is necessarily tied to the specific packages, versions, and datasets we selected at the time of writing, our code repository includes container and conda-environment definition files that pin every software dependency used in this study, along with instructions to rebuild the computing environments and re-run any part of the benchmark (see Code availability), thereby lowering the barrier for others to adapt and extend our work.

Finally, we have chosen to benchmark the most basic workflow available from each of the software frameworks, mimicking the “basic tutorials” or “getting started” guides. Obviously, modern single-cell RNA-seq studies require more complex analyses, involving multi-sample comparisons, batch-effect removal [52, 53], reference-based annotation [54], and multi-modal integration [46], to name a few. It is likely that the impact of optimized code and GPU acceleration is even more important for these more complex algorithms.

## 4 Methods

### 4.1 An introduction to PCA

Here, we briefly describe PCA and its relation to SVD. We refer the reader to [22] and [23] for a more thorough introduction and the mathematical details.

Let *X* be the *n*×*m* matrix that contains the log-transformed, normalized expression of *m* genes in *n* cells. We assume that the matrix *X* has been centered such that each row has mean 0.

The principal components are the eigenvectors of the covariance matrix and can be computed by its eigendecomposition. Alternatively, the PCs can be computed directly by the Singular Value Decomposition (SVD) of the original matrix *X*. Specifically, we construct the following decomposition of *X*:

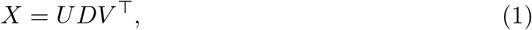

where *U* is a *n* × *p* orthogonal matrix of left singular vectors, *V* is a *m* × *p* orthogonal matrix of right singular vectors, and *D* is a *p* × *p* diagonal matrix, whose elements *d*_1_ ≥ *d*_2_ ≥ *. . .* ≥ *d_p_* ≥ 0 are the singular values. The first columns of *UD* correspond to the principal components, ordered by decreasing explained variance.

### 4.2 Algorithms to compute the SVD

#### 4.2.1 Exact algorithm

In R, an exact SVD algorithm is implemented in the base `svd` function, which is called by the *BiocSingular* package’s `runPCA` function when selecting the “exact” method. In Python, *scikit-learn*’s PCA function calls the `scipy.linalg.svd` function. Both R’s and Python’s svd functions internally call the `gesdd` funcion in the LAPACK (Linear Algebra Package) library in Fortran. This function implements an efficient divide-and-conquer algorithm to compute the SVD [55].

#### 4.2.2 Truncated SVD

Often, one is interested in computing only a few, say *k*, singular values and vectors. For instance, in scRNA-seq, we are often limiting ourselves in calculating the top 30 or 50 PCs. Specifically, we may solve the problem:

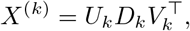

where *U_k_* and *V_k_* denote the leading *k* columns of *U* and *V*, respectively, *D_k_* denotes the diagonal matrix with the *k* largest singular values. It can be shown that *X*^(*k*)^ is the best rank-*k* approximation of *X*.

When *k* ≪ min{*n, m*}, as in the case of scRNA-seq analysis, truncated SVD algorithms exist to efficiently calculate the *k* largest singular values and the corresponding singular vectors. Here, we review a few of these methods.

##### ARPACK

ARPACK [28] is a collection of Fortran77 subroutines designed to solve large-scale eigenvalue problems. ARPACK stands for ARnoldi PACKage and is based on the Arnoldi iteration method [56]. This method provides a good approximation of the *k* largest singular values and the corresponding singular vector of a matrix by constructing an orthonormal basis of its Krylov subspace [56]. Thanks to the relation between eigendecomposition and SVD, the ARPACK algorithm can be used to efficiently calculate the *k* largest singular values and the corresponding singular vectors.

Consider the decomposition in (1), one can show that

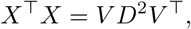

and that *U* = *XV D*^−1^. Thus, the singular values and singular vectors of *X* can be computed from the eigenvalues and eigenvectors of *X*^⊤^*X*.

The Arnoldi method is particularly useful when dealing with large sparse matrices, as the algorithm does not require explicitly the whole matrix, but only a matrix-vector multiplication result. Therefore, if the matrix-vector product can be computed efficiently, which is the case when *X* is sparse, ARPACK will be very efficient for large-scale matrices.

The ARPACK Fortran routine is used by Scanpy’s PCA function through Scipy’s `svds` function. In R, the *RSpectra* function provides an interface to the Spectra C++ library [35] that reimplements the ARPACK algorithms. Since Spectra implements the same underlying algorithm, we refer to both methods as ARPACK in the main text.

##### Random SVD

The randomized SVD algorithm [29] computes a near-best rank *k* approximation of a matrix using matrix-vector products with standard Gaussian vectors. The randomized SVD algorithm uses a random projection matrix to sample the column space of the original matrix, allowing approximation of the SVD of the original matrix by computing SVD on a smaller matrix.

In detail, given a *n* × *l* orthonormal matrix *Q*, with *k* ≤ *l* ≤ *m*, such that *X* ≈ *QQ*^⊤^*X*:

- Form *B* = *Q*^⊤^*X*;
- Compute the SVD of *B*, i.e., *B* = *Ũ*Σ*V* ^⊤^;
- Set *U* = *QŨ*.

Since *X* ≈ *QQ*^⊤^*X* = *Q*(*Ũ*Σ*V* ^⊤^), it follows that *X* ≈ *U* Σ*V* ^⊤^. This algorithm is efficient when *l* ≪ *m*, because we can efficiently compute the matrix *B* and then we just need to compute the SVD of a much smaller matrix.

The randomness comes from the construction of the orthonormal matrix *Q*, which is obtained by randomly projecting the original data matrix onto a low-dimensional space, using *l* randomly generated Gaussian random vectors [29]. The matrix *Q* can be a poor approximation of the column space of the input matrix; to improve the approximation, oversampling (i.e., choosing *l > k*) and power sampling iterations can be used. Briefly, instead of acting directly on *X*, the data matrix is first preprocessed as *X*^(*q*)^ = (*A*^⊤^*A*)*^q^A*, where *q* is the number of power iterations. The choice of *q* represents a trade-off between accuracy and computational efficiency, with larger values corresponding to a more precise approximation at a larger computational cost [57].

The randomized SVD algorithm is implemented in the *rsvd* R package, internally called by *BiocSingular* and in the `truncatedSVD` function in *scikit-learn*, which is called by Scanpy. Notably, *rsvd* uses *q* = 2 power iterations by default, while *scikit-learn* defaults to *q* = 4 or *q* = 7 depending on the number of required componenents.

##### IRLBA

The Augmented Implicitly Restarted Lanczos Bidiagonalization Algorithm (IRLBA) [31] is a fast and memory-efficient method for computing truncated SVD of a matrix. It is based on the augmented implicitly restarted Lanczos bidiagonalization algorithm, which is a variant of the Lanczos algorithm that uses a bidiagonalization step to reduce the dimensionality of the matrix.

IRLBA computes sequences of projections of *X* onto specific low-dimensional subspaces that reduce the dimensionality of the problem. Restarting is implemented to iteratively reduce the approximation error.

More specifically, the algorithm applies *l* steps of partial Lanczos bidiagonalization to *X* and yields the decomposition

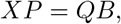

where *P* and *Q* are orthonormal matrices of dimension *m* × *l* and *n* × *l*, respectively, while *B* is upper bidiagonal. An approximate SVD of *X* can be obtained starting from the SVD of *B* and the matrices *P* and *Q* [31].

When *X* is large, the storage requirement of the partial Lanczos bidiagonalization is large, unless the number of Lanczos bidiagonalization steps is small. However, for a small value of *l*, the SVD of *X* may be approximated poorly by the algorithm. To avoid this problem, the algorithm uses a sequence of initial vectors to form *P*, a technique known as restarted partial Lanczos bidiagonalization. Finally, to avoid numerical instabilities, restarting is carried out by augmentation of Krylov subspaces (see [31] for details).

The IRLBA algorithm is implemented in the *irlba* R package, internally called by *BiocSingular*.

The IRLBA algorithm is implemented in the *irlba* R package, internally called by *BiocSingular*. Notably, Seurat’s default PCA implementation (`RunPCA`) also relies on the same *irlba* package, calling the `irlba` function directly on the scaled, HVG-restricted expression matrix. The scrapper package’s `runPca` function also uses IRLBA by default, but via a derived C++ implementation provided by the *libscran*/*irlba* library [58].

#### 4.2.3 IPCA

Incremental PCA [33] is a method for computing the principal components of a matrix in an incremental manner. This means that the algorithm processes the data in batches, computing the principal components for each batch and then combining them to obtain the final result. This approach avoids the need to load the entire dataset into memory, making it suitable for large-scale data analysis. The basic idea is to update the principal components incrementally as new data batches become available.

Here, we briefly illustrate the algorithm, assuming two batches of data. Let us denote with *X_A_* and *X_B_* the two batches of data. For the first batch, we compute the SVD in the usual way, say, *X_A_* = *U_A_D_A_V* ^⊤^. The goal is to find the SVD of the concatenation of *X_A_*and *X_B_*, which can be expressed as

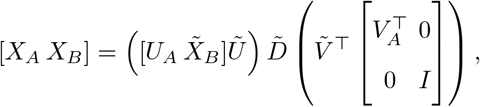

where *X̃_B_* is the component of *X_B_* orthogonal to *U_A_* and *R̃* = *ŨD̃Ṽ*^⊤^ is the SVD of a matrix of size *k* + *d*, with *d* the number of observation in the batch *X_B_*, which can be computed efficiently even for large *n*. The algorithm is slightly more complicated because it needs to take into account the fact that the sample mean of the data will change as more batches are added, the full details are in [33].

The IPCA algorithm is implemented in the `IncrementalPCA` function in *scikit-learn*.

#### 4.2.4 Jacobi

The Jacobi SVD algorithm [32] is an iterative algorithm that applies a sequence of plane rotations to a symmetric matrix to bring it to diagonal form. The Jacobi SVD algorithm is particularly known for its numerical stability and accuracy, making it suitable for high-precision applications.

The Jacobi method is an algorithm for the diagonalization of symmetric matrices and can be used to compute the SVD of general matrices. Indeed, applying the Jacobi method to *H* = *XX*^⊤^ leads to the sequence *A*^(*k*+1)^ = *A*^(*k*)^*V* ^(*k*)^, whose limit matrix is *UD*, while the accumulated product of Jacobi rotations is *V* [32].

The Jacobi algorithm is implemented in the *RAPIDS* library.

### 4.3 SVD Performance Evaluation

#### 4.3.1 SVD Accuracy

To evaluate the degree of similarity among the principal components obtained from different SVD algorithms, we computed the absolute Pearson correlation coefficient between the PC computed with the “exact” method and the corresponding PC derived from all other methods. The Pearson correlation coefficient measures the linear correlation between two variables, ranging from -1 to 1, where values close to zero indicate weak or no linear correlation, and values close to one indicate a strong linear correlation. By computing the absolute value of the correlation we account for PCA’s invariance with respect to rotations. High values of absolute correlation coefficients indicate high similarity in the patterns captured by the principal components.

#### 4.3.2 Computational Time

To measure the computational time for PCA computation in R, we employed the `proc.time` function, capturing the start and end times for each computational step. Conversely, in Python, we imported the time module and used the `time.time` function to capture the start time of the process, then using the `sys.argv` command we calculate the end time of the process.

#### 4.3.3 Memory Usage

To monitor memory usage efficiently, we employed different strategies in R and Python. In R, we used the `Rprof` tool to profile memory consumption during program execution. With Rprof, we were able to track the memory consumption and then identify the peak of memory usage. Conversely, in Python, we used the `/usr/bin/time -v` command within our Slurm job scripts to measure and record the maximum memory usage. By appending this command to our job scripts, we captured memory usage statistics at the process level, allowing us to monitor and analyze resource utilization effectively.

#### 4.3.4 Ranking SVD methods

The Plackett-Luce model [42, 43] provides a flexible framework for modeling preferences and rankings in various domains. It handles ties of arbitrary order in the ranking. This means that the model can accommodate rankings in which items are tied for a particular rank. To model such rankings, we utilized the *PlackettLuce* R package, which extends the traditional Plackett-Luce model by incorporating methods that accommodate tied rankings. Using the `as.rankings` function, the data were restructured into a format suitable for analysis by converting preferences into rankings. The functions used included the weights argument, which allows for the weighting of observations based on the frequency of preferences, and the `log = FALSE` option, which transforms the scores into normalized probabilities, ensuring that their sum equals one. This step is crucial, as the first item in the ranking is treated as the reference for others, allowing the calculation of relative probabilities for each item to be placed in the top position.

Subsequently, we performed a summary analysis of the results using the `summary` function, which provided detailed information on the estimated coefficients of the model. To visualize the relative importance of each method, we used the `qvcalc` function, which calculates the worth values for each method, enabling the graphical representation of the results.

#### 4.3.5 Default SVD algorithms across workflows

Table S24 summarizes the default SVD algorithm and underlying implementation used for PCA by each of the five benchmarked workflows, together with the corresponding method evaluated in the PCA benchmark (Section 2.1, Fig. 3). This mapping allows the results of the PCA benchmark to be related to the performance differences observed between full workflows (Section 2.2).

### 4.4 A typical workflow for Single Cell RNA-seq data analyses

Here, we describe the steps of a typical workflow for single-cell RNA-seq (scRNA-seq) data. These steps are common to all five pipelines that we benchmarked in this paper. After a general description of all the steps, we describe how each specific workflow differ in the implementations of these steps.

#### 4.4.1 Preprocessing

Low-quality libraries in scRNA-seq data can result from various sources, such as cell damage during dissociation or errors in library preparation (e.g., inefficient reverse transcription or PCR amplification). These typically manifest as “cells” with low total counts, few expressed genes, and high proportions of mitochondrial reads, leading to misleading results in downstream analyses [59].

After excluding empty droplets and identifying potential doublets, droplets containing damaged cells or low-quality reads are filtered out. Droplets are small compartments that capture single cells and barcoded beads during scRNA-seq experiments. Empty droplets lack cells, while doublets occur when two cells are mistakenly captured in the same droplet, creating mixed signals that can compromise data accuracy.

The library size, defined as the total sum of counts across all relevant features for each cell, is a commonly used metric for filtering. Cells with small library sizes are more likely to be of low quality due to RNA loss during library preparation, either from cell lysis or inefficient cDNA capture and amplification. Another metric is the number of expressed features per cell, defined as the number of endogenous genes with non-zero counts for that cell. Cells with very few expressed genes are likely of poor quality as the diverse transcript population has not been successfully captured. The proportion of reads mapped to mitochondrial genes can also be used; high proportions suggest possible loss of cytoplasmic RNA due to cell damage, as mitochondria, being larger than individual transcript molecules, are less likely to escape through cell membrane holes.

#### 4.4.2 Normalization

Normalization aims to adjust raw counts for variable sampling effects by scaling the observed counts. This process helps to reduce technical noise and preserve biological variation. Systematic differences in coverage between libraries in scRNA-seq data, often due to variations in cDNA capture or PCR amplification efficiency, can interfere with expression profile comparisons. Normalization removes these differences, ensuring accurate clustering and differential expression analyses [60].

#### 4.4.3 Feature selection

Feature selection aims at reducing the dimension of the input matrix to identify genes whose expression is variable enough across cells to be able to capture most of the cell heterogeneity filtering out noise. Methods such as dimensionality reduction and clustering are generally performed using the filtered matrix [61–63].

#### 4.4.4 Dimensionality Reduction: PCA, t-SNE, UMAP

Hao = c(Seurat = 0.59, OSCA = 0.58, Scanpy = 0.55, Rapids = 0.62, scrapper = 0.56) Dimensionality reduction techniques are typically used for the visualization of complex datasets. These methods aim to transform high-dimensional data into a lower-dimensional space preserving the underlying structure and variability inherent in the data.

Principal Component Analysis (PCA) is widely employed in scRNA-seq to identify orthogonal axes (principal components) that capture the maximum variance in the dataset. By projecting cells onto these components, PCA provides a simplified representation that retains the most significant sources of variation.

In contrast, t-Distributed Stochastic Neighbor Embedding (t-SNE) [64] focuses on preserving local similarities between cells in the high-dimensional space. It maps cells into a low-dimensional space (often 2D) that aims at placing similar cells together. Uniform Manifold Approximation and Projection (UMAP) [65] is a more computationally efficient alternative that has gained popularity in recent years.

It is a standard practice that both t-SNE and UMAP take as input the first 50 PCs obtained from the pre-processed gene expression matrix, where 50 is a number of PCs that should guarantee to capture most of the cell heterogeneity present in the data.

#### 4.4.5 Clustering: Louvain, Leiden

Clustering approaches such as Louvain [66] and Leiden [67] are used to identify discrete cell populations based on gene expression patterns.

These methods operate on a graph representation of the data, often constructed using k-nearest neighbors (KNN) or shared nearest neighbors (SNN) algorithm, where nodes represent cells, and edges represent similarities or connections based on gene expression profiles.

Although graph construction and community detection are conceptually distinct steps, they are often executed as a single call in standard pipelines. In this benchmark, we measured the computational time of these two steps separately for all five workflows, in order to assess their individual contribution to the overall clustering time (Fig. 5); the specific functions used for graph construction and for each clustering algorithm are detailed for each workflow in the following subsections.

An important concept to this clustering approaches is modularity, which is used to evaluate and optimize the grouping of cells into meaningful clusters, also referred to as communities. Modularity is a measure used to evaluate how well cells are grouped into clusters based on their gene expression profiles. It quantifies the quality of clustering by comparing the density of connections (or similarities) between cells within the same cluster to those between cells in different clusters. Higher modularity indicates that cells within a cluster are more tightly connected (e.g., share similar gene expression patterns) compared to connections between clusters. Lower modularity implies that the boundaries between clusters are less distinct, potentially reflecting noise or suboptimal clustering. In the single-cell RNA-seq context, communities refer to groups of cells that are more connected or similar to one another based on similarity metrics, such as shared gene expression patterns, than they are to cells in other groups. These communities may represent biologically meaningful structures, such as cell types, subtypes, or states, within scRNA-seq datasets.

### 4.5 Workflows for Single Cell RNA-seq data analyses

#### 4.5.1 Seurat

Seurat, developed and maintained by the Satija Lab, is an R toolkit tailored for single-cell genomics. For our analysis, we utilized Seurat version 5 [19].

Specific steps and default values of the Seurat workflow include:

- Find Mitochondrial Genes: This step identifies mitochondrial genes by calculating the proportion of transcripts mapping to mitochondrial genes, using the `PercentageFeatureSet` function with the regex pattern `"^MT-"`or `"^mt-"` from *Seurat*. Cells with high mitochondrial content (less than 5 %) or low total counts are flagged and filtered out.
- Filtering: A filtering step is applied to remove cells and genes with undesired properties. To do the subset is used the `subset` function from *SeuratObject*. Cells are retained if they express between 200 and 5000 features, have mitochondrial content below 5%, and a total count under 25,000 for BE1 dataset. Cells are retained if they express between 200 and 2500 features, have mitochondrial content below 5%, and a total count under 4,000 for cord blood dataset. Cells are retained if they express between 200 and 6200 features, have mitochondrial content below 5%, and a total count under 60,000 for sc_mixology dataset.
- Normalization: The data is normalized using the `LogNormalize` method, which scales gene expression counts by the total expression for each cell, followed by a log transformation to stabilize variance.
- Highly Variable Genes: Highly variable genes are identified using the variance-stabilizing transformation (vst) method, with the top 1,000 features selected. The function used is `FindVariableFeatures` from *Seurat*
- Scaling: All genes are scaled using the `ScaleData` function, which centers the expression values around zero and normalizes variance.
- PCA: Principal Component Analysis (PCA) is conducted using the highly variable genes and retaining the first 50 PCs. The algorithm used is IRLBA.
- t-SNE: `RunTSNE` from *Seurat* is applied on the first 50 PCs. The perplexity parameter is set to 18.
- UMAP: The `runUMAP` function of the `Seurat` package reduces dimensionality starting from the top 50 PCs using the *uwot* R package implementation, using the cosine metric.
- Neighbor graph (SNN): The `FindNeighbors` function constructs a Shared Nearest Neighbor (SNN) graph based on the top 50 PCs, with *k* set to 20.
- Louvain: The `FindClusters` function is applied to the SNN graph computed in the previous step, setting the *algorithm* parameter equal to 1 (Louvain), with a resolution of 0.2 for BE1, 0.04 for Hao, 0.1 for sc_mixology, and 0.2 for cord blood, matching the number of clusters in the ground truth.
- Leiden: The `FindClusters` function is applied to the same SNN graph, setting the *algorithm* parameter equal to 4 (Leiden), with a resolution of 0.2 for BE1, 0.08 for sc_mixology, 0.04 for Hao, and 0.2 for cord blood, matching the number of clusters in the ground truth.

#### 4.5.2 Orchestrating Single-Cell Analysis (OSCA)

Orchestrating Single-Cell Analysis (OSCA) with Bioconductor is a framework and set of tools within the Bioconductor ecosystem for analyzing scRNA-seq data. It focuses on integrating multiple R packages to provide a comprehensive workflow for processing, analyzing, and visualizing single-cell datasets. Bioconductor uses the SingleCellExper-iment class for storing single-cell assay data and metadata: count matrices are stored in the assays component, where rows represent features (e.g. genes, transcripts) and columns represent cells. In addition, low-dimensional representations of the primary data, and metadata describing cell or feature characteristics can also be stored in the SingleCellExperiment object. By standardizing the storage of single-cell data and results, Bioconductor supports interoperability between single-cell analysis packages.

Specific steps and default values of the OSCA workflow include:

- Find Mitochondrial Genes: Mitochondrial genes are identified by mapping gene symbols to their chromosomal location using the *EnsDb.Hsapiens.v75* package for human datasets and the *EnsDb.Mmusculus.v79* package for mouse datasets. Metrics such as mitochondrial percentage and total counts per cell are calculated using the `perCellQCMetrics` function from *scuttle*. Cells with high mitochondrial content (less than 5%) or low total counts are flagged and filtered out.
- Filtering: Cells are filtered based on sequencing depth and quality control metrics obtained in the previous step.
- Normalization: The `logNormCounts` function of the *scuttle* package normalizes gene expression data by calculating size factors and performing log-transformation.
- Highly Variable Genes: The `modelGeneVar` function of the *scran* Bioconductor package models gene expression variance across cells, and the `getTopHVGs` function selects the top 1,000 most variable genes.
- Scaling: In the OSCA suggestion there isn’t a specific function for running the scaling step. In this case, we set `scale = TRUE` in the `runPCA` function to compute the scaling.
- PCA: The `runPCA` function of the *scater* package reduces dimensionality using the previously identified HVGs, by default using the Exact algorithm from `runSVD` in *BioCSingular* package and computing the first 50 PCs. The `runPCA` function performs scaling as part of the PCA computation.
- t-SNE: The `runTSNE` function of the *scater* package reduces dimensionality starting from the top 50 PCs. By default, the function will set a perplexity that scales with the number of cells.
- UMAP: The `runUMAP` function of the *scater* package reduces dimensionality starting from the top 50 PCs.
- Neighbor graph (SNN): A Shared Nearest Neighbor (SNN) graph is constructed from the top 50 PCs using the makeSNNGraph function from the *bluster* package (internally invoked by clusterCells with the NNGraphParam parameter).
- Louvain: The Louvain algorithm (`cluster louvain` from *igraph*, v2.1.4) is applied to the weighted SNN graph computed in the previous step, using default parameters (modularity optimization, resolution = 1).
- Leiden: The Leiden algorithm (`cluster leiden` from *igraph*, v2.1.4) is applied to the same weighted SNN graph, using default parameters (Constant Potts Model objective, resolution = 1, two iterations).

#### 4.5.3 scrapper

The Bioconductor package *scrapper* reimplements some of the OSCA functions in C++. It provides R bindings to C++ code for the analysis of single-cell expression data, primarily utilizing various *libscran* libraries. Each function within the workflow addresses a specific step in the single-cell analysis pipeline, including tasks such as quality control, clustering, and marker detection.

Specific steps and default values of the scrapper workflow include:

- Find Mitochondrial Genes: Mitochondrial genes are identified by applying a regular expression pattern `"^mt-"` or `"^MT-"` to the row names of the count matrix. This step allowed for the selection of genes associated with mitochondrial function.
- Filtering: RNA quality control metrics are computed using the `computeRnaQcMetrics` function, which included subsets for mitochondrial genes. Suggested thresholds for filtering were determined using the `suggestRnaQcThresholds` function. The `filterRnaQcMetrics` function was then applied to retain cells that met the established quality criteria.
- Normalization: The filtered count matrix was log-normalized using size factors derived from the RNA quality metrics using `centerSizeFactors`. This log-normalization was performed using the `normalizeCounts` function, ensuring that the data were adjusted for sequencing depth and other technical variations.
- Highly Variable Genes: Gene variances were modeled using the `modelGeneVariances` function. Highly variable genes were identified using the `chooseHighlyVariableGenes` function, selecting the top 1000 genes based on their variability.
- Scaling: In this package there isn’t a specific function for running the scaling step. In this case, we set `scale = TRUE` in the `runPca` function to compute the scaling.
- PCA: PCA was conducted on the normalized data of the highly variable genes using the runPca function, which by default computes the first 25 PCs. To keep the PCA results comparable, we decided to compute the first 50 PCs.
- t-SNE: t-SNE was applied to the fist 50 PCs and a default perplexity of 30. This was executed using the runTsne function.
- UMAP: UMAP was performed on the first 50 PCs, and the default number of neighbors set to 15.
- Neighbor graph (SNN): A shared nearest neighbor (SNN) graph is constructed from the top 50 PCs using the `buildSnnGraph` function.
- Louvain: The Louvain (Multilevel) algorithm is applied to the SNN graph computed in the previous step, with a resolution of 0.18 for BE1, 0.18 for sc_mixology, 0.18 for Hao, and 0.20 for cord blood.
- Leiden: Similarly, the Leiden algorithm is applied to the same SNN graph, with a resolution of 0.18 for BE1, 0.16 for sc_mixology, 0.20 for Hao, and 0.20 for cord blood.

#### 4.5.4 Scanpy

Scanpy [18] is a scalable Python package for analyzing single-cell gene expression data, encompassing a wide array of functionalities such as preprocessing, visualization, clustering, pseudotime and trajectory inference, differential expression testing, and simulation of gene regulatory networks.

Specific steps and default values of the Scanpy workflow include:

- Find Mitochondrial Genes: This step identifies mitochondrial genes by annotating those whose names start with `"MT-"` or `"mt-"`. Quality control (QC) metrics, including the percentage of mitochondrial counts per cell, are calculated using sc.pp.calculate qc metrics
- Filtering: A filtering step is applied to remove cells and genes with undesired characteristics using `scanpy.pp.filter cells` and `scanpy.pp.filter genes` functions in *scanpy* package. For BE1 and sc_mixology dataset cells expressing fewer than 200 genes and genes present in fewer than 3 cells are excluded. Additionally, cells with an excessive number of expressed genes (*>* 5000) or a high percentage of mitochondrial counts (*>* 5%) are filtered out. For cord blood dataset cells expressing fewer than 200 genes and genes present in fewer than 3 cells are excluded. Additionally, cells with an excessive number of expressed genes (*>* 2500) or a high percentage of mitochondrial counts (*>* 5%) are filtered out.
- Normalization: `scanpy.pp.normalize total` function normalizes each cell by total counts over all genes. `scanpy.pp.log1p` perform log-transformation on the normalized gene expression.
- Highly Variable Genes: Highly variable genes are identified using `sc.pp.highly variable genes` function using Seurat method. Specific thresholds for mean expression (min mean=0.0125, max mean=3) and dispersion (min disp=0.5). Up to 1,000 highly variable genes are selected for downstream analyses.
- Scaling: All genes are scaled using `scanpy.pp.scale` function, data is scaled to center values around zero and normalize variance, with a maximum absolute value of ±10.
- PCA: Principal Component Analysis (PCA) is performed using the Arpack SVD solver and retaining the first 50 PCs. The function used is `scanpy.tl.pca`
- t-SNE: t-SNE was applied to the first 50 PCs and a default perplexity of 30. This was executed using the `scanpy.tl.tsne` function.
- UMAP: UMAP is applied to the first 50 PCs and the number of neighbors is set by default to 10. The function used is `scanpy.tl.umap`.
- Neighbor graph: The nearest-neighbor distance matrix and the corresponding neighborhood graph are computed using the scanpy.pp.neighbors function, with the local neighborhood size set to 10 on the top 50 PCs.
- Louvain: The Louvain clustering algorithm is applied to the neighborhood graph computed in the previous step using the scanpy.tl.louvain function, with a resolution of 0.13 for BE1, sc_mixology, 0.05 for Hao, and cord blood.
- Leiden: Similarly, the Leiden algorithm is applied to the same neighborhood graph using the scanpy.tl.leiden function, with a resolution of 0.13 for BE1, sc_mixology, 0.07 for Hao, and cord blood.

#### 4.5.5 rapids_singlecell

rapids_singlecell [68] is a Python-based workflow that utilizes GPU computing to enhance the analysis of single-cell data. This workflow is integrated within the *scverse* environment, and its structure is built upon *Scanpy*. By leveraging GPU capabilities through CuPy and NVIDIA’s RAPIDS framework, this approach prioritizes computational efficiency, facilitating the rapid analysis of large-scale single-cell datasets.

Specific steps and default values of the rapids singlecell workflow include:

- Find Mitochondrial Genes: Mitochondrial (MT) and ribosomal (RIBO) genes are identified and annotated based on their respective prefixes (`"MT-"` and `"RPS"`) using *rsc.pp.flag genefamily*. Quality control (QC) metrics, including percentages of MT and RIBO counts, are calculated to assess sequencing quality using *rsc.pp.calculate geneqc genemetrics* function.
- Filtering: A filtering step is applied to remove cells and genes with undesired characteristics using `rsc.pp.filter cells` and `rsc.pp.filter genes` functions in *rapids singlecell* package. For BE1, sc dataset cells expressing fewer than 200 genes and genes present in fewer than 3 cells are excluded. Additionally, cells with an excessive number of expressed genes (*>* 5000) or a high percentage of mitochondrial counts (*>* 5%) are filtered out.
- Normalization: `rsc.pp.normalize total` function normalizes each cell by total counts over all genes. rsc.pp.log1p perform log-transformation on the normalized gene expression.
- Highly Variable Genes: Highly variable genes are identified using the Seurat v3 method with n top genes=1000 and the counts layer.
- Scaling: Data is scaled to zero mean and unit variance, with a maximum absolute value of plus/minus 10 to prevent extreme values from dominating the analysis.
- PCA: PCA is performed using 50 components and the auto algorithm could be one between Exact or Jacobi.
- t-SNE: t-SNE is applied to the first 50 PCs and with a perplexity of 30. *rapids singlecell.tl.tsne* is used to perform t-SNE.
- UMAP: UMAP is applied to the first 50 PCs and the number of neighbors is set by default to 10. *rapids singlecell.tl.umap* is used to perform umap.
- Neighbor graph: The nearest-neighbor distance matrix and the corresponding neighborhood graph are computed using the `rapids singlecell.pp.neighbors` function, with the local neighborhood size set to 10 on the top 50 PCs.
- Louvain: The Louvain clustering algorithm is applied to the neighborhood graph computed in the previous step using the `rapids singlecell.tl.louvain` function, with a resolution of 0.6 for BE1, 0.1 for sc_mixology, 0.05 for Hao, and 0.1 for cord blood.
- Leiden: Similarly, the Leiden algorithm is applied to the same neighborhood graph using the `rapids singlecell.tl.leiden` function, with a resolution of 0.6 for BE1, 0.1 for sc_mixology, 0.07 for Hao, and 0.1 for cord blood.

### 4.6 Datasets

#### 4.6.1 1.3M cell dataset (1.3M)

This dataset contains the gene expression of 1.3 million brain cells isolated from E18 mice and was generated using the 10x Genomics platform for the investigation of cellular heterogeneity within the developing mouse brain at embryonic day 18 (E18) [39]. Cells from the cortex, hippocampus, and ventricular zone of two embryonic mice were dissociated and used to create 133 scRNA-Seq libraries. The samples were sequenced on 11 Illumina Hiseq 4000 flow cells, resulting in a read-depth of approximately 18,500 reads per cell. The sequencing data were processed by Cell Ranger 1.2 to generate single-cell expression profiles of 1,308,421 cells. We refer to this dataset as the 1.3M cell dataset.

This dataset has been used for benchmarking the scalability of both PCA methods and different pipelines comparing different dataset subsampling. This dataset lacks a ground truth for the cell-type annotation. Notably, both Seurat and rapids-singlecell use this dataset as a reference example in their official vignettes^1^, and we followed these guides when setting up the corresponding workflows.

The downsampling was performed at the cellular level using the sample function in R, selecting random subsets of 100k, 500k, and 1M cells without replacement. The total number of genes remained constant across all subsets.

#### 4.6.2 Cite-seq cord blood (cb)

CITE-seq (Cellular Indexing of Transcriptomes and Epitopes by sequencing) applied to cord blood samples provides a comprehensive and high-dimensional dataset that captures both transcriptomic and proteomic information at the single-cell level [45]. This innovative technique combines single-cell RNA sequencing with the measurement of surface protein markers, allowing for the simultaneous profiling of gene expression and cell surface protein expression within individual cells. CITE-seq data are a combination of two data types extracted simultaneously from the same cell. The first data type is scRNA-seq data, while the second one consists of up to a hundred antibody-derived tags (ADT). We analyze the transcriptomic measurements paired with abundance estimates for 11 surface proteins, whose levels are quantified with DNA-barcoded antibodies. The dataset contains 20,400 genes and 7,858 cells. Cell type annotation was performed by leveraging ADT surface protein marker data to manually gate and classify cells based on immunological knowledge of marker distributions. This manual gating was informed by established marker-to-cell type mappings, as described in [69]. These annotations were used to generate a reference set of labels independent of scRNA-seq-derived predictions, enabling robust benchmarking of scRNA-seq analysis methods.

#### 4.6.3 BE1

BE1 is a single-cell RNAseq benchmark dataset providing a controlled heterogeneity environment using lung cancer cell lines characterised by expressing seven different driver genes (EGFR, ALK, MET, ERBB2, KRAS, BRAF, ROS1) [44]. Specifically the following lung cancer cell lines are included:

- PC9, 4492 sequenced cells (EGFR Del19, activating mutation [70])
- A549, 6898 sequenced cells (KRAS p.G12S, growth and proliferation, [71])
- NCI-H596 (HTB178), 2965 sequenced cells (MET Del14, enhanced protection from apoptosis and cellular migration [72])
- NCI-H1395 (CRL5868), 2673 sequenced cells (BRAF p.G469A, gain of function, resistant to all tested MEK ± BRAF inhibitors, [73])
- DV90, 2998 sequenced cells (ERBB2 p.V842I, increases kinase activity, [74])
- HCC78, 2748 sequenced cells (SLC34A2-ROS1 Fusion, ROS1 inhibitors have antiproliferative effect [75])
- CCL.185.IG, 6354 sequenced cells EML4-ALK Fusion-A549 Isogenic Cell.

The experiment was done using CellPlex technology from 10x Genomics allowing multiplexing samples into a single channel and therefore removing unwanted batch effects. BE1 is composed of 36,753 genes and 29,606 cells. This dataset has been used for benchmarking the pipelines and the cell line is used as the ground truth for the cell-type annotation.

#### 4.6.4 sc_mixology

sc_mixology uses three human lung adenocarcinoma cell lines HCC827, A549, H1975, H838 and H2228, which were cultured separately and then processed in three different ways [17]. Single cells from each cell line were mixed in equal proportions, with libraries generated using three different protocols: CEL-seq2, Drop-seq (with Dolomite equipment) and 10x Chromium. In this work, we use only single cells from the mixture of five cell lines with 10x Chronium protocol (GSM3618022). sc_mixology containing 11,786 genes and 3,918 cells. This dataset has been used for benchmarking the pipelines and the cell line is used as the ground truth for the cell-type annotation.

#### 4.6.5 Whole mouse brain MERFISH (4.2M)

This dataset is part of the Allen Brain Cell Atlas and consists of spatially resolved single-cell gene expression data generated using multiplexed error-robust fluorescence in situ hybridization (MERFISH) on a single adult mouse brain, as part of a comprehensive whole-brain cell atlas [76]. A targeted panel of 1,122 genes, selected as marker genes from a companion whole-brain scRNA-seq atlas [77], was imaged across 147 coronal sections spanning one hemisphere of the brain. After image segmentation and quality control, the dataset contains gene expression profiles for approximately 4.2 million cells. Compared with the other datasets used in this work, Zhuang-ABCA-1 is characterized by a much larger number of cells but a substantially smaller and fixed gene panel (1,122 genes), making it a useful benchmark for evaluating the scalability of PCA methods on very large cell numbers with low-dimensional input data. We refer to this dataset as the 4M dataset.

#### 4.6.6 Hao dataset (Hao)

This dataset is a large-scale CITE-seq reference of human peripheral blood mononuclear cells (PBMCs), in which transcriptomic (scRNA-seq) and surface protein (antibody-derived tag, ADT) measurements were obtained from the same cells [46]. It is publicly available through the Gene Expression Omnibus (accession GSE164378) and comprises gene expression profiles for 33,538 genes across 161,764 cells profiled with the 10x Genomics 3^′^ chemistry, collected from eight volunteers at three time points (before and 3 and 7 days after administration of an HIV vaccine). Celltype annotations are provided by the original authors at three levels of granularity (`celltype.l1`, `celltype.l2` and `celltype.l3`). In this work, the `celltype.l1` annotation, comprising eight cell types, is used as ground truth for computing the Adjusted Rand Index (ARI) and evaluating clustering quality. These labels were derived through a multimodal procedure: RNA data (SCTransform-normalized and integrated across the 24 donor/time point samples via reciprocal PCA) and ADT data (228 antibodies, CLR-normalized) were each reduced by PCA and combined into a weighted nearest neighbor (WNN) graph. This graph was clustered at high resolution,and the resulting clusters were curated manually through pairwise differential expression of RNA and protein markers into 57 clusters (`celltype.l3`), which were then grouped *a posteriori* into eight broad categories: CD4^+^ T, CD8^+^ T, unconventional T, B, NK, monocytes, dendritic cells, and other. Because these reference labels incorporate protein information not available to transcriptome-only workflows, and because the original annotation itself relied on PCA of batch-corrected RNA data, agreement with them should be interpreted as concordance with an expert-curated multimodal partition rather than as an absolute measure of transcriptomic accuracy.

### 4.7 Workflow performance evaluation

#### 4.7.1 ARI

The Adjusted Rand Index (ARI) [78] is a statistical measure commonly used to assess the similarity between two clustering results. It takes into account both the agreement and disagreement between data points in the original and clustered datasets, providing a normalized index that ranges from 0 to 1. In this benchmark, we opt to employ the ARI for the comparative assessment of various pipelines comparing the known cell type annotation with that obtained after the cluster algorithms.

#### 4.7.2 Clustering parameter selection

For the four datasets with ground-truth cell-type or cell-line annotations (BE1, sc_mixology, cord blood and Hao), the resolution parameter of the Louvain and Leiden clustering algorithms was selected, for each workflow independently, so that the number of resulting clusters matched the number of ground-truth cell types (Supplementary Table S1). This choice provides a comparable basis for assessing how well each workflow’s clustering recovers the known cell populations, as measured by the ARI, without directly tuning the resolution to maximize the ARI itself. For the 1.3M cell dataset, which lacks ground-truth cell-type annotations and was used exclusively to assess scalability, the default resolution value provided by each framework was used.

#### 4.7.3 Cell type purity

Cell type purity is defined as the percentage of neighbors for each cell that belong to the same cell type. Well-separated cell types should exhibit high purity as the cells from different types do not mix. Low purity corresponds to a data representation in which cells of different types are mixed together. We computed the cell type purity using the function `neighborPurity` from the *bluster* Bioconductor package [79], using the known cell types as “clusters” and the default value of *k* = 50 nearest neighbors.

#### 4.7.4 Variability explained by cell types

To quantify the proportion of total variability explained by the cell types, we employed a strategy inspired from the *scDiagnostics* Bioconductor package [80]. Briefly, for each method and each PC, we multiplied the percentage of variance explained by each PC by the *R*^2^ index of a linear model that uses that PC as a response variable and the cell type as covariate.

### 4.8 Infrastructure

#### 4.8.1 PCA

The computational resources employed for the generation of the results presented in this research paper consisted of a dual-processor configuration featuring 2x AMD EPYC 7301 CPUs clocked at 2.7 GHz, specifically provided by DiBio UniPD (referred to as *xen6* in Table S16) and a workstation equipped with a 12th Gen Intel Core i5-12600 CPU running Ubuntu 22.04 (referred to as *medium03* in Table S16)). Additionally, substantial computational acceleration was achieved through the utilization of four NVIDIA A100 SXM4 GPUs, each equipped with 40GB of memory and integrated into the JetStream platform. These high-performance computing resources played a pivotal role in facilitating the execution of complex calculations and simulations, thereby contributing to the robustness and accuracy of the findings presented in this study.

#### 4.8.2 Pipeline Single Cell

In the pursuit of benchmarking the single-cell pipelines presented in this research paper, computational resources from the CAPRI facility at the University of Padova were leveraged. The CPU component of this infrastructure was comprised of 16 Intel(R) Xeon(R) Gold 6130 processors, each operating at a clock speed of 2.10GHz (CAPRI UniPD). Additionally, we utilized 2 NVIDIA Tesla P100 GPUs for GPU-intensive tasks, each equipped with 16GB of memory (CAPRI UniPD). These resources, available through CAPRI, played a crucial role in executing the computational workflows associated with the benchmarking analyses, ensuring the efficiency and accuracy of the results detailed in this study.

#### 4.8.3 BLAS/LAPACK version

To optimize the performance of linear algebra operations in R, the default Ubuntu BLAS/LAPACK implementation can be replaced with the OpenBLAS library, which provides substantial computational improvements. On Ubuntu systems, this configuration can be readily modified using the apt package manager. Initially, the desired OpenBLAS version must be installed using `sudo apt install libopenblas0-pthread` for the parallelized version, or `libopenblas0-serial` for the single-threaded implementation. Subsequently, the `update-alternatives` command is employed to register the new library as an available alternative, specifying a priority level (e.g., 110 for the pthread version). The system administrator can then interactively select the desired implementation through `sudo update-alternatives --config`. This procedure must be repeated for both BLAS and LAPACK libraries, replacing the `libblas.so.3` and `liblapack.so.3` files respectively. Proper configuration can be verified by executing `sessionInfo()` in R, which will display the path of the currently utilized library. This approach enables seamless transitions between different BLAS implementations without requiring R recompilation, thereby allowing researchers to select the configuration most suitable for their specific computational requirements.

#### 4.8.4 Code availability

The code to reproduce all the analysis described in this paper, including the benchmark of SVD methods and scRNA-seq workflows is available at https://github.com/billila/pca_scwf_paper. In the same repository, we provide definition files for the containers to reproduce all the results. The integration of the ARPACK algorithm into the BiocSingular package is available at https://github.com/billila/BiocSingular. The package versions and software dependencies used in this study are provided in the definition files available at https://github.com/billila/pca_scwf_paper/envs.

#### 4.8.5 Data availability

The 1.3M cell dataset is available as part of the TENxBrainData Bioconductor package at https://bioconductor.org/packages/TENxBrainData. sc_mixology dataset is available at https://ncbi.nlm.nih.gov/geo/query/acc.cgi?acc=GSE118767 and more tecnichal details at https://github.com/LuyiTian/sc_mixology. The BE1 dataset is available at https://doi.org/10.6084/m9.figshare.23939481.v1. The cord blood dataset is available as part of the *SingleCellMultiModal* package at https://bioconductor.org/packages/SingleCellMultiModal

The 4M dataset is available as part of the Allen Brain Cell Atlas, with expression matrices and metadata hosted on AWS S3 at https://allen-brain-cell-atlas.s3.us-west-2.amazonaws.com/index.html#expression_matrices/Zhuang-ABCA-1/20230830/ under a CC BY 4.0 license; further details are provided at https://alleninstitute.github.io/abc_atlas_access/descriptions/Zhuang-ABCA-1.html. The Hao dataset is available through the Gene Expression Omnibus at https://ncbi.nlm.nih.gov/geo/query/acc.cgi?acc=GSE164378.

## Acknowledgements

We thank Aaron Lun for his insightful comments and helpful discussions on computational performance and workflow optimization, which contributed to improving the clarity and accuracy of our analyses. We thank Alexandru Mahmoud for serving as the Allocation Manager for BIR190004 in Jestream2 and for handling all the paperwork related to ACCESS credits for that allocation since 2021. We also thank Mark Robinson’s lab at the University of Zurich, in particular Mark Robinson, Atreya Choudhury, Ben Carillo, and Diana Stepushkina, for their code review and help with the interpretation of some results. This work used Jetstream2 at Indiana University through allocation BIR190004 from the Advanced Cyberinfrastructure Coordination Ecosystem: Services & Support (ACCESS) program, which is supported by National Science Foundation grants #2138259, #2138286, #2138307, #2137603, and #2138296. This work also used compute infrastructure funded by the University of Padova Strategic Research Infrastructure Grant 2017: “CAPRI: Calcolo ad Alte Prestazioni per la Ricerca e l’Innovazione”, and compute infrastructure CONVECS at the University of Padova, funded by the PR Veneto FESR 2021–2027 program (Priority 1, Objective 1.1, Action 1.1.2).

## Author contributions

D.R. and I.B. conceived the study. D.R., C.R., G.S., and I.B. developed the methodological framework. D.R., C.R., and G.S. supervised the work. I.B. performed the data analysis, developed the computational workflows, and prepared the figures. D.R., I.B., and C.R. wrote the manuscript. G.S., H.P., L.W., and V.C. critically revised the manuscript and provided valuable input on results, interpretative aspects, and code implementation. All authors read and approved the final manuscript.

## Ethics, Consent to Participate, and Consent to Publish declarations

Not applicable.

## Footnotes

1 Seurat: https://satijalab.org/seurat/articles/seurat5_sketch_analysis and https://satijalab.org/seurat/articles/seurat5_bpcells_interaction_vignette; rapids-singlecell: https://rapids-singlecell.scverse.org/en/latest/notebooks/brain_1M.html.

