## Supplementary figures and tables for "Benchmarking large-scale single-cell RNA-seq analysis"

### Supplementary Material for "Benchmarking large-scale single-cell RNA-seq analysis"

October 16, 2026

#### 1 Supplementary Figures and Tables

| | Number of Genes | Number of Cells | Species | Sample | Technology | Ground Truth |
|---|---|---|---|---|---|---|
| **1.3M** | 27,998 | 1,306,127 | Mouse | Brain | 10X Genomics Chromium Protocol | No |
| **BE1** | 36,753 | 29,606 | Human | Lung Adenocarcinoma | 10X Genomics Chromium Protocol | Lung Cancer Cell Lines |
| **cb** | 20,400 | 7,858 | Human | Cord Blood Mononuclear | CITE-seq | Manual Gating Antibody Expression Reference |
| **sc_mixology** | 11,786 | 3,918 | Human | Lung Adenocarcinoma | 10X Genomics Chromium Protocol | Lung Cancer Cell Lines |
| **4.2M** | 1,122 | 4,200,000 | Mouse | Brain | MERFISH | No |
| **hao** | 33,538 | 161,764 | Human | PBMC | CITE-seq (10X Genomics Chromium Protocol) | celltype.l1 (WNN-derived) |

Table S1: Summary of datasets used in the workflow and PCA benchmarks

| Method's Name | SVD Algorithm | Mat type | Library | software | CPU/GPU | Deferred | Reference |
|---|---|---|---|---|---|---|---|
| rspectra_sparse_arpack | arpack | sparse | RSpectra | R | CPU | no | (Lehoucq et al., 1998) |
| rapids_dense_exact | exact | dense | Rapids | Python | GPU | no | (Kalman, 1996) |
| scanpy_sparse_arpack | arpack | sparse | Scanpy | Python | CPU | no | (Lehoucq et al., 1998) |
| rapids_dense_jacobi | jacobi | dense | Rapids | Python | GPU | no | (Drmač and Veselić, 2008) |
| sciktlearn_sparse_IPCA | IPCA | sparse | scikt-learn | Python | CPU | no | (Ross et al., 2008) |
| scanpy_dense_arpack | arpack | dense | scanpy | Python | CPU | no | (Lehoucq et al., 1998) |
| bioc_sparse_def_random | random | sparse | BiocSingular | R | CPU | yes | (Halko et al., 2011) |
| bioc_sparsearray_def_random | random | sparse | BiocSingular | R | CPU | yes | (Halko et al., 2011) |
| scanpy_dense_random | random | dense | Scanpy | Python | CPU | no | (Halko et al., 2011) |
| scanpy_sparse_random | random | sparse | Scanpy | Python | CPU | no | (Halko et al., 2011) |
| sciktlearn_dense_IPCA | IPCA | dense | scikt-learn | Python | CPU | no | (Ross et al., 2008) |
| rspectra_dense_arpack | arpack | dense | RSpectra | R | CPU | no | (Lehoucq et al., 1998) |
| bioc_sparse_def_irlba | irlba | sparse | BiocSingular | R | CPU | yes | (Baglama and Reichel, 2005) |
| sciktlearn_dense_exact | exact | dense | scikt-learn | Python | CPU | no | (Kalman, 1996) |
| bioc_dense_random | random | dense | BiocSingular | R | CPU | no | (Halko et al., 2011) |
| bioc_sparse_random | random | sparse | BiocSingular | R | CPU | no | (Halko et al., 2011) |
| bioc_sparsearray_random | random | sparse | BiocSingular | R | CPU | no | (Halko et al., 2011) |
| bioc_hdf5_dense_random | random | hdf5 | BiocSingular | R | CPU | no | (Halko et al., 2011) |
| bioc_sparse_irlba | irlba | sparse | BiocSingular | R | CPU | no | (Baglama and Reichel, 2005) |
| bioc_sparse_def_exact | exact | sparse | BiocSingular | R | CPU | yes | (Kalman, 1996) |
| bioc_dense_irlba | irlba | dense | BiocSingular | R | CPU | no | (Baglama and Reichel, 2005) |
| bioc_sparse_exact | exact | sparse | BiocSingular | R | CPU | no | (Kalman, 1996) |
| bioc_hdf5_dense_exact | exact | hdf5 | BiocSingular | R | CPU | no | (Kalman, 1996) |
| bioc_hdf5_dense_irlba | irlba | hdf5 | BiocSingular | R | CPU | no | (Baglama and Reichel, 2005) |
| bioc_dense_exact | exact | dense | BiocSingular | R | CPU | no | (Kalman, 1996) |
| bioc_sparsearray_exact | exact | sparse | BiocSingular | R | CPU | no | (Kalman, 1996) |
| rspectra_hdf5_dense_arpack | arpack | dense | RSpectra | R | CPU | no | (Lehoucq et al., 1998) |
| rspectra_hdf5_sparse_arpack | arpack | sparse | RSpectra | R | CPU | no | (Lehoucq et al., 1998) |
| rspectra_hdf5_sd_arpack | arpack | sparse | RSpectra | R | CPU | no | (Lehoucq et al., 1998) |
| bioc_hdf5_sd_random | random | hdf5 | BiocSingular | R | CPU | no | (Halko et al., 2011) |
| bioc_hdf5_sd_exact | exact | hdf5 | BiocSingular | R | CPU | no | (Kalman, 1996) |
| bioc_hdf5_sd_irlba | irlba | hdf5 | BiocSingular | R | CPU | no | (Baglama and Reichel, 2005) |

Table S2: Comparison of various singular value decomposition (SVD) methods applied to dense or sparse data, specifying the library, programming language, computation type (CPU/GPU), use of deferred computation, and the ranking estimated using the Plackett–Luce model. Method names follow a naming convention that encodes the library, data type, and specific SVD algorithm used.

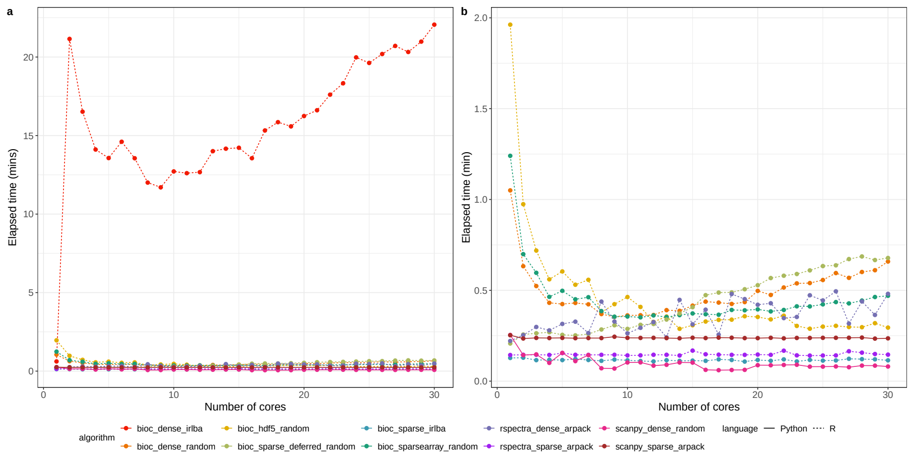


Figure S1: (a) Elapsed time for increasing number of cores for best performing method for 100k in R and Python. (b) Same as (a) without *bioc_dense_irlba*.

| Methods | n cells | Algorithms | mean_time | sd_time | Matrix type | core_eng | software |
|---|---|---|---|---|---|---|---|
| bioc_hdf5_dense | 100k | random | 108.35 | 12.91 | dense matrix | CPU | R |
| bioc_hdf5_dense | 100k | exact | 333.85 | 9.01 | dense matrix | CPU | R |
| bioc_hdf5_dense | 100k | irlba | 676.50 | 32.81 | dense matrix | CPU | R |
| rapids_dense | 100k | exact | 3.00 | 0.00 | dense matrix | GPU | python |
| rapids_dense | 100k | jacobi | 3.00 | 0.00 | dense matrix | GPU | python |
| scanpy_sparse | 100k | random | 6.20 | 0.42 | sparse matrix | CPU | python |
| scanpy_sparse | 100k | arpack | 8.10 | 0.32 | sparse matrix | CPU | python |
| sciktlearn_sparse | 100k | IPCA | 23.90 | 1.10 | sparse matrix | CPU | python |
| rspectra_dense | 100k | arpack | 33.81 | 2.64 | dense matrix | CPU | R |
| rspectra_sparse | 100k | arpack | 4.76 | 0.05 | sparse matrix | CPU | R |
| sciktlearn_dense | 100k | exact | 35.80 | 2.35 | dense matrix | CPU | python |
| bioc_dense | 100k | random | 62.59 | 0.41 | dense matrix | CPU | R |
| bioc_dense | 100k | exact | 320.42 | 5.15 | dense matrix | CPU | R |
| bioc_dense | 100k | irlba | 23.47 | 0.11 | dense matrix | CPU | R |
| bioc_sparse_def | 100k | random | 8.65 | 0.03 | sparse matrix | CPU | R |
| bioc_sparse_def | 100k | exact | 315.40 | 3.18 | sparse matrix | CPU | R |
| bioc_sparse_def | 100k | irlba | 11.59 | 0.08 | sparse matrix | CPU | R |
| bioc_sparse | 100k | random | 69.36 | 0.57 | sparse matrix | CPU | R |
| bioc_sparse | 100k | exact | 319.23 | 5.57 | sparse matrix | CPU | R |
| bioc_sparse | 100k | irlba | 11.72 | 0.08 | sparse matrix | CPU | R |
| scanpy_dense | 100k | random | 7.10 | 0.57 | dense matrix | CPU | python |
| scanpy_dense | 100k | arpack | 14.90 | 2.42 | dense matrix | CPU | python |
| sciktlearn_dense | 100k | IPCA | 37.60 | 1.51 | dense matrix | CPU | python |
| bioc_hdf5_sparse | 100k | random | 92.71 | 4.27 | sparse matrix | CPU | R |
| bioc_hdf5_sparse | 100k | exact | 80.33 | 2.37 | sparse matrix | CPU | R |
| bioc_hdf5_sparse | 100k | irlba | 1346.71 | 68.95 | sparse matrix | CPU | R |
| rspectra_hdf5_dense | 100k | arpack | 1259.35 | 0.52 | dense matrix | CPU | R |
| rspectra_hdf5_sparse | 100k | arpack | 2000.26 | 0.00 | sparse matrix | CPU | R |
| bioc_hdf5_sd | 100k | random | 1102.98 | 0.81 | sparse matrix | CPU | R |
| bioc_hdf5_sd | 100k | exact | 1359.89 | 7.74 | sparse matrix | CPU | R |
| bioc_hdf5_sd | 100k | irlba | 15703.10 | 84.08 | sparse matrix | CPU | R |
| rspectra_hdf5_sd | 100k | arpack | 529.27 | 5.72 | sparse matrix | CPU | R |

Table S3: Computational Time for PCA benchmark for all methods presented in the work. (100k)

| **Methods** | **n cells** | **algorithm** | **mean_time** | **sd_time** | **matrix type** | **core_eng** | **software** |
|---|---|---|---|---|---|---|---|
| bioc_hdf5_dense | 500k | random | 548.66 | 12.59 | dense matrix | CPU | R |
| bioc_hdf5_dense | 500k | exact | 1687.71 | 72.66 | dense matrix | CPU | R |
| bioc_hdf5_dense | 500k | irlba | 5506.23 | 214.87 | dense matrix | CPU | R |
| rapids_dense | 500k | exact | 4.10 | 0.32 | dense matrix | GPU | python |
| rapids_dense | 500k | jacobi | 4.00 | 0.00 | dense matrix | GPU | python |
| scanpy_sparse | 500k | random | 33.70 | 1.95 | sparse matrix | CPU | python |
| scanpy_sparse | 500k | arpack | 40.20 | 0.42 | sparse matrix | CPU | python |
| sciktlearn_sparse | 500k | IPCA | 117.00 | 5.73 | sparse matrix | CPU | python |
| rspectra_dense | 500k | arpack | 179.49 | 15.27 | dense matrix | CPU | R |
| rspectra_sparse | 500k | arpack | 23.63 | 0.20 | sparse matrix | CPU | R |
| sciktlearn_dense | 500k | exact | 192.00 | 8.07 | dense matrix | CPU | python |
| bioc_dense | 500k | random | 309.98 | 1.37 | dense matrix | CPU | R |
| bioc_dense | 500k | exact | 1664.83 | 9.19 | dense matrix | CPU | R |
| bioc_dense | 500k | irlba | 126.88 | 0.60 | dense matrix | CPU | R |
| bioc_sparse_def | 500k | random | 45.13 | 0.21 | sparse matrix | CPU | R |
| bioc_sparse_def | 500k | exact | 1596.32 | 8.62 | sparse matrix | CPU | R |
| bioc_sparse_def | 500k | irlba | 51.33 | 0.34 | sparse matrix | CPU | R |
| bioc_sparse | 500k | random | 395.89 | 3.38 | sparse matrix | CPU | R |
| bioc_sparse | 500k | exact | 1654.41 | 15.08 | sparse matrix | CPU | R |
| bioc_sparse | 500k | irlba | 52.10 | 0.39 | sparse matrix | CPU | R |
| scanpy_dense | 500k | random | 39.30 | 1.83 | dense matrix | CPU | python |
| scanpy_dense | 500k | arpack | 85.80 | 11.79 | dense matrix | CPU | python |
| sciktlearn_dense | 500k | IPCA | 197.80 | 1.99 | dense matrix | CPU | python |
| bioc_hdf5_sparse | 500k | random | 381.18 | 17.72 | sparse matrix | CPU | R |
| bioc_hdf5_sparse | 500k | exact | 400.42 | 12.33 | sparse matrix | CPU | R |
| bioc_hdf5_sparse | 500k | irlba | 7142.68 | 182.73 | sparse matrix | CPU | R |
| rspectra_hdf5_dense | 500k | arpack | 6213.26 | 7.51 | dense matrix | CPU | R |
| rspectra_hdf5_sparse | 500k | arpack | 10754.67 | 0.00 | sparse matrix | CPU | R |
| bioc_hdf5_sd | 500k | random | 5822.47 | 58.36 | sparse matrix | CPU | R |
| bioc_hdf5_sd | 500k | exact | 6027.23 | 8.05 | sparse matrix | CPU | R |
| bioc_hdf5_sd | 500k | irlba | 7274.23 | 85.89 | sparse matrix | CPU | R |
| rspectra_hdf5_sd | 500k | arpack | 2451.26 | 12.84 | sparse matrix | CPU | R |

Table S4: Computational Time for PCA benchmark for all methods presented in the work. (500k)

| Methods | n cells | Algorithm | mean_time | sd_time | MAtrix type | core_eng | software |
|---|---|---|---|---|---|---|---|
| bioc_hdf5_dense | 1M | random | 1147.51 | 30.52 | dense matrix | CPU | R |
| bioc_hdf5_dense | 1M | exact | 3440.95 | 174.00 | dense matrix | CPU | R |
| bioc_hdf5_dense | 1M | irlba | 12680.44 | 335.62 | dense matrix | CPU | R |
| rapids_dense | 1M | exact | 5.00 | 0.00 | dense matrix | GPU | python |
| rapids_dense | 1M | jacobi | 5.20 | 0.42 | dense matrix | GPU | python |
| scanpy_sparse | 1M | random | 69.00 | 3.74 | sparse matrix | CPU | python |
| scanpy_sparse | 1M | arpack | 82.00 | 0.47 | sparse matrix | CPU | python |
| sciktlearn_sparse | 1M | IPCA | 222.30 | 7.01 | sparse matrix | CPU | python |
| rspectra_dense | 1M | arpack | 376.11 | 31.68 | dense matrix | CPU | R |
| rspectra_sparse | 1M | arpack | 47.41 | 0.46 | sparse matrix | CPU | R |
| sciktlearn_dense | 1M | exact | 411.50 | 69.64 | dense matrix | CPU | python |
| bioc_dense | 1M | random | 631.87 | 1.97 | dense matrix | CPU | R |
| bioc_dense | 1M | exact | 3356.71 | 5.58 | dense matrix | CPU | R |
| bioc_dense | 1M | irlba | 271.36 | 1.51 | dense matrix | CPU | R |
| bioc_sparse_def | 1M | random | 94.47 | 0.35 | sparse matrix | CPU | R |
| bioc_sparse_def | 1M | exact | 3226.88 | 22.44 | sparse matrix | CPU | R |
| bioc_sparse_def | 1M | irlba | 163.69 | 0.95 | sparse matrix | CPU | R |
| bioc_sparse | 1M | random | 925.95 | 5.16 | sparse matrix | CPU | R |
| bioc_sparse | 1M | exact | 3343.25 | 22.82 | sparse matrix | CPU | R |
| bioc_sparse | 1M | irlba | 163.28 | 0.31 | sparse matrix | CPU | R |
| scanpy_dense | 1M | random | 81.70 | 2.87 | dense matrix | CPU | python |
| scanpy_dense | 1M | arpack | 205.80 | 13.26 | dense matrix | CPU | python |
| sciktlearn_dense | 1M | IPCA | 369.10 | 16.38 | dense matrix | CPU | python |
| bioc_hdf5_sparse | 1M | random | 771.28 | 25.33 | sparse matrix | CPU | R |
| bioc_hdf5_sparse | 1M | exact | 785.08 | 33.18 | sparse matrix | CPU | R |
| bioc_hdf5_sparse | 1M | irlba | 17663.29 | 961.17 | sparse matrix | CPU | R |
| rspectra_hdf5_dense | 1M | arpack | 13034.39 | 126.77 | dense matrix | CPU | R |
| rspectra_hdf5_sparse | 1M | arpack | 22256.76 | 0.00 | sparse matrix | CPU | R |
| bioc_hdf5_sd | 1M | random | 11105.98 | 0.44 | sparse matrix | CPU | R |
| bioc_hdf5_sd | 1M | exact | 1117.49 | 27.51 | sparse matrix | CPU | R |
| bioc_hdf5_sd | 1M | irlba | 1434.19 | 14.87 | sparse matrix | CPU | R |
| rspectra_hdf5_sd | 1M | arpack | 469.60 | 2.78 | sparse matrix | CPU | R |

Table S5: Computational Time for PCA benchmark for all methods presented in the work. (1M)

| Methods | n cells | Algorithm | mean_time | sd_time | Matrix type | core_eng | software |
|---|---|---|---|---|---|---|---|
| bioc_hdf5_dense | 1.3M | random | 1515.55 | 47.78 | dense matrix | CPU | R |
| bioc_hdf5_dense | 1.3M | exact | 4513.47 | 250.32 | dense matrix | CPU | R |
| bioc_hdf5_dense | 1.3M | irlba | 17995.83 | 436.47 | dense matrix | CPU | R |
| rapids_dense | 1.3M | exact | 7.50 | 0.53 | dense matrix | GPU | python |
| rapids_dense | 1.3M | jacobi | 8.00 | 0.00 | dense matrix | GPU | python |
| scanpy_sparse | 1.3M | random | 93.40 | 4.70 | sparse matrix | CPU | python |
| scanpy_sparse | 1.3M | arpack | 106.60 | 0.97 | sparse matrix | CPU | python |
| sciktlearn_sparse | 1.3M | IPCA | 292.40 | 9.88 | sparse matrix | CPU | python |
| rspectra_dense | 1.3M | arpack | 467.73 | 25.02 | dense matrix | CPU | R |
| rspectra_sparse | 1.3M | arpack | 61.96 | 0.92 | sparse matrix | CPU | R |
| sciktlearn_dense | 1.3M | exact | 546.20 | 69.81 | dense matrix | CPU | python |
| bioc_dense | 1.3M | random | 855.92 | 2.42 | dense matrix | CPU | R |
| bioc_dense | 1.3M | exact | 4423.35 | 6.07 | dense matrix | CPU | R |
| bioc_dense | 1.3M | irlba | 372.98 | 1.05 | dense matrix | CPU | R |
| bioc_sparse_def | 1.3M | random | 122.01 | 0.63 | sparse matrix | CPU | R |
| bioc_sparse_def | 1.3M | exact | 4281.73 | 22.22 | sparse matrix | CPU | R |
| bioc_sparse_def | 1.3M | irlba | 173.29 | 1.22 | sparse matrix | CPU | R |
| bioc_sparse | 1.3M | random | 1293.50 | 6.83 | sparse matrix | CPU | R |
| bioc_sparse | 1.3M | exact | 4379.43 | 22.55 | sparse matrix | CPU | R |
| bioc_sparse | 1.3M | irlba | 174.36 | 0.39 | sparse matrix | CPU | R |
| scanpy_dense | 1.3M | random | 109.40 | 4.86 | dense matrix | CPU | python |
| scanpy_dense | 1.3M | arpack | 263.70 | 16.61 | dense matrix | CPU | python |
| sciktlearn_dense | 1.3M | IPCA | 520.80 | 18.94 | dense matrix | CPU | python |
| bioc_hdf5_sparse | 1.3M | random | 811.56 | 11.06 | sparse matrix | CPU | R |
| bioc_hdf5_sparse | 1.3M | exact | 763.67 | 18.25 | sparse matrix | CPU | R |
| bioc_hdf5_sparse | 1.3M | irlba | 18659.81 | 1360.73 | sparse matrix | CPU | R |
| rspectra_hdf5_dense | 1.3M | arpack | 17218.16 | 176.35 | dense matrix | CPU | R |
| rspectra_hdf5_sparse | 1.3M | arpack | 22308.97 | 0.00 | sparse matrix | CPU | R |
| bioc_hdf5_sd | 1.3M | random | 15448.62 | 425.69 | sparse matrix | CPU | R |
| bioc_hdf5_sd | 1.3M | exact | 15703.10 | 84.07 | sparse matrix | CPU | R |
| bioc_hdf5_sd | 1.3M | irlba | 18039.03 | 26.24 | sparse matrix | CPU | R |
| rspectra_hdf5_sd | 1.3M | arpack | 5980.74 | 45.84 | sparse matrix | CPU | R |

Table S6: Computational Time for PCA benchmark for all methods presented in the work. (1.3M)

| n cells | Methods | algorithm | mean_max_mem | sd_max_mem | matrix type | core_eng | soft |
|---|---|---|---|---|---|---|---|
| 100k | bioc_hdf5 | random | 1329.894025 | 0.01 | dense matrix | CPU | R |
| 100k | bioc_hdf5 | exact | 4341.543908 | 11.6 | dense matrix | CPU | R |
| 100k | bioc_hdf5 | irlba | 49355.08551 | 0.01 | dense matrix | CPU | R |
| 100k | bioc_hdf5 | random | 1020.8476 | 0.01 | sparse matrix | CPU | R |
| 100k | bioc_hdf5 | exact | 5049.938 | 0.01 | sparse matrix | CPU | R |
| 100k | bioc_hdf5 | irlba | 26891.1733 | 0.01 | sparse matrix | CPU | R |
| 100k | bioc_dense | random | 16210.72193 | 0 | dense matrix | CPU | R |
| 100k | bioc_dense | exact | 22858.62363 | 0 | dense matrix | CPU | R |
| 100k | bioc_dense | irlba | 49560.05276 | 18.88 | dense matrix | CPU | R |
| 100k | bioc_sparse | random | 9037.863437 | 0 | sparse matrix | CPU | R |
| 100k | bioc_sparse | exact | 14623.11574 | 0 | sparse matrix | CPU | R |
| 100k | bioc_sparse | irlba | 37400.22575 | 18.88 | sparse matrix | CPU | R |
| 100k | bioc_sparse_def | random | 6310.129139 | 0 | sparse matrix | CPU | R |
| 100k | bioc_sparse_def | exact | 14843.84114 | 0 | sparse matrix | CPU | R |
| 100k | bioc_sparse_def | irlba | 38855.68576 | 12181.2 | sparse matrix | CPU | R |
| 100k | scanpy_dense | random | 1722.33429 | 0.3140134532 | dense matrix | CPU | python |
| 100k | scanpy_dense | arpack | 1652.267075 | 0.6657700317 | dense matrix | CPU | python |
| 100k | sciktlearn_dense | IPCA | 1760.367203 | 0.5872789718 | dense matrix | CPU | python |
| 100k | rapids_dense | exact | 4920 | 329.3090409 | dense matrix | GPU | python |
| 100k | rapids_dense | jacobi | 5120 | 413.1182236 | dense matrix | GPU | python |
| 100k | scanpy_sparse | random | 1750.650406 | 2.317954596 | sparse matrix | CPU | python |
| 100k | scanpy_sparse | arpack | 986.8175507 | 0.2107660095 | sparse matrix | CPU | python |
| 100k | sciktlearn_sparse | IPCA | 1116.742706 | 31.31881767 | sparse matrix | CPU | python |
| 100k | rspectra_dense | arpack | 12133.5152 | 0 | dense matrix | CPU | R |
| 100k | rspectra_sparse | arpack | 5263.775693 | 0 | sparse matrix | CPU | R |
| 100k | sciktlearn_dense | exact | 2251.295471 | 405.3892638 | dense matrix | CPU | python |
| 100k | rspectra_hdf5_dense | arpack | 943.05 | 0 | dense matrix | CPU | R |
| 100k | rspectra_hdf5_sparse | arpack | 1015.37 | 0 | sparse matrix | CPU | R |
| 100k | rspectra_hdf5_sd | arpack | 5263.77 | 0 | sparse matrix | CPU | R |
| 100k | bioc_hdf5_sd | random | 963.15 | 0.01 | dense matrix | CPU | R |
| 100k | bioc_hdf5_sd | exact | 943.05 | 11.6 | dense matrix | CPU | R |
| 100k | bioc_hdf5_sd | irlba | 1015.37 | 0.01 | dense matrix | CPU | R |

Table S7: Memory usage for PCA benchmark for all methods presented in the work. (100k)

| n cells | Methods | algorithm | mean_max_mem | sd_max_mem | matrix type | core_eng | soft |
|---|---|---|---|---|---|---|---|
| 500k | bioc_hdf5 | random | 2879.39 | 0.01 | dense matrix | CPU | R |
| 500k | bioc_hdf5 | exact | 16523.93 | 0.01 | dense matrix | CPU | R |
| 500k | bioc_hdf5 | irlba | 49259.32 | 27.27 | dense matrix | CPU | R |
| 500k | bioc_hdf5 | random | 2530.24 | 0.01 | sparse matrix | CPU | R |
| 500k | bioc_hdf5 | exact | 12821.22 | 0.01 | sparse matrix | CPU | R |
| 500k | bioc_hdf5 | irlba | 7073.86 | 0.01 | sparse matrix | CPU | R |
| 500k | bioc_dense | random | 19384.29 | 0.00 | dense matrix | CPU | R |
| 500k | bioc_dense | exact | 27037.59 | 0.00 | dense matrix | CPU | R |
| 500k | bioc_dense | irlba | 49581.63 | 0.00 | dense matrix | CPU | R |
| 500k | bioc_sparse | random | 9632.38 | 0.00 | sparse matrix | CPU | R |
| 500k | bioc_sparse | exact | 20630.80 | 0.00 | sparse matrix | CPU | R |
| 500k | bioc_sparse | irlba | 38742.98 | 0.00 | sparse matrix | CPU | R |
| 500k | bioc_sparse_def | random | 9050.79 | 0.00 | sparse matrix | CPU | R |
| 500k | bioc_sparse_def | exact | 20625.30 | 0.00 | sparse matrix | CPU | R |
| 500k | bioc_sparse_def | irlba | 40071.07 | 12567.15 | sparse matrix | CPU | R |
| 500k | scanpy_dense | random | 7735.24 | 4.25 | dense matrix | CPU | python |
| 500k | scanpy_dense | arpack | 7375.11 | 0.29 | dense matrix | CPU | python |
| 500k | sciktlearn_dense | IPCA | 7723.03 | 0.69 | dense matrix | CPU | python |
| 500k | rapids_dense | exact | 7720.00 | 3661.45 | dense matrix | GPU | python |
| 500k | rapids_dense | jacobi | 7000.00 | 573.49 | dense matrix | GPU | python |
| 500k | scanpy_sparse | random | 7925.61 | 0.38 | sparse matrix | CPU | python |
| 500k | scanpy_sparse | arpack | 4073.94 | 0.29 | sparse matrix | CPU | python |
| 500k | sciktlearn_sparse | IPCA | 4215.45 | 215.38 | sparse matrix | CPU | python |
| 500k | rspectra_dense | arpack | 15385.56 | 0.00 | dense matrix | CPU | R |
| 500k | rspectra_sparse | arpack | 5467.15 | 0.00 | sparse matrix | CPU | R |
| 500k | sciktlearn_dense | exact | 10598.58 | 2289.88 | dense matrix | CPU | python |
| 500k | rspectra_hdf5_dense | arpack | 1371.18 | 0 | dense matrix | CPU | R |
| 500k | rspectra_hdf5_sparse | arpack | 1404.82 | 0 | sparse matrix | CPU | R |
| 500k | rspectra_hdf5_sd | arpack | 1227.49 | 0 | sparse matrix | CPU | R |
| 500k | bioc_hdf5$_sd$ | random | 2933.211 | 0.01 | dense matrix | CPU | R |
| 500k | bioc_hdf5$_sd$ | exact | | 15994.353 | dense matrix | CPU | R |
| 500k | bioc_hdf5$_sd$ | irlba | 11093.391 | 0.01 | dense matrix | CPU | R |

Table S8: Memory usage for PCA benchmark for all methods presented in the work. (500k)

| n cells | Methods | Algorithm | Mean Max Mem | SD Max Mem | Matrix Type | Core Eng | Soft |
|---|---|---|---|---|---|---|---|
| 1M | bioc_hdf5 | random | 4953.075776 | 0.01 | dense matrix | CPU | R |
| 1M | bioc_hdf5 | exact | 32601.15021 | 0.01 | dense matrix | CPU | R |
| 1M | bioc_hdf5 | irlba | 37319.30883 | 240.21 | dense matrix | CPU | R |
| 1M | bioc_hdf5 | random | 4634.66163 | 0.01 | sparse matrix | CPU | R |
| 1M | bioc_hdf5 | exact | 25186.17480 | 0.01 | sparse matrix | CPU | R |
| 1M | bioc_hdf5 | irlba | 3234.25666 | 0.01 | sparse matrix | CPU | R |
| 1M | bioc_dense | random | 19458.31991 | 0 | dense matrix | CPU | R |
| 1M | bioc_dense | exact | 38645.91365 | 0 | dense matrix | CPU | R |
| 1M | bioc_dense | irlba | 42516.13618 | 0.25 | dense matrix | CPU | R |
| 1M | bioc_sparse | random | 12854.02404 | 0 | sparse matrix | CPU | R |
| 1M | bioc_sparse | exact | 32239.12414 | 0 | sparse matrix | CPU | R |
| 1M | bioc_sparse | irlba | 41376.78263 | 0.01 | sparse matrix | CPU | R |
| 1M | bioc_sparse_def | random | 10873.2858 | 0 | sparse matrix | CPU | R |
| 1M | bioc_sparse_def | exact | 29864.4428 | 0 | sparse matrix | CPU | R |
| 1M | bioc_sparse_def | irlba | 41082.69968 | 12884.5 | sparse matrix | CPU | R |
| 1M | scanpy_dense | random | 15202.94571 | 0.4429 | dense matrix | CPU | python |
| 1M | scanpy_dense | arpack | 14530.27153 | 0.5370 | dense matrix | CPU | python |
| 1M | sciktlearn_dense | IPCA | 15061.72638 | 0.6498 | dense matrix | CPU | python |
| 1M | rapids_dense | exact | 10720 | 413.1182 | dense matrix | GPU | python |
| 1M | rapids_dense | jacobi | 11840 | 1413.5849 | dense matrix | GPU | python |
| 1M | scanpy_sparse | random | 15576.70822 | 0.4238 | sparse matrix | CPU | python |
| 1M | scanpy_sparse | arpack | 7930.33981 | 0.1863 | sparse matrix | CPU | python |
| 1M | sciktlearn_sparse | IPCA | 7914.76173 | 0.3344 | sparse matrix | CPU | python |
| 1M | rspectra_dense | arpack | 19555.02417 | 0 | dense matrix | CPU | R |
| 1M | rspectra_sparse | arpack | 5872.58629 | 0 | sparse matrix | CPU | R |
| 1M | sciktlearn_dense | exact | 21041.25252 | 4612.26238 | dense matrix | CPU | python |
| 1M | rspectra_hdf5_dense | arpack | 1371.17 | 0 | dense matrix | CPU | R |
| 1M | rspectra_hdf5_sparse | arpack | 2013.85 | 0 | sparse matrix | CPU | R |
| 1M | rspectra_hdf5_sd | arpack | 948.72 | 0 | sparse matrix | CPU | R |
| 1M | bioc_hdf5$_sd$ | random | 2149.01 | 0.01 | dense matrix | CPU | R |
| 1M | bioc_hdf5$_sd$ | exact | | 13304.79 | dense matrix | CPU | R |
| 1M | bioc_hdf5$_sd$ | irlba | 9347.01 | 0.01 | dense matrix | CPU | R |

Table S9: Memory usage for PCA benchmark for all methods presented in the work. (1M)

| n cells | Methods | algorithm | mean_max_mem | sd_max_mem | matrix type | core_eng | soft |
|---|---|---|---|---|---|---|---|
| 1.3M | bioc_hdf5 | random | 7041.36567 | 0.01 | dense matrix | CPU | R |
| 1.3M | bioc_hdf5 | exact | 40995.85332 | 0.01 | dense matrix | CPU | R |
| 1.3M | bioc_hdf5 | irlba | 38906.17193 | 600.52 | dense matrix | CPU | R |
| 1.3M | bioc_hdf5 | random | 5430.44560 | 0.01 | sparse matrix | CPU | R |
| 1.3M | bioc_hdf5 | exact | 26411.33458 | 0.01 | sparse matrix | CPU | R |
| 1.3M | bioc_hdf5 | irlba | 4115.74381 | 0.01 | sparse matrix | CPU | R |
| 1.3M | bioc_dense | random | 23396.41941 | 0 | dense matrix | CPU | R |
| 1.3M | bioc_dense | exact | 48543.04005 | 0 | dense matrix | CPU | R |
| 1.3M | bioc_dense | irlba | 44921.05819 | 0 | dense matrix | CPU | R |
| 1.3M | bioc_sparse | random | 13994.26933 | 0 | sparse matrix | CPU | R |
| 1.3M | bioc_sparse | exact | 40612.50722 | 0 | sparse matrix | CPU | R |
| 1.3M | bioc_sparse | irlba | 44815.26027 | 0.25 | sparse matrix | CPU | R |
| 1.3M | bioc_sparse_def | random | 15338.3199 | 0 | sparse matrix | CPU | R |
| 1.3M | bioc_sparse_def | exact | 42124.57184 | 0 | sparse matrix | CPU | R |
| 1.3M | bioc_sparse_def | irlba | 24266.42509 | 7610.52 | sparse matrix | CPU | R |
| 1.3M | scanpy_dense | random | 19767.25197 | 0.4525564595 | dense matrix | CPU | python |
| 1.3M | scanpy_dense | arpack | 19419.6537 | 448.6105733 | dense matrix | CPU | python |
| 1.3M | sciktlearn_dense | IPCA | 19640.01923 | 0.56599918 | dense matrix | CPU | python |
| 1.3M | rapids_dense | exact | 14640 | 539.9588462 | dense matrix | GPU | python |
| 1.3M | rapids_dense | jacobi | 15480 | 1208.120671 | dense matrix | GPU | python |
| 1.3M | scanpy_sparse | random | 20260.21881 | 0.2514549124 | sparse matrix | CPU | python |
| 1.3M | scanpy_sparse | arpack | 10275.97389 | 0.2320058763 | sparse matrix | CPU | python |
| 1.3M | sciktlearn_sparse | IPCA | 10626.25923 | 113.9357804 | sparse matrix | CPU | python |
| 1.3M | rspectra_dense | arpack | 22396.61597 | 0 | dense matrix | CPU | R |
| 1.3M | rspectra_sparse | arpack | 6123.724961 | 0 | sparse matrix | CPU | R |
| 1.3M | sciktlearn_dense | exact | 29338.12294 | 12.39934045 | dense matrix | CPU | python |
| 1.3M | rspectra_hdf5_dense | arpack | 2238.89 | 0 | dense matrix | CPU | R |
| 1.3M | rspectra_hdf5_sparse | arpack | 2013.85 | 0 | sparse matrix | CPU | R |
| 1.3M | rspectra_hdf5_sd | arpack | 1984.38 | 0 | sparse matrix | CPU | R |
| 1.3M | bioc_hdf5$_sd$ | random | 5834.24 | 0.01 | dense matrix | CPU | R |
| 1.3M | bioc_hdf5$_sd$ | exact | 40601.7 | 0 | dense matrix | CPU | R |
| 1.3M | bioc_hdf5$_sd$ | irlba | 5872.65 | 0.01 | dense matrix | CPU | R |

Table S10: Memory usage for PCA benchmark for all methods presented in the work. (1.3M)

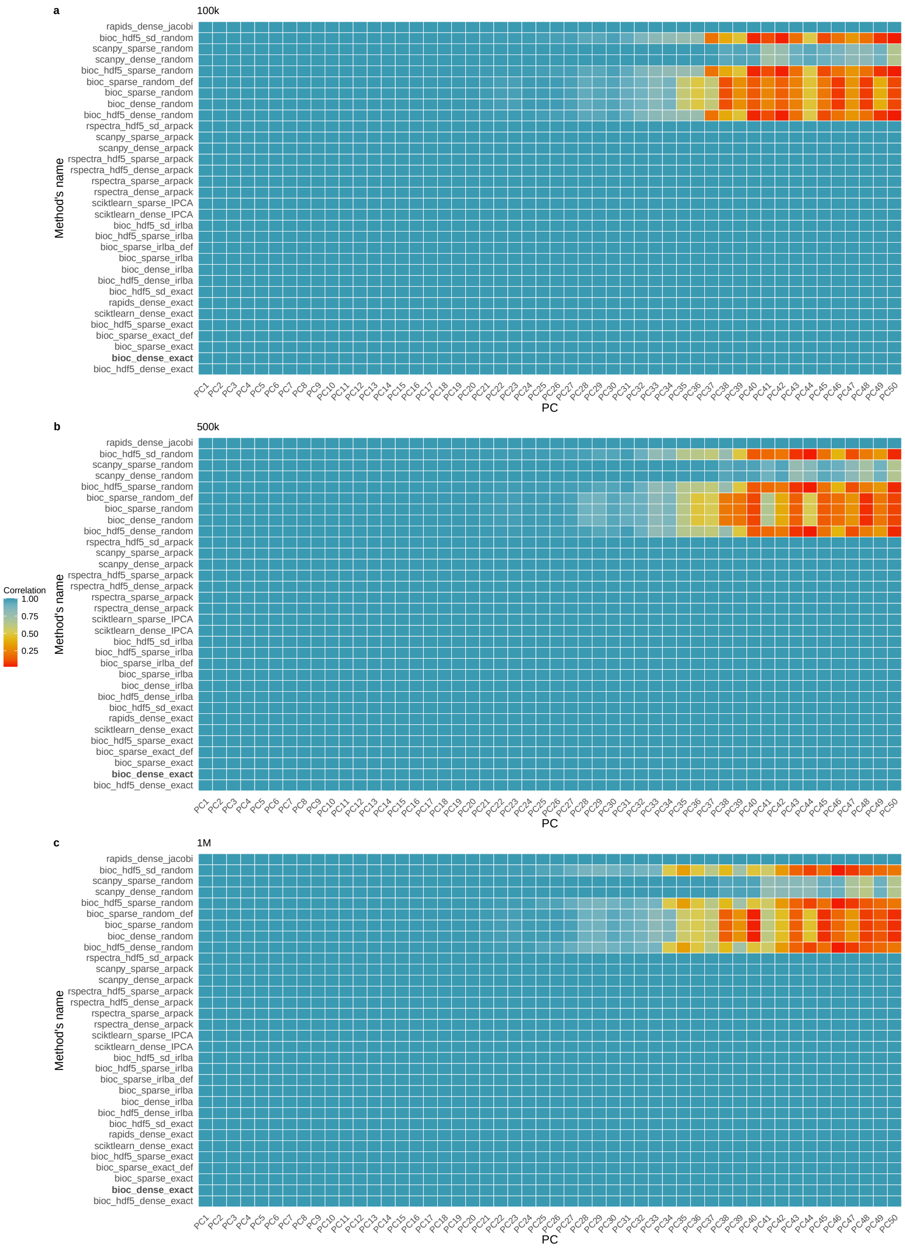


Figure S2: **Correlation heatmaps between principal components (PC)** Correlation heatmaps between principal components (PCs) obtained using different PCA algorithms and the Exact Algorithm as reference. Each row represents a different combination of PCA algorithm and input matrix, and each column a principal component (PC1–PC50). The color indicates the Pearson correlation between the PC obtained from each method and the corresponding PC from the exact reference. Panels correspond to datasets containing (a) 100k, (b) 500k, and (c) 1M cells, respectively.

| | PC1 | PC2 | PC3 | PC4 | PC5 | PC6 | PC7 | PC8 | PC9 | PC10 |
|---|---|---|---|---|---|---|---|---|---|---|
| 100k | 75.28 | 17.13 | 1.71 | 1.15 | 1.04 | 0.87 | 0.51 | 0.30 | 0.29 | 0.22 |
| 500k | 74.36 | 17.75 | 1.78 | 1.21 | 1.04 | 0.92 | 0.53 | 0.32 | 0.29 | 0.23 |
| 1M | 74.47 | 17.72 | 1.77 | 1.20 | 1.03 | 0.92 | 0.53 | 0.32 | 0.28 | 0.23 |
| 1.3M | 74.40 | 17.76 | 1.78 | 1.21 | 1.03 | 0.92 | 0.53 | 0.32 | 0.29 | 0.23 |

Table S11: Explained variance percentages for the bioc_dense_random method. Each value represents the proportion of total variance captured by the corresponding singular value component, computed using a randomized SVD algorithm on dense matrices within the BiocSingular framework. (PC1 – PC10)

| | PC11 | PC12 | PC13 | PC14 | PC15 | PC16 | PC17 | PC18 | PC19 | PC20 |
|---|---|---|---|---|---|---|---|---|---|---|
| 100k | 0.19 | 0.14 | 0.13 | 0.08 | 0.07 | 0.07 | 0.06 | 0.05 | 0.05 | 0.04 |
| 500k | 0.20 | 0.15 | 0.13 | 0.08 | 0.07 | 0.07 | 0.06 | 0.06 | 0.05 | 0.05 |
| 1M | 0.20 | 0.15 | 0.13 | 0.09 | 0.08 | 0.07 | 0.06 | 0.06 | 0.05 | 0.05 |
| 1.3M | 0.20 | 0.15 | 0.13 | 0.09 | 0.08 | 0.07 | 0.06 | 0.06 | 0.05 | 0.05 |

Table S12: Explained variance percentages for the bioc_dense_random method. Each value represents the proportion of total variance captured by the corresponding singular value component, computed using a randomized SVD algorithm on dense matrices within the BiocSingular framework. (PC11 – PC20)

| | PC21 | PC22 | PC23 | PC24 | PC25 | PC26 | PC27 | PC28 | PC29 | PC30 |
|---|---|---|---|---|---|---|---|---|---|---|
| 100k | 0.04 | 0.04 | 0.03 | 0.03 | 0.03 | 0.03 | 0.03 | 0.03 | 0.02 | 0.02 |
| 500k | 0.05 | 0.04 | 0.04 | 0.04 | 0.04 | 0.03 | 0.03 | 0.03 | 0.03 | 0.03 |
| 1M | 0.04 | 0.04 | 0.04 | 0.03 | 0.03 | 0.03 | 0.03 | 0.03 | 0.03 | 0.03 |
| 1.3M | 0.04 | 0.04 | 0.04 | 0.04 | 0.03 | 0.03 | 0.03 | 0.03 | 0.03 | 0.03 |

Table S13: Explained variance percentages for the bioc_dense_random method. Each value represents the proportion of total variance captured by the corresponding singular value component, computed using a randomized SVD algorithm on dense matrices within the BiocSingular framework. (PC21 – PC30)

| | PC31 | PC32 | PC33 | PC34 | PC35 | PC36 | PC37 | PC38 | PC39 | PC40 |
|---|---|---|---|---|---|---|---|---|---|---|
| 100k | 0.02 | 0.02 | 0.02 | 0.02 | 0.02 | 0.02 | 0.02 | 0.01 | 0.01 | 0.01 |
| 500k | 0.02 | 0.02 | 0.02 | 0.02 | 0.02 | 0.02 | 0.02 | 0.02 | 0.02 | 0.01 |
| 1M | 0.02 | 0.02 | 0.02 | 0.02 | 0.02 | 0.02 | 0.02 | 0.02 | 0.02 | 0.01 |
| 1.3M | 0.02 | 0.02 | 0.02 | 0.02 | 0.02 | 0.02 | 0.02 | 0.02 | 0.02 | 0.01 |

Table S14: Explained variance percentages for the bioc_dense_random method. Each value represents the proportion of total variance captured by the corresponding singular value component, computed using a randomized SVD algorithm on dense matrices within the BiocSingular framework. (PC31 – PC40)

| | PC41 | PC42 | PC43 | PC44 | PC45 | PC46 | PC47 | PC48 | PC49 | PC50 |
|---|---|---|---|---|---|---|---|---|---|---|
| 100k | 0.01 | 0.01 | 0.01 | 0.01 | 0.01 | 0.01 | 0.01 | 0.01 | 0.01 | 0.01 |
| 500k | 0.01 | 0.01 | 0.01 | 0.01 | 0.01 | 0.01 | 0.01 | 0.01 | 0.01 | 0.01 |
| 1M | 0.01 | 0.01 | 0.01 | 0.01 | 0.01 | 0.01 | 0.01 | 0.01 | 0.01 | 0.01 |
| 1.3M | 0.01 | 0.01 | 0.01 | 0.01 | 0.01 | 0.01 | 0.01 | 0.01 | 0.01 | 0.01 |

Table S15: Explained variance percentages for the bioc_dense_random method. Each value represents the proportion of total variance captured by the corresponding singular value component, computed using a randomized SVD algorithm on dense matrices within the BiocSingular framework. (PC41 – PC50)

| Size | Methods | Unoptimized Elapsed Time Medium03 R | Optimized Elapsed Time Medium03 R | xen6 Unoptimized | xen6 Optimized |
|---|---|---|---|---|---|
| 100k | bioc_exact_dense | 156.14 | 16.255 | 408 | 11.876 |
| 500k | bioc_exact_dense | 858.491 | 88.155 | 2088 | 76.858 |
| 1M | bioc_exact_dense | 1655.495 | 165.451 | 4392 | 151.293 |
| 1.3M | bioc_exact_dense | 2125.143 | 233.62 | 5916 | 197.526 |

Table S16: Computation times (in seconds) for exact SVD using different LAPACK/BLAS configurations in R.

| Methods | n cells | Algorithm | mean_time | sd_time | Matrix type | core_eng | software |
|---|---|---|---|---|---|---|---|
| bioc_hdf5_dense | 4.2M | random | 4101.635 | | dense matrix | CPU | R |
| bioc_hdf5_dense | 4.2M | exact | 4254.945 | | dense matrix | CPU | R |
| bioc_hdf5_dense | 4.2M | irlba | 122100.100 | | dense matrix | CPU | R |
| rapids_dense | 4.2M | exact | OOT | | dense matrix | GPU | python |
| rapids_dense | 4.2M | jacobi | OOT | | dense matrix | GPU | python |
| scanpy_sparse | 4.2M | random | 619.000 | | sparse matrix | CPU | python |
| scanpy_sparse | 4.2M | arpack | 814.000 | | sparse matrix | CPU | python |
| sciktlearn_sparse | 4.2M | IPCA | 1768.000 | | sparse matrix | CPU | python |
| rspectra_dense | 4.2M | arpack | 1017.467 | | dense matrix | CPU | R |
| rspectra_sparse | 4.2M | arpack | 368.679 | | sparse matrix | CPU | R |
| sciktlearn_dense | 4.2M | exact | Error[a] | | dense matrix | CPU | python |
| bioc_dense | 4.2M | random | 1905.905 | | dense matrix | CPU | R |
| bioc_dense | 4.2M | exact | 3579.933 | | dense matrix | CPU | R |
| bioc_dense | 4.2M | irlba | 830.935 | | dense matrix | CPU | R |
| bioc_sparse | 4.2M | random | 3905.850 | | sparse matrix | CPU | R |
| bioc_sparse | 4.2M | exact | 3389.299 | | sparse matrix | CPU | R |
| bioc_sparse | 4.2M | irlba | 365.773 | | sparse matrix | CPU | R |
| scanpy_dense | 4.2M | random | 475.000 | | dense matrix | CPU | python |
| scanpy_dense | 4.2M | arpack | 920.000 | | dense matrix | CPU | python |
| sciktlearn_dense | 4.2M | IPCA | 2456.000 | | dense matrix | CPU | python |
| bioc_hdf5_sparse | 4.2M | random | 15182.931 | | sparse matrix | CPU | R |
| bioc_hdf5_sparse | 4.2M | exact | 6495.954 | | sparse matrix | CPU | R |
| bioc_hdf5_sparse | 4.2M | irlba | OOT | | sparse matrix | CPU | R |
| rspectra_hdf5_dense | 4.2M | arpack | OOT | | dense matrix | CPU | R |
| rspectra_hdf5_sparse | 4.2M | arpack | OOT | | sparse matrix | CPU | R |
| rspectra_hdf5_sd | 4.2M | arpack | OOT | | sparse matrix | CPU | R |
| bioc_hdf5_sd | 4.2M | random | 13678.900 | | sparse matrix | CPU | R |
| bioc_hdf5_sd | 4.2M | exact | 6805.234 | | sparse matrix | CPU | R |
| bioc_hdf5_sd | 4.2M | irlba | OOT | | sparse matrix | CPU | R |

Table S17: Computational Time for PCA benchmark for all methods presented in the work. (4.2M) OOT: run exceeded the 48h wall-clock limit and was terminated. [a]scikit-learn's exact/full SVD failed on the dense representation with a LAPACK 32-bit indexing overflow (matrix size 4,676,350,140 elements).

| n cells | Methods | algorithm | mean_max_mem | sd_max_mem | matrix type | core_eng | soft |
|---|---|---|---|---|---|---|---|
| 4.2M | bioc_hdf5_dense | random | 19.01 | | dense matrix | CPU | R |
| 4.2M | bioc_hdf5_dense | exact | 118.50 | | dense matrix | CPU | R |
| 4.2M | bioc_hdf5_dense | irlba | 13.18 | | dense matrix | CPU | R |
| 4.2M | rapids_dense | exact | – | | dense matrix | GPU | python |
| 4.2M | rapids_dense | jacobi | – | | dense matrix | GPU | python |
| 4.2M | scanpy_sparse | random | 123.93 | | sparse matrix | CPU | python |
| 4.2M | scanpy_sparse | arpack | 50.50 | | sparse matrix | CPU | python |
| 4.2M | sciktlearn_sparse | IPCA | 74.75 | | sparse matrix | CPU | python |
| 4.2M | rspectra_dense | arpack | 49.24 | | dense matrix | CPU | R |
| 4.2M | rspectra_sparse | arpack | 7.18 | | sparse matrix | CPU | R |
| 4.2M | sciktlearn_dense | exact | – | | dense matrix | CPU | python |
| 4.2M | bioc_dense | random | 86.20 | | dense matrix | CPU | R |
| 4.2M | bioc_dense | exact | 165.45 | | dense matrix | CPU | R |
| 4.2M | bioc_dense | irlba | 58.74 | | dense matrix | CPU | R |
| 4.2M | bioc_sparse | random | 32.91 | | sparse matrix | CPU | R |
| 4.2M | bioc_sparse | exact | 123.40 | | sparse matrix | CPU | R |
| 4.2M | bioc_sparse | irlba | 11.11 | | sparse matrix | CPU | R |
| 4.2M | scanpy_dense | random | 83.83 | | dense matrix | CPU | python |
| 4.2M | scanpy_dense | arpack | 82.89 | | dense matrix | CPU | python |
| 4.2M | sciktlearn_dense | IPCA | 69.87 | | dense matrix | CPU | python |
| 4.2M | bioc_hdf5_sparse | random | 19.49 | | sparse matrix | CPU | R |
| 4.2M | bioc_hdf5_sparse | exact | 122.86 | | sparse matrix | CPU | R |
| 4.2M | bioc_hdf5_sparse | irlba | – | | sparse matrix | CPU | R |
| 4.2M | rspectra_hdf5_dense | arpack | – | | dense matrix | CPU | R |
| 4.2M | rspectra_hdf5_sparse | arpack | – | | sparse matrix | CPU | R |
| 4.2M | rspectra_hdf5_sd | arpack | – | | sparse matrix | CPU | R |
| 4.2M | bioc_hdf5_sd | random | 36.77 | | sparse matrix | CPU | R |
| 4.2M | bioc_hdf5_sd | exact | 119.46 | | sparse matrix | CPU | R |
| 4.2M | bioc_hdf5_sd | irlba | – | | sparse matrix | CPU | R |

Table S18: Memory usage for PCA benchmark for all methods presented in the work. (4.2M) OOT: run exceeded the 48h wall-clock limit and was terminated; entries marked with – have no associated memory measurement (OOT or failed run).

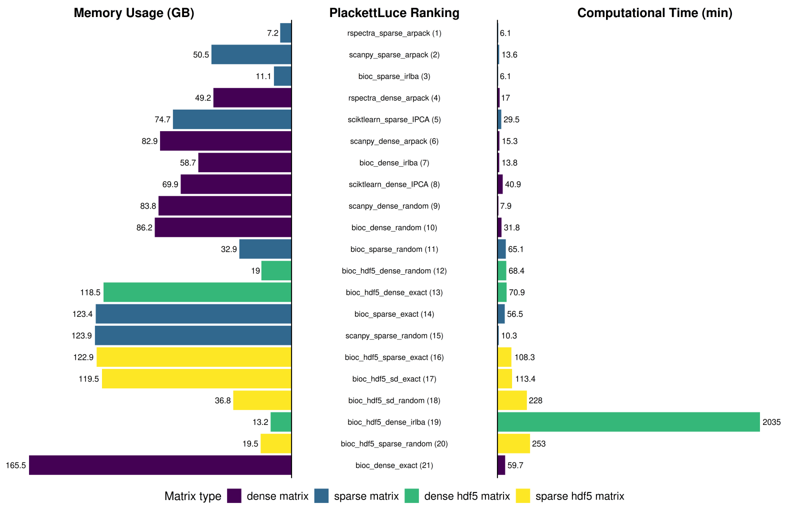


Figure S3: **Combined PlackettLuce ranking of PCA methods on the 4.2M-cell Zhuang-ABCA-1 MERFISH dataset.** Barplots show (left) peak memory usage (GB), (center) PlackettLuce ranking combining computational time, memory usage, and average correlation with the exact-SVD reference across PC1–PC50 (lower is better), and (right) computational time (minutes), for the 21 method/input combinations that completed within the 48-hour time limit. Bars are colored by matrix representation: dense matrix, sparse matrix, dense HDF5 matrix, sparse HDF5 matrix. Methods excluded from the ranking (RAPIDS exact/Jacobi, RSpectra on all three HDF5-backed representations, BiocSingular IRLBA on HDF5-backed sparse matrices, and scikit-learn's exact SVD) are listed in Supplementary Tables S17–S18.

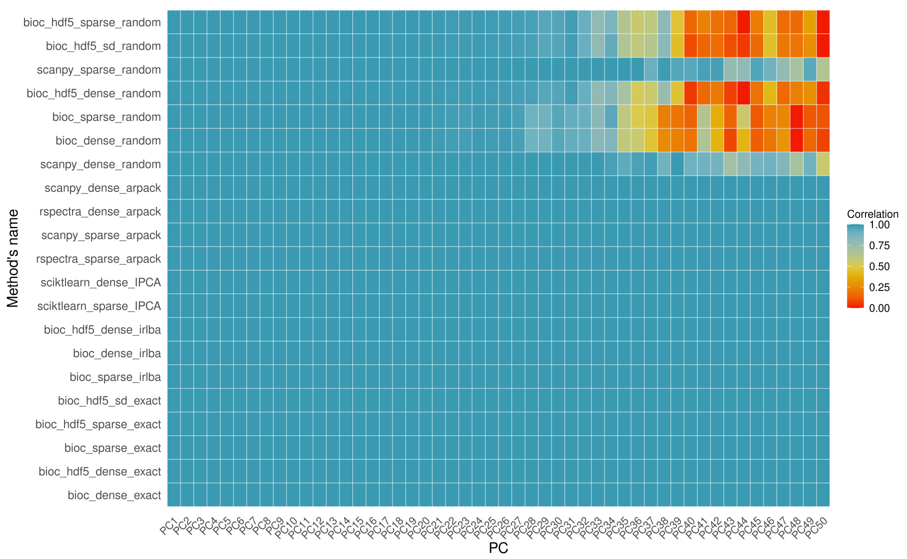


Figure S4: **Correlation heatmaps between principal components (PC) on the 4.2M-cell Zhuang-ABCA-1 MERFISH dataset.** Correlation heatmaps between principal components (PCs) obtained using different PCA algorithms and the Exact Algorithm as reference. Each row represents a different combination of PCA algorithm and input matrix, and each column a principal component (PC1–PC50). The color indicates the Pearson correlation between the PC obtained from each method and the corresponding PC from the exact reference. respectively.

| | step | Seurat | OSCA | Scanpy | Rapids | scrapper |
|---|---|---|---|---|---|---|
| 1.3M | find mitochondrial gene | 15.544 | 0.47 | 98.8152 | 4.685 | 1010.838 |
| | filter | 139.8 | 637.842 | 209.1486 | 272.8392 | 0 |
| | normalization | 62.4 | 2674.482 | 47.0995 | 60.6309 | 0.1137 |
| | hgv | 69 | 3038.392 | 55.6003 | 90.9464 | 1393.488 |
| | scaling | 67.44 | 0 | 15.2298 | 11.978 | 0 |
| | PCA | 7344 | 1350.96 | 85.1788 | 43.69 | 536.052 |
| | t-sne | 5472 | 0.00 | 4513.9915 | 101.72 | 13638.96 |
| | umap | 1180.8 | 1200.00 | 1766.2184 | 350.452 | 1581.12 |
| | snn | 356.8 | | 139.54 | 12.56 | 0.152 |
| | louvain | 2657.4 | | 572.534 | 388.746 | 1207.2 |
| | leiden | 2636.16 | 480.00 | 745.631 | 392.798 | 1139.4 |
| cb | find mitochondrial gene | 0.9312 | 8.82 | 1.479 | 0.0195 | 0.8726 |
| | filter | 1.5802 | 0 | 1.6923 | 0.5368 | 0.05 |
| | normalization | 0.9081 | 16.8966 | 1.4094 | 0.1287 | 0.0615 |
| | hgv | 1.2972 | 0.6691 | 1.2623 | 0.6082 | 1.3087 |
| | scaling | 8.7062 | 0 | 0.5298 | 0.1742 | 0 |
| | PCA | 2.8238 | 9.399 | 0.3642 | 0.0619 | 2.238 |
| | t-sne | 24.1447 | 47.7723 | 23.3397 | 0.8886 | 18.8953 |
| | umap | 17.1268 | 15.7401 | 13.0273 | 0.3203 | 12.5144 |
| | snn | 2.9737 | 4.0636 | 0.3642 | 0.0156 | 1.2244 |
| | louvain | 1.8413 | 9.5092 | 0.6226 | 0.6467 | 2.1765 |
| | leiden | 3.7774 | 4.2469 | 0.4624 | 0.1568 | 0.9744 |
| sc_mixology | find mitochondrial gene | 0.8108 | 9.6708 | 0.8919 | 0.319 | 2.1094 |
| | filter | 1.7855 | 0.0384 | 1.511 | 0.7111 | 0.1672 |
| | normalization | 1.2739 | 3.4238 | 1.4187 | 0.1414 | 0.1361 |
| | hgv | 1.8879 | 0.6615 | 0.8054 | 2.7365 | 0.6094 |
| | scaling | 5.3893 | 0 | 0.4628 | 0.2836 | 0 |
| | PCA | 0.6137 | 3.2252 | 0.2226 | 0.8274 | 0.8305 |
| | t-sne | 14.5182 | 21.5843 | 12.3548 | 0.7482 | 7.3939 |
| | umap | 9.2598 | 7.4909 | 12.3549 | 1.512 | 0.5433 |
| | snn | 1.5163 | 0.6287 | 7.8075 | 0.586 | 5.6087 |
| | louvain | 0.4124 | 1.1276 | 0.2406 | 4.8281 | 0.4511 |
| | leiden | 1.2434 | 0.3997 | 0.1536 | 0.2254 | 0.1082 |
| BE1 | find mitochondrial gene | 4.1242 | 10.5309 | 6.6525 | 0.1141 | 0.5945 |
| | filter | 5.3468 | 0.0483 | 19.8324 | 3.8953 | 0 |
| | normalization | 6.6973 | 7.0506 | 10.1501 | 0.8609 | 1.3716 |
| | hgv | 9.672 | 0.768 | 10.1801 | 4.1441 | 1.9575 |
| | scaling | 80.112 | 0 | 2.2946 | 1.1441 | 0 |
| | PCA | 9.1503 | 50.8965 | 9.5193 | 0.1374 | 10.3883 |
| | t-sne | 123.33 | 174.798 | 89.7087 | 1.8762 | 61.074 |
| | umap | 43.3076 | 25.6546 | 22.2646 | 0.5136 | 34.9697 |
| | snn | 11.2909 | 12.1387 | 44.3398 | 0.2698 | 4.6458 |
| | louvain | 11.4551 | 96.726 | 5.3419 | 1.1544 | 9.3857 |
| | leiden | 11.4578 | 34.564 | 3.2753 | 0.5563 | 3.0145 |
| hao | find mitochondrial gene | 28.4448 | 23.3307 | 13.5638 | 0.0721 | 1.4287 |
| | filter | 15.5465 | 0.0731 | 11.5095 | 1.2585 | 0.1207 |
| | normalization | 11.2094 | 25.125 | 5.885 | 0.3775 | 5.7969 |
| | hgv | 12.1218 | 3.0055 | 5.4292 | 0.5521 | 3.4317 |
| | scaling | 2.1327 | 0 | 1.5056 | 0.1253 | 0 |
| | PCA | 23.6852 | 9.5177 | 6.8241 | 0.1801 | 57.3762 |
| | t-sne | 288.984 | 36.4513 | 157.4789 | 2.1913 | 675.54 |
| | umap | 86.268 | 144.156 | 42.7829 | 1.1003 | 2.9476 |
| | snn | 24.3218 | 13.4861 | 43.7341 | 0.5623 | 32.1316 |
| | louvain | 25.3361 | 40.1359 | 9.6037 | 1.743 | 2.6137 |
| | leiden | 25.5792 | 12.2965 | 6.6037 | 0.7243 | 37.8182 |

Table S19: Computational time for each scRNA-seq workflow and each database in input

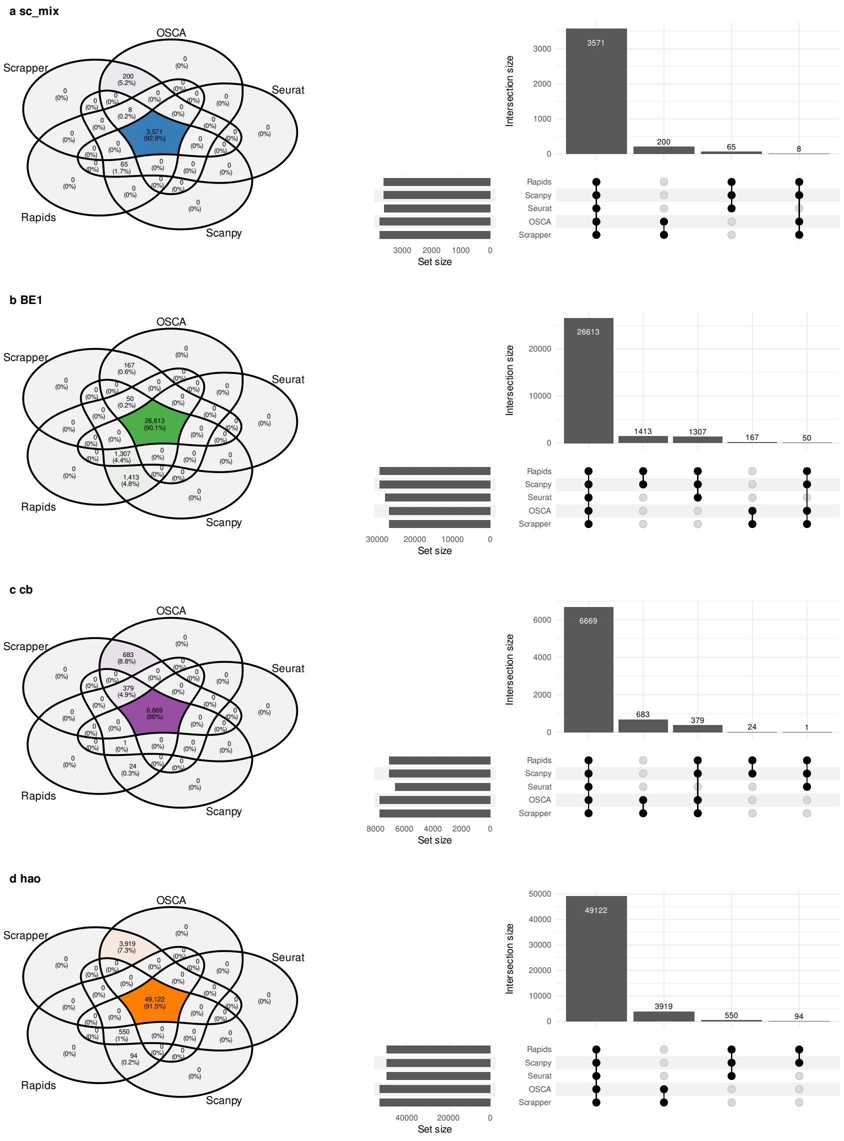


Figure S5: **Overlap of cells retained after quality control across single-cell workflows.** For each dataset with ground-truth annotations ((a) sc_mixology, (b) BE1, (c) cb, (d) hao), the Venn diagram (left) shows the number and percentage of cells retained by each combination of the five workflows (Seurat, OSCA, Scanpy, Rapids, scrapper); the highlighted central region corresponds to cells retained by all workflows. The UpSet plot (right) shows the size of the non-empty intersections (top) and the total number of cells retained by each workflow (set size, left).

| Peak Memory (GB) | Seurat | OSCA | Scanpy | Rapids | scrapper |
|---|---|---|---|---|---|
| cb | 5.86 | 7.34 | 2.30 | 2.05 | 4.93 |
| sc_mix | 3.24 | 3.67 | 2.32 | 0.93 | 2.66 |
| BE1 | 4.97 | 24.06 | 25.89 | 1.11 | 3.30 |
| hao | 40.19 | 49.36 | 16.14 | 4.53 | 7.79 |

Table S20: Peak memory usage (in GB) for each workflow and each single-cell RNA-seq dataset.

| Workflow | HVG source | Scaling | ARI | | Resolution | |
|---|---|---|---|---|---|---|
| | | | Louvain | Leiden | Louvain | Leiden |
| Seurat | own (vst) | yes (default) | 0.66 | 0.66 | 0.09 | 0.09 |
| | OSCA | yes (default) | 0.65 | 0.66 | 0.09 | 0.09 |
| | own (vst) | no | 0.67 | 0.67 | 0.05 | 0.05 |
| | OSCA | no | 0.96 | 0.96 | 0.01 | 0.01 |
| Scanpy | own (dispersion-based) | yes (default) | 0.68 | 0.68 | 0.13 | 0.13 |
| | OSCA | yes (default) | 0.65 | 0.65 | 0.13 | 0.13 |
| | own (dispersion-based) | no | 0.66 | 0.67 | 0.07 | 0.05 |
| | OSCA | no | 0.97 | 0.97 | 0.03 | 0.02 |
| OSCA | own (modelGeneVar) | no (default) | 0.97 | 0.95 | 1.0 | 1.0 |
| | own (modelGeneVar) | yes | 0.91 | 0.91 | 1.0 | 1.0 |

Table S21: Effect of highly variable gene (HVG) selection and gene-wise scaling on clustering accuracy (ARI) on the BE1 dataset. For each workflow, PCA and clustering were repeated changing one component at a time: the HVG set (the workflow's own default selection or the HVGs selected by OSCA) and whether genes were scaled to unit variance before PCA (`ScaleData(do.scale = FALSE)` in Seurat, omitting `scanpy.pp.scale` in Scanpy, `runPCA(scale = TRUE)` in OSCA), keeping all other pipeline steps unchanged. For each configuration, the clustering resolution of the Louvain and Leiden algorithms was tuned to obtain a number of clusters as close as possible to the number of cell lines.

| Workflow | HVG source | Scaling | A549 | CCL-185-IG | CRL5868 | DV90 | HCC78 | HTB178 | PBMCs | PC9 |
|---|---|---|---|---|---|---|---|---|---|---|
| Seurat | own (vst) | yes (default) | 0.9005 | 0.8964 | 0.9946 | 0.9977 | 0.9983 | 0.9968 | 0.9925 | 0.9967 |
| | OSCA | yes (default) | 0.9537 | 0.9358 | 0.9912 | 0.9947 | 0.9947 | 0.9928 | 0.9892 | 0.9948 |
| | own (vst) | no | 0.9352 | 0.9076 | 0.9979 | 0.9998 | 0.9990 | 0.9988 | 0.9969 | 0.9998 |
| | OSCA | no | 0.9989 | 0.9981 | 0.9962 | 0.9976 | 0.9960 | 0.9940 | 0.9921 | 0.9985 |
| Scanpy | own (dispersion-based) | yes (default) | 0.8206 | 0.8279 | 0.9947 | 0.9977 | 0.9986 | 0.9957 | 0.9981 | 0.9911 |
| | OSCA | yes (default) | 0.9545 | 0.9373 | 0.9886 | 0.9933 | 0.9939 | 0.9929 | 0.9893 | 0.9946 |
| | own (dispersion-based) | no | 0.8975 | 0.8603 | 0.9936 | 0.9946 | 0.9940 | 0.9945 | 0.9891 | 0.9925 |
| | OSCA | no | 0.9990 | 0.9981 | 0.9969 | 0.9976 | 0.9960 | 0.9944 | 0.9921 | 0.9987 |
| OSCA | own (modelGeneVar) | no (default) | 0.9996 | 0.9947 | 0.9966 | 0.9988 | 0.9799 | 0.9750 | 0.9567 | 0.9989 |
| | own (modelGeneVar) | yes | 0.9550 | 0.9317 | 0.9938 | 0.9981 | 0.9895 | 0.9877 | 0.9702 | 0.9976 |

Table S22: **Average cell line purity on the BE1 dataset as a function of HVG selection and scaling.** The table reports the mean neighbor purity ($k = 50$, computed on the first 50 PCs) for each cell line, for each combination of HVG set (the workflow's own default selection or the HVGs selected by OSCA) and gene-wise scaling before PCA, as in Supplementary Table S21.

| Dataset | Sample | Seurat | OSCA | Scrapper | Scanpy | rapids_singlecell |
|---|---|---|---|---|---|---|
| BE1 | A549 | 0.9002 | 0.9996 | 0.9997 | 0.8206 | 0.8206 |
| | CCL-185-IG | 0.8941 | 0.9947 | 0.9947 | 0.8279 | 0.8279 |
| | CRL5868 | 0.9925 | 0.9966 | 0.9971 | 0.9947 | 0.9947 |
| | DV90 | 0.9972 | 0.9988 | 0.9984 | 0.9977 | 0.9977 |
| | HCC78 | 0.9982 | 0.9799 | 0.9796 | 0.9986 | 0.9986 |
| | HTB178 | 0.9970 | 0.9750 | 0.9839 | 0.9957 | 0.9957 |
| | PBMCs | 0.9943 | 0.9567 | 0.9631 | 0.9981 | 0.9981 |
| | PC9 | 0.9966 | 0.9989 | 0.9993 | 0.9911 | 0.9911 |
| cb | B-cells | 1.0000 | 0.9821 | 0.9824 | 1.0000 | 1.0000 |
| | CD4 T-cells | 0.7577 | 0.5181 | 0.5285 | 0.6578 | 0.6578 |
| | CD8 T-cells | 0.7557 | 0.5418 | 0.5519 | 0.6372 | 0.6372 |
| | Monocytes CD14+ | 0.9965 | 0.9545 | 0.9517 | 0.9434 | 0.9434 |
| | Monocytes CD16+ | 0.9132 | 0.9424 | 0.9349 | 0.9802 | 0.9802 |
| | Natural Killers | 0.9107 | 0.9435 | 0.9392 | 0.9930 | 0.9930 |
| | Precursors | 1.0000 | 0.9497 | 0.9481 | 1.0000 | 1.0000 |
| | T-cells | 0.7487 | 0.6420 | 0.6162 | 0.6642 | 0.6642 |
| sc_mix | A549 | 1.0000 | 0.9994 | 0.9995 | 1.0000 | 1.0000 |
| | H1975 | 0.9535 | 0.9827 | 0.9824 | 0.9593 | 0.9593 |
| | H2228 | 0.9987 | 0.9992 | 0.9992 | 0.9987 | 0.9987 |
| | H838 | 0.9929 | 0.9976 | 0.9975 | 1.0000 | 1.0000 |
| | HCC827 | 0.9900 | 0.9966 | 0.9962 | 0.9636 | 0.9636 |
| hao | B | 1.0000 | 1.0000 | 1.0000 | 1.0000 | 1.0000 |
| | CD4 T | 0.8380 | 0.8300 | 0.8250 | 0.7310 | 0.7310 |
| | CD8 T | 0.8670 | 0.8440 | 0.8430 | 0.7450 | 0.7450 |
| | DC | 1.0000 | 0.9930 | 0.9930 | 1.0000 | 1.0000 |
| | Mono | 1.0000 | 0.9250 | 0.9240 | 1.0000 | 1.0000 |
| | NK | 0.9920 | 0.9870 | 0.9880 | 0.9940 | 0.9940 |
| | other | 0.9710 | 0.9070 | 0.9060 | 0.9670 | 0.9670 |
| | other T | 0.9060 | 0.9040 | 0.9010 | 0.8530 | 0.8530 |

Table S23: **Average cell type purity across datasets and workflows.**The table reports the mean neighbor purity values computed for each cell type across the BE1, cb, sc_mix, and hao datasets, using five single-cell analysis workflows (Seurat, OSCA, Scrapper, Scanpy, and rapidssc_singlecell).

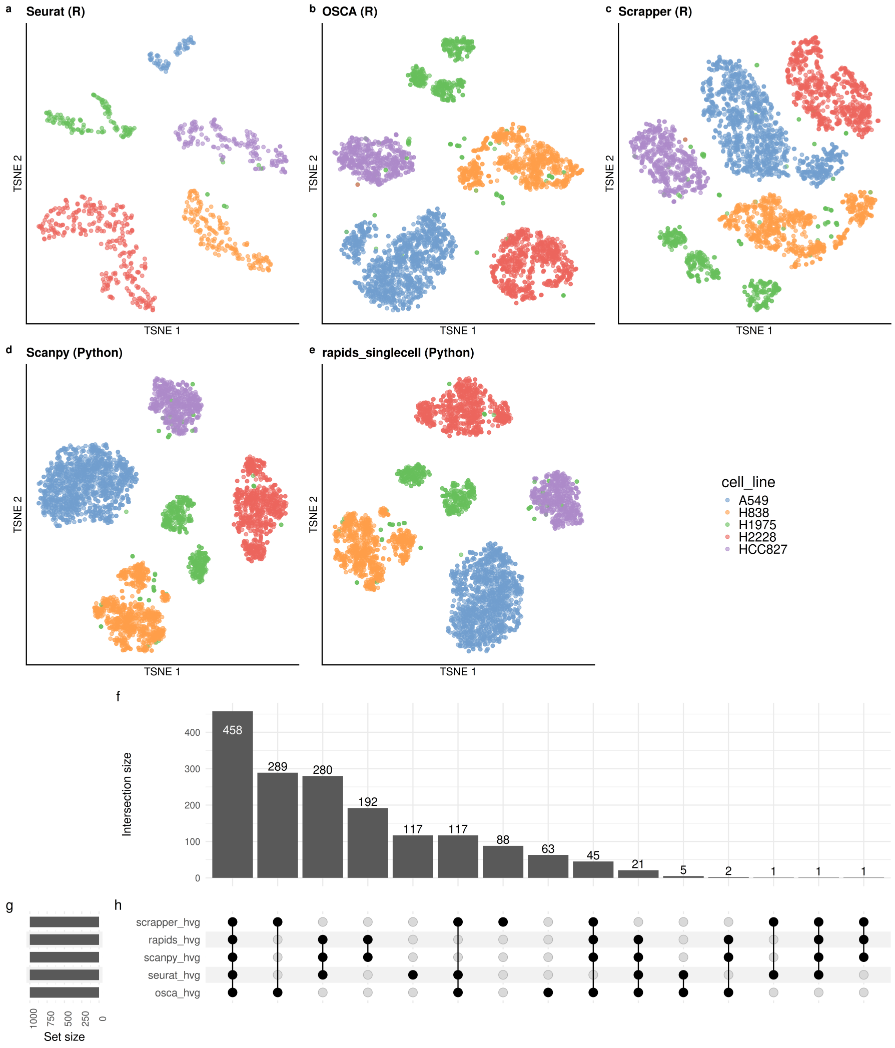


Figure S6: **Dimensionality reduction and feature selection comparison using the sc_mix dataset.** Dimensionality reduction and feature selection comparison using the sc_mix dataset. (a–e) t-SNE embeddings of the sc_mix dataset colored by sample identity, generated using five different single-cell workflows: Seurat (a), OSCA (b), Scrapper (c), Scanpy (d), and rapids_singlecell (e). Each workflow applies its own normalization and highly variable gene (HVG) selection procedure prior to dimensionality reduction. (f–h) UpSet plot showing the intersection of HVG sets selected by each workflow. Panel (f) indicates the size of each intersection set, (g) shows the number of genes selected per method (set size), and (h) depicts the overlap structure across methods.

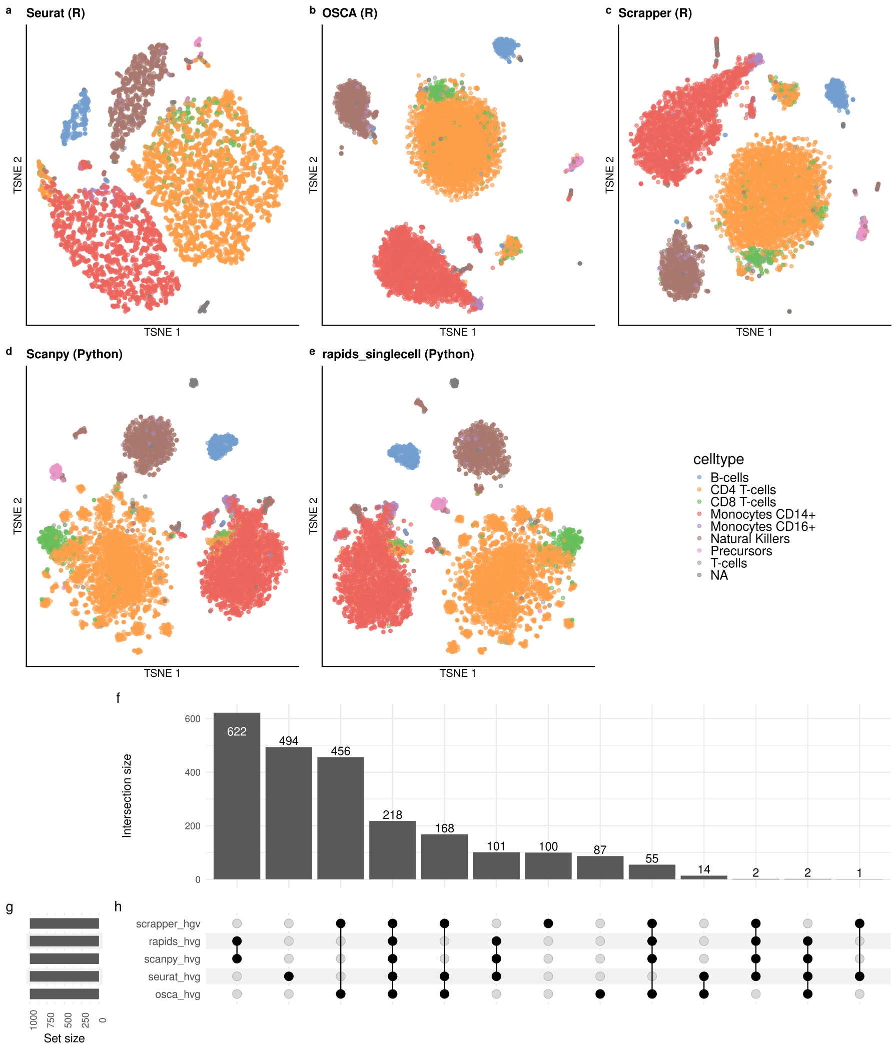


Figure S7: **Dimensionality reduction and feature selection comparison using the cb dataset.** (a–e) t-SNE embeddings of the cb dataset colored by sample identity, generated using five different single-cell workflows: Seurat (a), OSCA (b), Scrapper (c), Scanpy (d), and rapids_singlecell (e). Each workflow applies its own normalization and highly variable gene (HVG) selection procedure prior to dimensionality reduction. (f–h) UpSet plot showing the intersection of HVG sets selected by each workflow. Panel (f) indicates the size of each intersection set, (g) shows the number of genes selected per method (set size), and (h) depicts the overlap structure across methods.

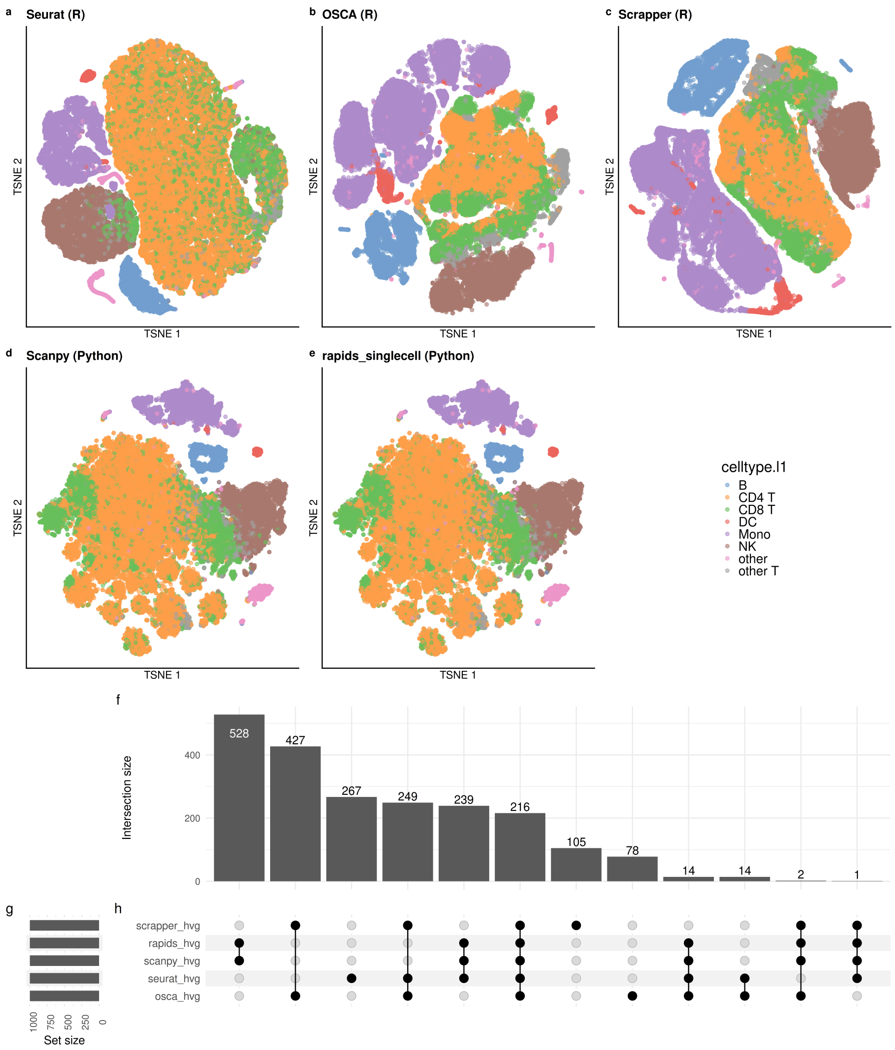


Figure S8: **Dimensionality reduction and feature selection comparison using the hao dataset.** (a–e) t-SNE embeddings of the hao dataset colored by ground-truth cell type (celltype.l1), generated using five different single-cell workflows: Seurat (a), OSCA (b), Scrapper (c), Scanpy (d), and rapids_singlecell (e). Each workflow applies its own normalization and highly variable gene (HVG) selection procedure prior to dimensionality reduction. (f–h) UpSet plot showing the intersection of HVG sets selected by each workflow. Panel (f) indicates the size of each intersection set, (g) shows the number of genes selected per method (set size), and (h) depicts the overlap structure across methods.

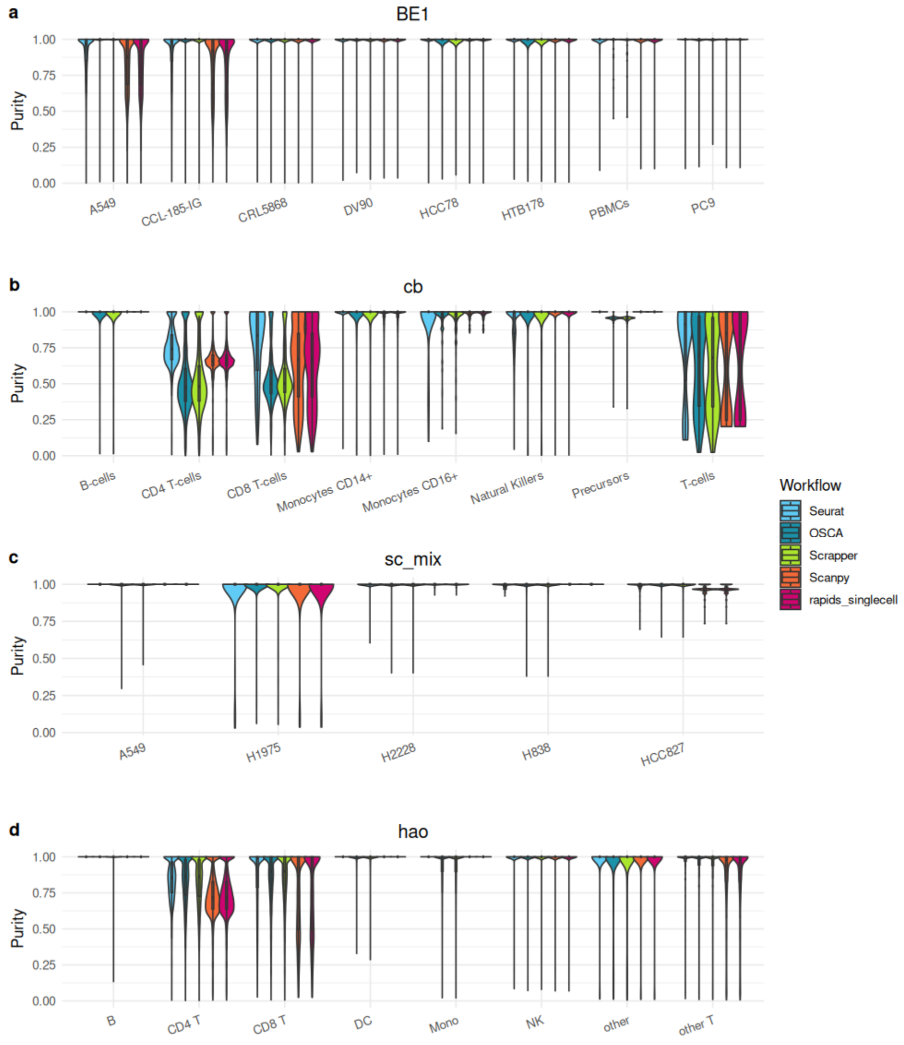


Figure S9: Comparison of clustering purity across single-cell analysis workflows on three datasets: (a) BE1, (b) cb, (c) sc_mix, (d) hao. Violin plots represent the distribution of clustering purity for each sample or cell type across five analysis workflows: Seurat, OSCA, Scrapper, Scanpy, and rapids_singlecell.

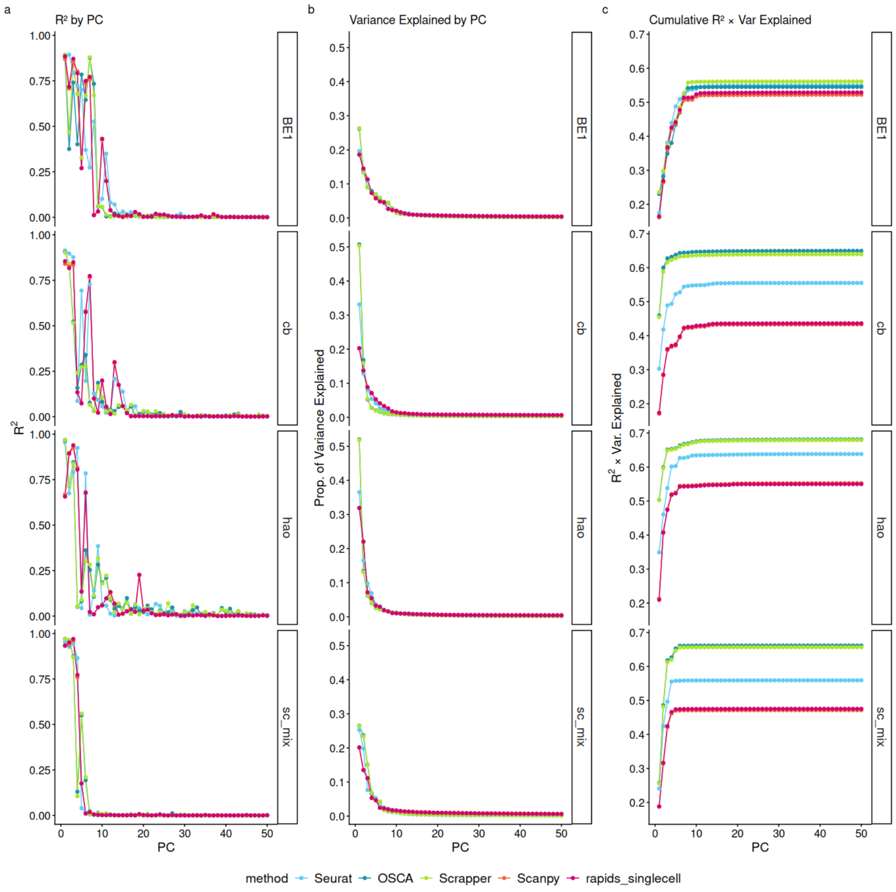


Figure S10: **Comparison of dimensionality reduction performance across single-cell workflows.** Line plots show three performance metrics computed across principal components (PCs) for three datasets: BE1, cb, sc_mix, and hao (rows) and five workflows: Seurat, OSCA, Scrapper, Scanpy, and rapids_singlecell (colors). (a) $R^2$ by PC: the proportion of variance in the cell-type labels explained by each PC. (b) Variance Explained by PC: the proportion of total variance in the gene expression data captured by each PC. (c) Cumulative $R^2$ $\times$ Variance Explained: a composite metric reflecting both biological signal and data structure captured by the PCs.

| Workflow | Default PCA/SVD algorithm | Underlying implementation | PCA benchmark method (Fig. 1c) |
|---|---|---|---|
| Seurat | IRLBA / Lanczos truncated SVD* | *irlba* R package; *BPCells* `svds` (C++ Spectra solver) for the 1.3M-cell dataset | `bioc_dense_irlba` / `rspectra_sparse`† |
| OSCA | Exact SVD / Randomized SVD* | *BiocSingular*; randomized SVD for the 1.3M-cell dataset | `bioc_dense_exact` / `bioc_sparse_exact` / `bioc_hdf5_random` |
| scrapper | IRLBA | *libscran/scran_pca* (C++), independent reimplementation | – |
| Scanpy | ARPACK | SciPy `svds` (ARPACK), sparse matrix | `scanpy_sparse_arpack` |
| rapids_singlecell | Exact / Jacobi | RAPIDS, GPU, dense matrix | `rapids_dense_exact` / `rapids_dense_jacobi` |

*For the smaller datasets, Seurat uses IRLBA, whereas OSCA uses exact SVD. For the 1.3M-cell dataset, Seurat operates on a BPCells `IterableMatrix` and uses `BPCells::svds`, based on the C++ Spectra solver, whereas OSCA uses randomized SVD through *BiocSingular* on HDF5-backed data.

†`BPCells::svds` is adapted from RSpectra and uses the same C++ Spectra solver as `RSpectra::svds`. Thus, `rspectra_sparse` represents the corresponding solver evaluated in the PCA benchmark, although the BPCells-specific implementation and matrix backend are not identical.

Table S24: Default PCA/SVD algorithms and underlying implementations used for PCA in each of the five benchmarked workflows. For Seurat and OSCA, the PCA implementation used for the 1.3M-cell dataset differs from that used for the smaller datasets because of the large-scale data backend. Where a workflow implementation corresponds to, or uses the same underlying solver as, a method evaluated in the PCA benchmark (Fig. 1c), the corresponding method name is indicated.
